# Quantitative Characterization and Prediction of the Binding Determinants and Immune Escape Hotspots for Groups of Broadly Neutralizing Antibodies Against Omicron Variants: Atomistic Modeling of the SARS-CoV-2 Spike Complexes with Antibodies

**DOI:** 10.1101/2024.12.19.629520

**Authors:** Mohammed Alshahrani, Vedant Parikh, Brandon Foley, Nishank Raisinghani, Gennady Verkhivker

## Abstract

The growing body of experimental and computational studies suggested that the cross-neutralization antibody activity against Omicron variants may be driven by balance and tradeoff of multiple energetic factors and interaction contributions of the evolving escape hotspots involved in antigenic drift and convergent evolution. However, the dynamic and energetic details quantifying the balance and contribution of these factors, particularly the balancing nature of specific interactions formed by antibodies with the epitope residues remain scarcely characterized. In this study, we performed molecular dynamics simulations, ensemble-based deep mutational scanning of SARS-CoV-2 spike residues and binding free energy computations for two distinct groups of broadly neutralizing antibodies : E1 group (BD55-3152, BD55-3546 and BD5-5840) and F3 group (BD55-3372, BD55-4637 and BD55-5514). Using these approaches, we examine the energetic determinants by which broadly potent antibodies can largely evade immune resistance. Our analysis revealed the emergence of a small number of immune escape positions for E1 group antibodies that correspond to R346 and K444 positions in which the strong van der Waals and interactions act synchronously leading to the large binding contribution. According to our results, E1 and F3 groups of Abs effectively exploit binding hotspot clusters of hydrophobic sites critical for spike functions along with selective complementary targeting of positively charged sites that are important for ACE2 binding. Together with targeting conserved epitopes, these groups of antibodies can lead to the expanded neutralization breadth and resilience to antigenic shift associated with viral evolution. The results of this study and the energetic analysis demonstrate excellent qualitative agreement between the predicted binding hotspots and critical mutations with respect to the latest experiments on average antibody escape scores. We argue that E1 and F3 groups of antibodies targeting binding epitopes may leverage strong hydrophobic interactions with the binding epitope hotspots critical for the spike stability and ACE2 binding, while escape mutations tend to emerge in sites associated with synergistically strong hydrophobic and electrostatic interactions.

## Introduction

Extensive structural and biochemical studies of the Spike (S) glycoprotein of SARS-CoV-2 have provided significant insights into the mechanisms that regulate virus transmission and immune evasion [1–9]. The S glycoprotein exhibits remarkable conformational flexibility, particularly within the S1 subunit. This subunit contains several critical domains: the N-terminal domain (NTD), the receptor-binding domain (RBD), and two structurally conserved subdomains, SD1 and SD2. This flexibility allows the S glycoprotein to adapt to various stages of the viral entry process [1–9]. The NTD is involved in initial host cell attachment, while the RBD is crucial for binding to the host cell receptor, angiotensin-converting enzyme 2 (ACE2). The SD1 and SD2 subdomains play roles in stabilizing the prefusion conformation of the S glycoprotein and facilitating the transition to the postfusion state, which is necessary for membrane fusion and viral entry. The transitions between closed and open states of the S protein, driven by conformational changes in the NTD and RBD enable the virus to effectively engage with host cell receptors while evading immune surveillance through structural variability [10–15]. Hence, the dynamic nature of the S protein enables SARS-CoV-2 to efficiently infect host cells and evade the immune system, contributing to its high transmissibility and pathogenicity. Biophysical investigations have elucidated how thermodynamic and kinetic factors govern the mechanisms of the S protein [16–18]. These studies have shown that mutations within the S protein, particularly in the S1 subunit, can induce structural alterations that affect its stability and conformational dynamics. These changes are crucial as they influence the protein’s ability to switch between open and closed states, thereby affecting the accessibility of the RBD, which is critical for viral attachment to host cells. Furthermore, the structural variability introduced by these mutations can enhance the virus’s ability to evade the immune system, making it more challenging for the host to mount an effective immune response. Understanding these mechanisms is vital for developing therapeutic strategies and vaccines aimed at targeting the S glycoprotein and preventing viral entry into host cells. The extensive collection of cryo-electron microscopy (cryo-EM) and X-ray structures of SARS-CoV-2 S protein variants of concern (VOCs) in various functional states, along with their interactions with antibodies (Abs) have shown that VOCs can induce structural changes in the dynamic equilibrium of the S protein [19–25]. These structural alterations influence the distribution of functional states and can affect the binding affinities of the S proteins with different classes of Abs and determine how effectively these Abs can neutralize the virus.

The SARS-CoV-2 XBB.1.5 subvariant emerged through recombination events within the BA.2 lineage, leading to enhanced growth and increased transmissibility which is due to its resistance to neutralization and improved binding affinity for ACE2 receptors [26–28]. These new variants are more infectious and transmissible than earlier Omicron variants. Several residues in the receptor-binding domain (RBD) are mutated in at least five new Omicron lineages. The XBB descendants EG.5 and EG.5.1 have an additional F456L mutation [29–32]. XBB subvariants with both L455F and F456L mutations are called “FLip” variants and include JG.3 (XBB.1.9.2.5.1.3.3), JF.1 (XBB.1.16.6.1), GK.3 (XBB.1.5.70.3), and JD.1.1, all of which emerged convergently. This convergence underscores that acquiring the L455F/F456L double mutation can provide a growth advantage to XBB within the human population [33].

The Omicron subvariant BA.2.86, originating from the BA.2 variant, shows substantial genetic divergence from earlier forms [34–38] and exhibits greater evasion against RBD-targeted Abs compared to the immune evasion observed in XBB.1.5 and EG.5.1 variants [34]. JN.1 is a variant of BA.2.86 that emerged independently from Omicron BA.2 and carries an additional L455S mutation, which enhances its ability to evade the immune system [39,40]. Comparative biochemical analysis using surface plasmon resonance (SPR) assays revealed a significant reduction in ACE2 binding affinity for JN.1 and the increased evasion against RBD class-1 Abs such as S2K146 and Omi-18, as well as the class-3 Abs S309 [34,39].A series of SARS-CoV-2 variants with mutations at the L455, F456, and R346 positions include the “SLip” variant, which carries the JN.1 mutation (L455S) along with an additional F456L mutation. More recently, the “FLiRT” variant has appeared, featuring an additional R346T mutation on the SLip backbone. Studies have shown that the F456L (SLip) and R346T (FLiRT) subvariants of JN.1 enhance the escape of JN.1-derived variants from neutralizing Abs [40–43]. JN.1 subvariants KP.2 (JN.1 + S:R346T, S:F456L, S:V1104L) and KP.3 (JN.1 + S:F456L, S:Q493E, S:V1104L) have convergently acquired S protein substitutions such as R346T, F456L, Q493E, and V1104L. JN.1 descendant, termed “FLuQE” (KP.3) features mutations R346T, L455S, F456L, Q493E, and V1104L and shows strong growth [44]. JN.1 subvariants LB.1 (JN.1 + S:S31-, S:Q183H, S:R346T, S:F456L), and KP.2.3 (JN.1+ S:R346T, S:H146Q, S:S31-) which convergently acquired S31 deletion in addition to the above substitutions, have spread as of June 2024 and contribute to immune evasion and the increased relative effective reproduction number [45]. KP.2 and KP.3 variants share the F456L mutation that is critical for enhancing antibody (Ab) evasion with KP.3 emerging as the most immune evasive variant and is also the fastest-growing JN.1 sublineage [46]. The epistatic impacts of mutations Q493E and F456L on the binding affinity of JN.1, KP.2 and KP.3 RBD with ACE2 were recently discovered owing to cryo-EM structures of JN.1+Q493E, KP.2 and KP.3 RBD and their complexes with ACE2 revealing that the F456L mutation can enhance binding potential of Q493E [47]. This leads to stronger receptor interactions and creates evolutionary advantage for incorporating additional evasive mutations [47]. XEC is a very recently emerged recombinant variant of KS.1.1/KP.3.3 and it carries two additional S mutations, F59S and T22N on the NTD showing the increasing potential to become the next dominant strain [49,50]. Functional assay studies showed that the infectivity of XEC was significantly higher than that of KP.3, and XEC exhibited more robust immune resistance [51,52].

High-throughput yeast display screening and deep mutational scanning (DMS) have mapped the escape mutation profiles of the RBD residues for human anti-RBD neutralizing Abs, revealing functionally significant classification of neutralizing Abs into six epitope groups (A– F) [53]. The recent seminal study exploited this approach to characterize the epitope distribution of Abs elicited by post-vaccination BA.1 infection and identified the mutational escape profiles for 1,640 RBD-binding Abs, which were classified into 12 epitope groups [54]. In this classification, groups A–C consist of Abs targeting the ACE2-binding motif. Group D Abs, such as REGN-10987, LY-CoV1404, and COV2-2130, bind to the epitope 440–449 on the RBD and are further divided into D1 and D2 subgroups. Groups E and F are subdivided into E1–E3 and F1–F3, respectively, covering the front and back of the RBD. These groups correspond to class 3 and class 4 Abs according to the earlier classification [55]. Group E Abs are sensitive to mutations of G339, T345, and R346, while neutralization by group F Abs can be compromised by changes in F374, T376, and K378 and in some Abs in this group to V503 and G504, similar to the epitopes of S2X259 suggesting that they can compete with ACE2 [53,54].

The molecular mechanisms underlying the broadly neutralizing Abs induced by XBB/JN.1 infections was conducted with high-throughput yeast-display-based DMS assays and the escape mutation profiles of a total of 2,688 Abs, including 1,874 isolated from XBB/JN.1 infection cohorts were determined producing a total of 22 Ab clusters [56]. This study showed that JN.1 reinfections elicit better Abs against emerging variants due to the enrichment of class 1 Abs and potential of F3 Omicron-specific Abs [56]. A seminal study by Cao and colleagues identified four E1 group Abs BD55-3546, BD55-3152, BD55-5585, BD55-5549 and BD55-5840 (SA58) Abs a well as F3 Abs BD55-4637, BD55-3372, BD55-5483, and BD55-5514 (SA55) [57]. This study discovered the non-competing RBD binding of SA55 and SA58 by SPR competition assays and showed that while accumulation of mutations on R346 and K444 may affect the neutralization of SA58, SA55 could efficiently neutralize all escaping mutants, including convergent mutations on RBD of BQ.1, BQ.1.1, and XBB [57]. Together, the results from recent studies revealed the evolving Ab response to Omicron antigenic shift from XBB to JN.1 where groups F3, A1, B, and D3 Abs retain potency and neutralizing activity against JN.1 subvariants, whereas A2, D2, D4, and E1/E2.1 largely escaped [46, 54, 57]. The latest studies showed that repeated Omicron infection stimulates a higher level of Omicron-specific Abs that have distinct RBD epitopes and escaping mutations as compared to WT-induced monoclonal Abs [46,58].

The two recently discovered Abs CYFN1006-1 and CYFN1006-2 demonstrated consistent neutralization of all tested SARS-CoV-2 variants, outperforming SA55 [59]. These Abs have binding epitopes overlapping with LY-CoV1404, REGN10987, and S309, located on the outer surface of the RBD. They bind to a different RBD region compared to SA55, suggesting that combining SA55 and CYFN1006-1 could be beneficial against JN.1, KP.2, KP.3, and evolving SARS-CoV-2 mutants [59]. A yeast-display system combined with a machine learning (ML)-guided approach for library design enabled an investigation of a larger number of Ab variants and the identification of a class 1 human Ab designated as VIR-7229, which targets the receptor-binding motif (RBM) potently neutralizing SARS-CoV-2 variants, including EG.5, BA.2.86, and JN.1 [60]. The structures of VIR-7229-bound to XBB.1.5 and EG.5 structures showed that the VIR-7229 interactions can accommodate both F456 and L456 in the corresponding genetic backgrounds and tolerate an extraordinary epitope variability exhibiting high barrier for the emergence of resistance, partly attributed to its high binding affinity [60]. High-throughput DMS assays were used to analyze 1,637 potent Abs against eight major SARS-CoV-2 variants, including B.1 (D614G), Omicron BA.1, BA.2, BA.5, BQ.1.1, XBB.1.5, HK.3, and JN.1 [61]. The study found that 296 Abs effectively neutralized XBB.1.5, and 147 neutralized JN.1. One notable discovery was BD55-1205, a class 1/group A1 Ab that showed broad neutralization with high affinity ranging from 1 pM to 18 nM against all major SARS-CoV-2 variants tested, including XBB, BA.2.86, and JN.1-derived subvariants, and a high barrier to escape [61].

Collectively, the growing number of structural, functional and biophysical studies revealed the diversity of mechanistic scenarios underlying Ab binding and catalogued the RBD escape mutations for a wide range of Abs unveiling the distinct signatures of Ab-resistant mutational hotspots. The recent investigations of the emerging new variants indicated that SARS-CoV-2 is more likely to evolve RBD mutations on the frequently-targeted by Abs “hotspot” epitopes while the virus is less likely to evolve mutations that could disrupt folding and stability of the RBD. The recently emerged broadly neutralizing Abs from groups E1 and F3 appeared to target unique RBD epitopes that encompass the RBD stability regions associated with critical spike function and RBM residues that are not essential for ACE2 binding affinity. Combined, these factors may enable broad immunity escape making these Abs (such as SA55, SA58 and their combinations) effective for existing VOCs and currently circulating COVID-19 Variants of Interest (VOIs).

Computer simulations have significantly advanced our understanding of the dynamics and functions of the S protein and S complexes with ACE2 and Abs at the atomic level. Our recent studies demonstrated that convergent Omicron mutations such as G446S, F486V, F486P, F486S, and F490S can display epistatic couplings with the major stability and binding affinity hotspots which may allow for the observed broad Ab resistance [62]. Molecular dynamics (MD) simulations and Markov state models (MSM) have systematically characterized the conformational landscapes of XBB.1 and XBB.1.5 Omicron variants and their complexes [63]. Mutational scanning and binding analysis of the Omicron XBB spike variants with ACE2 and a panel of class 1 Abs provided a quantitative rationale for the experimental evidence [64,65]. Epistatic interactions of physically proximal binding hotspots, such as Y501, R498, Q493, L455F, and F456L, were found to determine strong ACE2 binding, while convergent mutation sites F456L and F486P were instrumental in mediating broad Ab resistance [64,65]. We combined AlphaFold2-based atomistic predictions of structures and conformational ensembles of the S complexes with the ACE2 for the most dominant Omicron variants JN.1, KP.1, KP.2 and KP.3 to examine the mechanisms underlying the role of convergent evolution hotspots in balancing ACE2 binding and Ab evasion [66]. This study identified binding energy hotspots and characterized epistatic interactions between convergent mutational sites at L455, F456, Q493 positions that can protect and restore ACE2 binding affinity while conferring beneficial immune escape. Our previous studies revealed that the S protein can often perform its functions as an allosteric regulatory machinery that leverages the intrinsic plasticity of functional regions controlled by stable allosteric hotspots to modulate the regulatory and binding functions [67–71]. We combined all-atom MD simulations, the ensemble-based mutational scanning of binding, and perturbation-based network profiling of allosteric interactions in the S complexes with a panel of cross-reactive and ultra-potent single Abs and Ab combinations [72]. Using this approach, we suggested a mechanism in which the pattern of specific escape mutants for ultrapotent Abs may not be solely determined by the binding interaction changes but are driven by a complex balance between the impact of mutations on structural stability, binding strength, and long-range communications [72].

Convergent evolution has led to the observation of different lineages acquiring an additional group of mutations at different amino acid residues, namely R346, K444, N450, N460, F486, F490, Q493, and S494. The convergent evolution of Omicron sublineages reflected the selective pressure by previous infection- or vaccine-elicited immunity and appeared to exploit charged interactions mediated by R346, N440K, K444 hotspots to balance ACE2 binding and immune escape [73]. Many studies noted the role of electrostatic interactions as an important thermodynamic force of the S-protein binding with the ACE2 receptor and resistance to Abs [74–76]. The increased accumulation of positive charges on the RBD for many Omicron variants reflected the evolutionary changes in which the S protein has a positive electrostatic surface that promotes ACE2 binding while the electrostatic properties of the Omicron RBD often mediate immune resistance as most Abs have positively charged S RBD-recognition surfaces, allowing recent variants to escape Ab surveillance [74–76]. The large scale of electrostatic changes were observed in the BA.2.86 and JN.1 RBD indicating that electrostatic changes in BA.2.86 contribute to immune evasion [77]. The emergence of the JN.1 descendants accompanied by the increased Ab evasion and decrease in affinity to ACE2 underscores the evolutionary trajectory of SARS-CoV-2 [78]. In the JN.1 variant, there is significant accumulation of lysine residues and charged RBD residues including R403K, N460K, N481K, A484K, F486P, R493Q, E554K that modulate ACE2 binding and immune resistance. Many studies suggested functionally balanced substitutions that optimize tradeoffs between immune evasion, high ACE2 affinity and sufficient conformational adaptability might be a common strategy of the virus evolution and serve as a primary driving force behind the emergence of new Omicron subvariants [78–80]. Our computational studies of the Omicron variants BA.2, BA.3, BA.4/5, BQ.1.1, XBB.1, XBB.1.5, XBB.1.5+L455F and XBB.1.5+F456L RBD binding with ACE2 showed the favorable binding contributions provided by RBD residues K378, R403, K424, K440, K444, K460, N477, K478 that are determined by strong electrostatic interactions mediated by lysine residues, which is the result of Omicron evolution leading to significant accumulation of positively charged substitutions interacting with the negatively charged ACE2 binding interface [81,82].

Together, experimental and computational studies suggested that the cross-neutralization Ab activity against Omicron variants may be driven by balance and tradeoff of multiple energetic factors and interaction contributions of the evolving escape hotspots involved in antigenic drift and convergent evolution. However, the dynamic and energetic details quantifying the balance and contribution of these factors, particularly the balancing nature of specific interactions formed by Abs with the epitope hotspot residues remains scarcely characterized. In this study, we focused on two distinct groups of broadly neutralizing Abs : E1 group Abs (BD55-3152, BD55-3546 and SA58) and F3 group Abs (BD55-3372, BD55-4637 and SA55) (Figure 1, Table 1).

**Figure 1.**
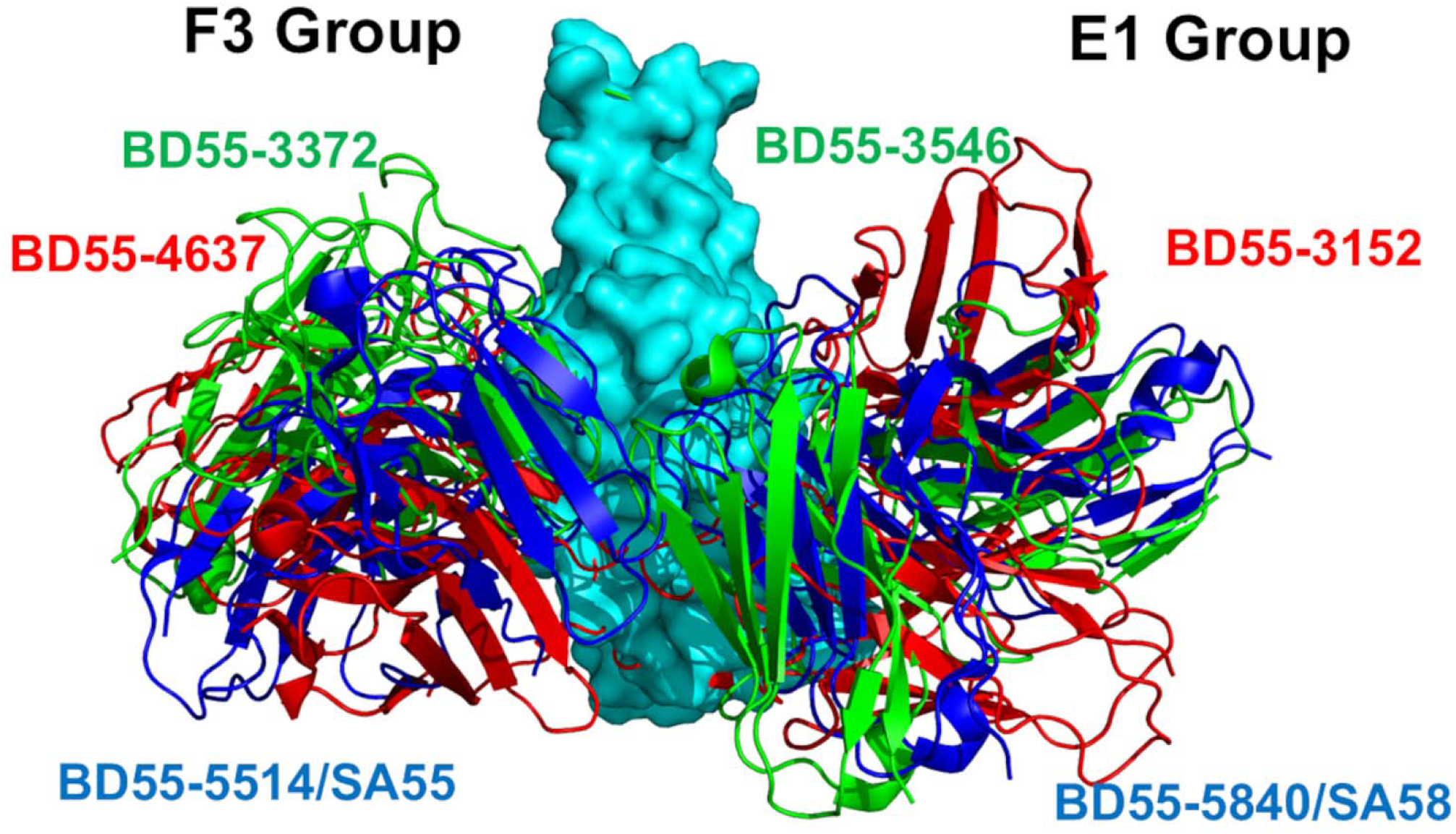
Structural organization of the SARS-CoV-2-RBD complexes with the E1 and F3 groups of Abs. Structural alignment of the E1 and F3 Abs. The S-RBD structure is shown in cyan surface. The F3 Abs are BD55-3372, pdb id 7WRO (in green ribbons). BD55-4637, 7WRJ (in red ribbons) and BD55-5514/SA555, pdb id 7Y0W (in blue ribbons). The corresponding E1 Abs are BD55-3546, pdb id 7WRY (in green ribbons), BD55-3152, pdb id 8WR8 (in red ribbons), an BD55-5840/SA58, pdb id 7Y0W (in blue ribbons). For clarity of presentation the heavy and light chains for the corresponding Abs are shown in the same color.

**Table 1.** Mutational landscape of the Omicron variants.

| Variant | Mutational landscape |
| --- | --- |
| BA.1 | A67, T95I, G339D, S371L, S373P, S375F, K417N, N440K, G446S, S477N, T478K, E484A, Q493R, G496S, Q498R, N501Y, Y505H, T547K, D614G, H655Y, N679K, P681H, N764K, D796Y, N856K, Q954H, N969K, L981F |
| BA.2 | T19I, G142D, V213G, G339D, S371F, S373P, S375F, T376A, D405N, R408S, K417N, N440K, S477N, T478K, E484A, Q493R, Q498R, N501Y, Y505H, D614G, H655Y, N679K, P681H, N764K, D796Y, Q954H, N969K |
| XBB.1.5 | T19I, V83A, G142D, Del144, H146Q, Q183E, V213E, G252V, G339H, R346T, L368I, S371F, S373P, S375F, T376A, D405N, R408S, K417N, N440K, V445P, G446S, N460K, S477N, T478K, E484A, F486P, F490S, R493Q reversal, Q498R, N501Y, Y505H, D614G, H655Y, N679K, P681H, N764K, D796Y, Q954H, |
|  | N969K |
| BA.2.86 | T19I, R21T, S50L, del69-70,V127F, delY144, F157S, R158G, delN211, L213I, L226F, H25N,A264D, I332V, D339H, K356T, R403K, V445H, G446S, N450D, L452W, N460K, N481K, del V483, A484K, F486P, R493Q, E554K, A570V, P612S, I670V, H68R, D939F, P1143L |
| JN.1 | T19I, R21T, S50L, del69-70,V127F, delY144, F157S, R158G, delN211, L213I, L226F, H25N, A264D, I332V, D339H, K356T, R403K, V445H, G446S, N450D, L452W, **L455S**, N460K, N481K, del V483, A484K, F486P, R493Q, E554K, A570V, P612S, I670V, H68R, D939F, P1143L |
| KP.2 | **JN.1 + S:R346T, S:F456L, S:V1104L**<br>T19I, R21T, S50L, del69-70,V127F, delY144, F157S, R158G, delN211, L213I, L226F, H25N,A264D, I332V, D339H, **R346T**, K356T, R403K, V445H, G446S, N450D, L452W, **L455S, F456L**, N460K, N481K, del V483, A484K, F486P, R493Q, E554K, A570V, P612S, I670V, H68R, D939F, **V1104L**, P1143L |
| KP.3 | **JN.1 + S:F456L, S:Q493E, S:V1104L**<br>T19I, R21T, S50L, del69-70,V127F, delY144, F157S, R158G, delN211, L213I, L226F, H25N,A264D, I332V, D339H, K356T, R403K, V445H, G446, N450D, L452W, **L455S, F456L**, N460K, N481K, del V483, A484K, F486P, **Q493E**, E554K, A570V, P612S, I670V, H68R, D939F, **V1104L**, P1143L |
| LB.1 | **JN.1+ S:S31-, S:Q183H, S:R346T, S:F456L**<br>T19I, R21T, **S31-**, S50L, del69-70,V127F, delY144, F157S, R158G, **Q183H**, delN211, L213I, L226F, H25N,A264D, I332V, D339H, **R346T**, K356T, R403K, |
|  | V445H, G446S, N450D, L452W, **L455S, F456L**, N460K, N481K, del V483, A484K, F486P, R493Q, E554K, A570V, P612S, I670V, H68R, D939F, P1143L |
| XEC | **JN.1 + S:T22N, S:F59S, S:F456L, S:Q493E, S:V1104L**<br>T19I, R21T, **T22N**, S50L, **F59S**, del69-70, V127F, delY144, F157S, R158G, delN211, L213I, L226F, H25N, A264D, I332V, D339H, K356T, R403K, V445H, G446S, N450D, L452W, **L455S**, F456L, N460K, N481K, del V483, A484K, F486P, **Q493E**, E554K, A570V, P612S, I670V, H68R, D939F, **V1104L**, P1143L |

Using dynamic ensembles of the S-Ab complexes and systematic mutational scanning of the RBD residues for binding with Abs, we characterize patterns of mutational sensitivity and compute mutational scanning heatmaps to identify escape hotspot centers. We also employ the Molecular Mechanics/Generalized Born Surface Area (MM-GBSA) approach for rigorous binding affinity computations of the S-Ab complexes and residue-based energy decomposition. Using these approaches, we examine the energetic determinants by which broadly potent Abs can largely evade immune resistance from the RBD sites of Omicron mutations. We show that E1 and F3 groups of Abs (Figure 1) targeting binding epitopes may leverage primarily strong hydrophobic interactions with the binding epitope hotspots critical for the RBD stability and ACE2 binding, while escape mutations tend to emerge in sites associated with synergistically strong hydrophobic and electrostatic interactions. Our results may be helpful in rationalizing the roles and balancing act of hydrophobic and electrostatic drivers of binding as well as to guide engineering of Abs by exploiting electrostatic complementarity to the RBD target sites, mitigating Ab resistance by engineering negatively charged neutralizing Abs with high affinity and targeting conserved epitopes void of Ab unfavorable interfaces with positively charged RBD residues. These findings suggest that RBD mutations and their associated immune escape may have been reaching a certain level of stability with recently emerged lineages showing a greater diversity in terms of their composition of ionizable amino acids. We argue that under the constraint of the RBD stability and ACE2 binding the evolution pressures may now explore epistatic effects and localized electrostatic changes to boost immune evasion. The insights from this investigation suggest therapeutic venues for targeted exploitation of the binding hotspots and allosteric communication centers that may potentially aid in evading drug resistance.

## 2. Materials and Methods

### Molecular Dynamics Simulations

The crystal and cryo-EM structures of the Omicron RBD-Ab complexes are obtained from the Protein Data Bank [83]. All-atom MD simulations were performed for the E1 group Abs (BD55-3546 Fab bound to SARS-COV2 Delta RBD complex, pdb id 7WRY; BD55-3152 Fab bound to Omicron BA.1, pdb id 7WR8, and SA58 bound to Omicron BA.1, pdb id 7Y0W) and F3 Abs (BD55-3372 bound to SARS-CoV-2 delta RBD, pdb id 7WRO, BD55-4637 Fab bound to Omicron BA.1, pdb id 7WRJ and SA55 bound to Omicron BA.1, pdb id 7Y0W). For simulated structures, hydrogen atoms and missing residues were initially added and assigned according to the WHATIF program web interface [84]. The missing regions are reconstructed and optimized using template-based loop prediction approach ArchPRED [85]. The side chain rotamers were refined and optimized by SCWRL4 tool [86]. The protonation states for all the titratable residues of the Ab and RBD proteins were predicted at pH 7.0 using Propka 3.1 software and web server [87,88]. The protein structures were then optimized using atomic-level energy minimization with composite physics and knowledge-based force fields implemented in the 3Drefine method [89,90]. We considered glycans that were resolved in the structures. NAMD 2.13-multicore-CUDA package [91] with CHARMM36 force field [92] was employed to perform 1µs all-atom MD simulations for the RBD-Ab complexes. The structures of the complexes were prepared in Visual Molecular Dynamics (VMD 1.9.3) [93] and with the CHARMM-GUI web server [94,95] using the Solutions Builder tool. Hydrogen atoms were modeled onto the structures prior to solvation with TIP3P water molecules [96] in a periodic box that extended 10 Å beyond any protein atom in the system. To neutralize the biological system before the simulation, Na^+^ and Cl^−^ ions were added in physiological concentrations to achieve charge neutrality, and a salt concentration of 150 mM of NaCl was used to mimic physiological concentration. All Na^+^ and Cl^−^ ions were placed at least 8 Å away from any protein atoms and from each other. MD simulations are typically performed in an aqueous environment in which the number of ions remains fixed for the duration of the simulation, with a minimally neutralizing ion environment or salt pairs to match the macroscopic salt concentration [97]. All protein systems were subjected to a minimization protocol consisting of two stages. First, minimization was performed for 100,000 steps with all the hydrogen-containing bonds constrained and the protein atoms fixed. In the second stage, minimization was performed for 50,000 steps with all the protein backbone atoms fixed and for an additional 10,000 steps with no fixed atoms. After minimization, the protein systems were equilibrated in steps by gradually increasing the system temperature in steps of 20 K, increasing from 10 K to 310 K, and at each step, a 1ns equilibration was performed, maintaining a restraint of 10 kcal mol^−1^ Å^−2^ on the protein C_α_ atoms. After the restraints on the protein atoms were removed, the system was equilibrated for an additional 10 ns. Long-range, non-bonded van der Waals interactions were computed using an atom-based cutoff of 12 Å, with the switching function beginning at 10 Å and reaching zero at 14 Å. The SHAKE method was used to constrain all the bonds associated with hydrogen atoms. The simulations were run using a leap-frog integrator with a 2 fs integration time step. The ShakeH algorithm in NAMD was applied for the water molecule constraints. The long-range electrostatic interactions were calculated using the particle mesh Ewald method [98] with a cut-off of 1.0 nm and a fourth-order (cubic) interpolation. The simulations were performed under an NPT ensemble with a Langevin thermostat and a Nosé–Hoover Langevin piston at 310 K and 1 atm. The damping coefficient (gamma) of the Langevin thermostat was 1/ps. In NAMD, the Nosé–Hoover Langevin piston method is a combination of the Nosé– Hoover constant pressure method [99] and piston fluctuation control implemented using Langevin dynamics [100,101]. An NPT production simulation was run on equilibrated structures for 1µs keeping the temperature at 310 K and a constant pressure (1 atm).

### Binding Free Energy Computations: Mutational Scanning and Sensitivity Analysis

We conducted mutational scanning analysis of the binding epitope residues for the S RBD-Ab complexes. Each binding epitope residue was systematically mutated using all substitutions and corresponding protein stability and binding free energy changes were computed. BeAtMuSiC approach [102–104] was employed that is based on statistical potentials describing the pairwise inter-residue distances, backbone torsion angles and solvent accessibilities, and considers the effect of the mutation on the strength of the interactions at the interface and on the overall stability of the complex. The binding free energy of protein-protein complex can be expressed as the difference in the folding free energy of the complex and folding free energies of the two protein binding partners:

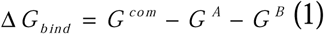

The change of the binding energy due to a mutation was calculated then as the following:

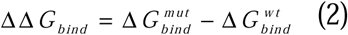

We leveraged rapid calculations based on statistical potentials to compute the ensemble-averaged binding free energy changes using equilibrium samples from simulation trajectories. The binding free energy changes were obtained by averaging the results over 1,000 and 10, 000 equilibrium samples for each of the systems studied.

### Binding Free Energy Computations

We calculated the ensemble-averaged changes in binding free energy using 1,000 equilibrium samples obtained from simulation trajectories for each system under study. Initially, the binding free energies of the RBD-Ab complexes were assessed using the MM-GBSA approach [105,106]. Additionally, we conducted an energy decomposition analysis to evaluate the contribution of each amino acid during the binding of RBD to Abs [107,108]. The binding free energy for the RBD-Ab complex was obtained using:

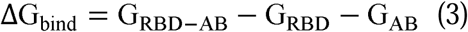

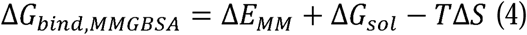

where Δ*E_MM_* is total gas phase energy (sum of Δ*E_internal_*, Δ*E_electrostatic_*, and Δ*Evdw*); Δ*Gsol* is sum of polar (Δ*G_GB_*) and non-polar (Δ*G_SA_*) contributions to solvation. Here, G_RBD–AB_ represent the average over the snapshots of a single trajectory of the complex, G_RBD_ and G_AB_ corresponds to the free energy of RBD and Ab protein, respectively. The polar and non-polar contributions to the solvation free energy is calculated using a Generalized Born solvent model and consideration of the solvent accessible surface area [109]. MM-GBSA is employed to predict the binding free energy and decompose the free energy contributions to the binding free energy of a protein– protein complex on per-residue basis. The binding free energy with MM-GBSA was computed by averaging the results of computations over 1,000 samples from the equilibrium ensembles. First, the computational protocol must be selected between the “single-trajectory” (one trajectory of the complex), or “separate-trajectory” (three separate trajectories of the complex, receptor and ligand). To reduce the noise in the calculations, it is common that each term is evaluated on frames from the trajectory of the bound complex. In this study, we choose the “single-trajectory” protocol, because it is less noisy due to the cancellation of intermolecular energy contributions. This protocol applies to cases where significant structural changes upon binding are not expected. Hence, the reorganization energy needed to change the conformational state of the unbound protein and ligand are also not considered. Entropy calculations typically dominate the computational cost of the MM-GBSA estimates. Therefore, it may be calculated only for a subset of the snapshots, or this term can be omitted [110,111]. However, for the absolute affinities, the entropy term is needed, owing to the loss of translational and rotational freedom when the ligand binds. In this study, the entropy contribution was not included in the calculations of binding free energies of the RBD-Ab complexes because the entropic differences in estimates of relative binding affinities are expected to be small owing to small mutational changes and preservation of the conformational dynamics [110,111]. MM-GBSA energies were evaluated with the MMPBSA.py script in the AmberTools21 package [112] and gmx_MMPBSA, a new tool to perform end-state free energy calculations from CHARMM and GROMACS trajectories [113].

## 3. Results

### Evolutionary Survey and Analysis of Omicron JN.1 Lineages

We first examined the evolutionary traits of SARS-CoV-2 lineages, particularly among the Omicron variants that are illustrated by the phylogenetic analysis using their corresponding clades nomenclature from Nextstrain an open-source project for real time tracking of evolving pathogen populations (https://nextstrain.org/) [114]. Nextstrain provides dynamic and interactive visualizations of the phylogenetic tree of SARS-CoV-2, enabling exploration and illustration of the evolutionary relationships between different lineages and variants. An overview of the phylogenetic analysis and SARS-CoV-2 clade classification (Figure 2, Supporting Information, Figure S1) highlights the evolution of SARSB-CoV-2 lineages. Clades 19A and 19B are ancestor lineages that emerged in Wuhan. Clades from 20D to 20J include Alpha (lineage B.1.1.7), Beta (lineage B.1.351), Gamma (lineage P.1). Clades from 21A to 21J include the Delta, Lambda (lineage C.37), Mu (lineage B.1.621), and Epsilon (lineages B.1.429). Clades 21K (BA.1) and 21L (BA.2) are Omicron sublineages emerged from the strain 21M (lineage B.1.1.529).

**Figure 2.**
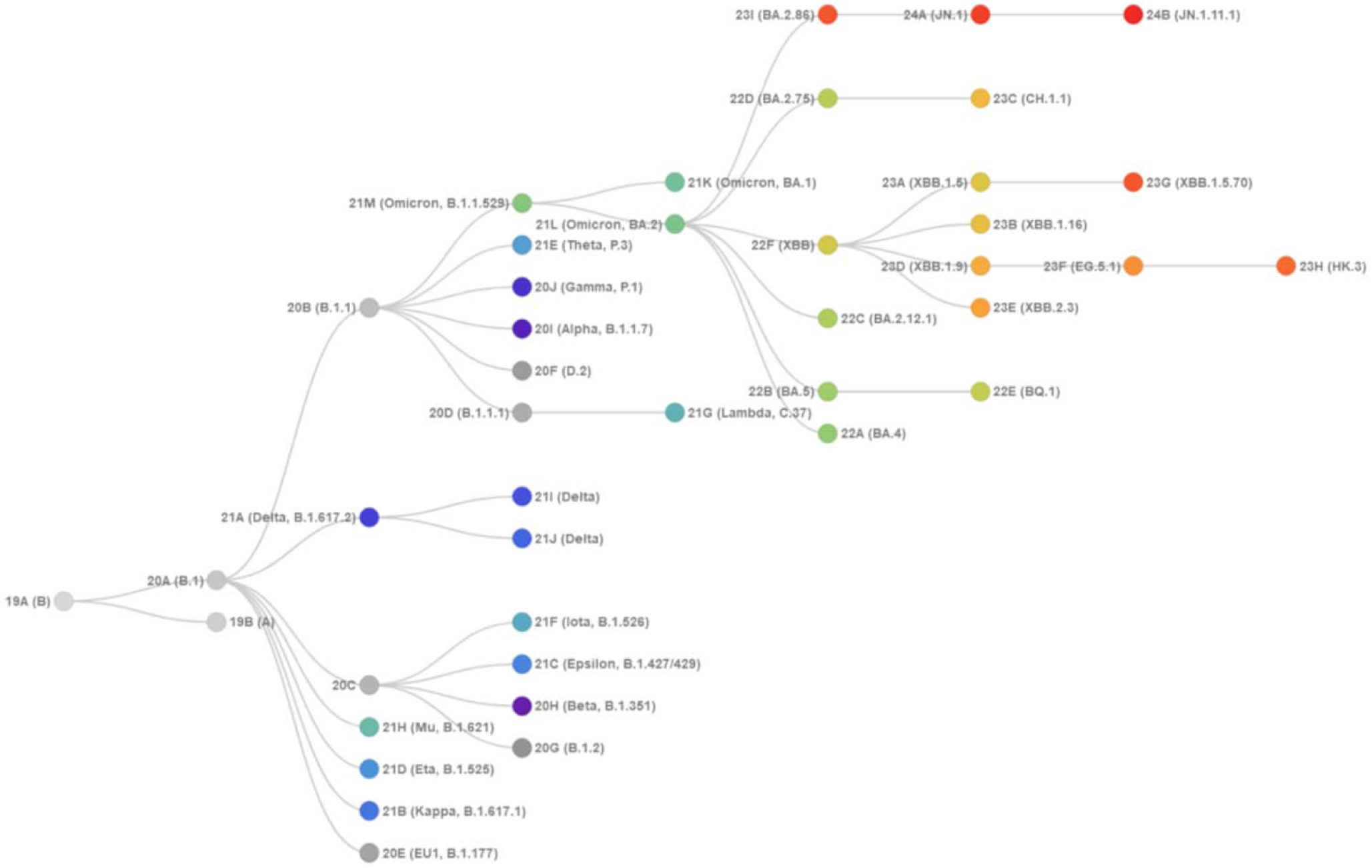
Phylogenetic relationships of existing SARS-CoV-2 clades. The graph is generated using Nextstrain, an open-source project for real time tracking of evolving pathogen populations (https://nextstrain.org/). This figure was generated using ncov-clade-schema https://ncov-clades-schema.vercel.app/

JN.1 is a variant of BA.2.86 which emerged independently from Omicron BA.2 and received Nextstrain clade classification 24A (Figure 2). JN.1 diversified into a number of sublineages that share recurrent mutations R346T (JN.1.18), F456L (JN.1.16), T572I (JN.1.7), or combinations of these mutations (KP.2 : JN.1 + S:R346T, S:F456L, S:V1104L) which was designated as Nextstrain clade 24B. KP.3 (JN.1.11.1.3), featured mutations R346T, L455S, F456L, Q493E and V1104L. KP.3 (JN.1+ S:F456L, S:Q493E, S:V1104L) received Nextstrain clade 24C. Clade 24G, lineage KP.2.3, is a descendant of clade 24B with extra spike substitutions S:R346T, S:H146Q and deletion S:S31-, and ORF3a:K67N (https://github.com/nextstrain/ncov/pull/1152). Clade 24D is XDV.1 variant and clade 24E is KP.3.1.1 variant (Supporting Information, Figure S1A-C).

The most recent designated clade 24F of lineage XEC that resulted from a recombination event of KS.1.1 and KP.3.3. KS.1.1 belongs to clade 24A (lineage JN.1) and KP.3.3 belongs to clade 24C (lineage KP.3), which itself descends from JN.1. Hence, XEC is a recombinant of JN.1-derived diversity. According to Coronavirus Network (CoViNet) (https://data.who.int/dashboards/covid19/variants) currently circulating COVID-19 VOIs are JN.1.7 (JN.1 + S:T572I, S:E1150D clade 24A), KP.2 (JN.1 + S:R346T, S:F456L, S:V1104L, clade 24B). KP.3 (JN.1 + S:F456L, S:Q493E, S:V1104L, clade 24C), KP.3.1.1 (KP.3 + S:S31-, clade 24C), JN.1.18 (JN.1 + S:R346T, clade 24A), LB.1 (JN.1 + S:S31-, S:Q183H, S:R346T, S:F456L, clade 24A), and XEC (JN.1 + S:T22N, S:F59S, S:F456L, S:Q493E, S:V1104L, clade 24F) (Figure 1).The current evolutionary divergences between XBB and JN.1 lineages illustrated by Nextstrain diagrams (Supporting Information, Figure S1C,D) indicate that evolutionary trajectories of Omicron lineages can proceed through complex recombination, antigenic drift and convergent evolution. The recent data on growth advantage relative to population average in US showed that XEC and KP.3.1.1 are the most common Pango lineage and have the highest fitness relative to the population average (https://github.com/nextstrain/ncov/pull/1152) (Supporting Information, Figure S1D). According to WHO Coronavirus Network (CoViNet) (https://data.who.int/dashboards/covid19/variants) in the recent month analysis of prevalence of SARS-CoV-2 variants of interest (VOI) and variants under monitoring (VUM) the dominant variants are KP.3.1.1 (Prevalence: 53.59%, VUM), XEC (Prevalence: 26.14%, VUM) JN.1 (Prevalence: 9.8%, VOI), KP.3 ((JN.1 + S:F456L, S:Q493E, S:V1104L; Prevalence: 4.14%, VUM), KP.2 (Prevalence: 1.53%, VUM), JN.1.18 ((JN.1 + S:R346T; Prevalence: 0.87%, VUM), and BA.2.86 (Prevalence: 0.22%, VOI). As of December 2024, current circulating variants in US are dominated by KP.3.1.1 (32.6%) XEC (19.5%) MC.16 (3.4%) MC.13 (2.7%) NL.2 (2.6%) MC.1 (2.2%) MC.10.1 (2%) XEC.2 (1.6%) MC.11 (1.5%) KP.2.3 (1.4%) LB.1.3.1 (1.4%) an XEK (1.3%) (https://public.tableau.com/app/profile/raj.rajnarayanan/viz/USAVariantDB/VariantDashboard). According to Cov-Spectrum data (https://cov-spectrum.org/explore/World/AllSamples/) the JN.1 variant accounts for 65.4% of the samples collected and analyzed within the specified date range (June 3, 2024, to November 26, 2024). This percentage represents the prevalence of the JN.1 variant among all the samples in the world tested during that period (Supporting Information, Figure S2). It is instructive to notice that the high proportion of the world samples collected and analyzed during this latest period is dominated by other JN.1 descendants, particularly 54.8% for KP.3, 49.2 % for KP.3.1.1 and 31.5 % for XEC variant (Supporting Information, Figure S2). Notice that the sum of percentages for the presented variants on Cov-Spectrum can exceed 100% because some samples may contain multiple variants. The Cov-Spectrum comparison showed the prevalence of different variants across various regions and helps to visualize how specific variants are spreading globally and to identify regional differences in variant distribution (Supporting Information, Figure S2). The proportion of sequences over the last six months showed a general decline for JN.1, FLiRT and KP.3 variants (Supporting Information, Figure S2). The current lineage frequency is considerably increased for the presently dominant XEC variant (Supporting Information, Figure S2). KP.3.1.1 is KP.3 that additionally convergently acquired S31 deletion have emerged as advantageous mutation and currently features high lineage frequency (Supporting Information, Figure S1D, S2).

### Structural Analysis of the RBD Complexes with E1 and F3 Group Abs

We began with the structural analysis of the S-RBD binding with the two classes of Abs E and F [54–58]. Groups E and F are categorized into E1–E3 and F1–F3, covering the front and back of the RBD. We specifically focused on two groups: E1 group Abs (BD55-3546 Fab bound to SARS-COV2 Delta RBD complex, pdb id 7WRY; BD55-3152 Fab bound to Omicron BA.1, pdb id 7WR8, and SA58 bound to Omicron BA.1, pdb id 7Y0W) and F3 Abs (BD55-3372 bound to SARS-CoV-2 delta RBD, pdb id 7WRO, BD55-4637 Fab bound to Omicron BA.1, pdb id 7WRJ and SA55 bound to Omicron BA.1, pdb id 7Y0W) (Figure 1). These two groups of Abs emerged as a result of high-throughput epitope determination and selection of broadly neutralizing Abs that target rare RBD epitopes that are associated with critical functions, such as ACE2 binding and RBD stability to avoid immunity-directed escape mutations [54,57].

The E1 group of Abs binds to the other side of the RBD and is known to be susceptible to mutations of T345 and R346 [54,57]. BD55-3546 binds to the binding epitope consisting of residues 340-347, 440-451 and P499 (Figure 3, Table S1). The topology of the binding epitope reveals two major patches where one corresponds to residues 340-347 and 356, while the second separate patch is formed by residues 440-451 and 499 (Figure 3, Table 1). This Ab makes extensive contacts to the large number of the RBD residues, where continuous stretches of the binding epitope are formed by residues 340-347 and especially 440-451 (Figure 3).

**Figure 3.**
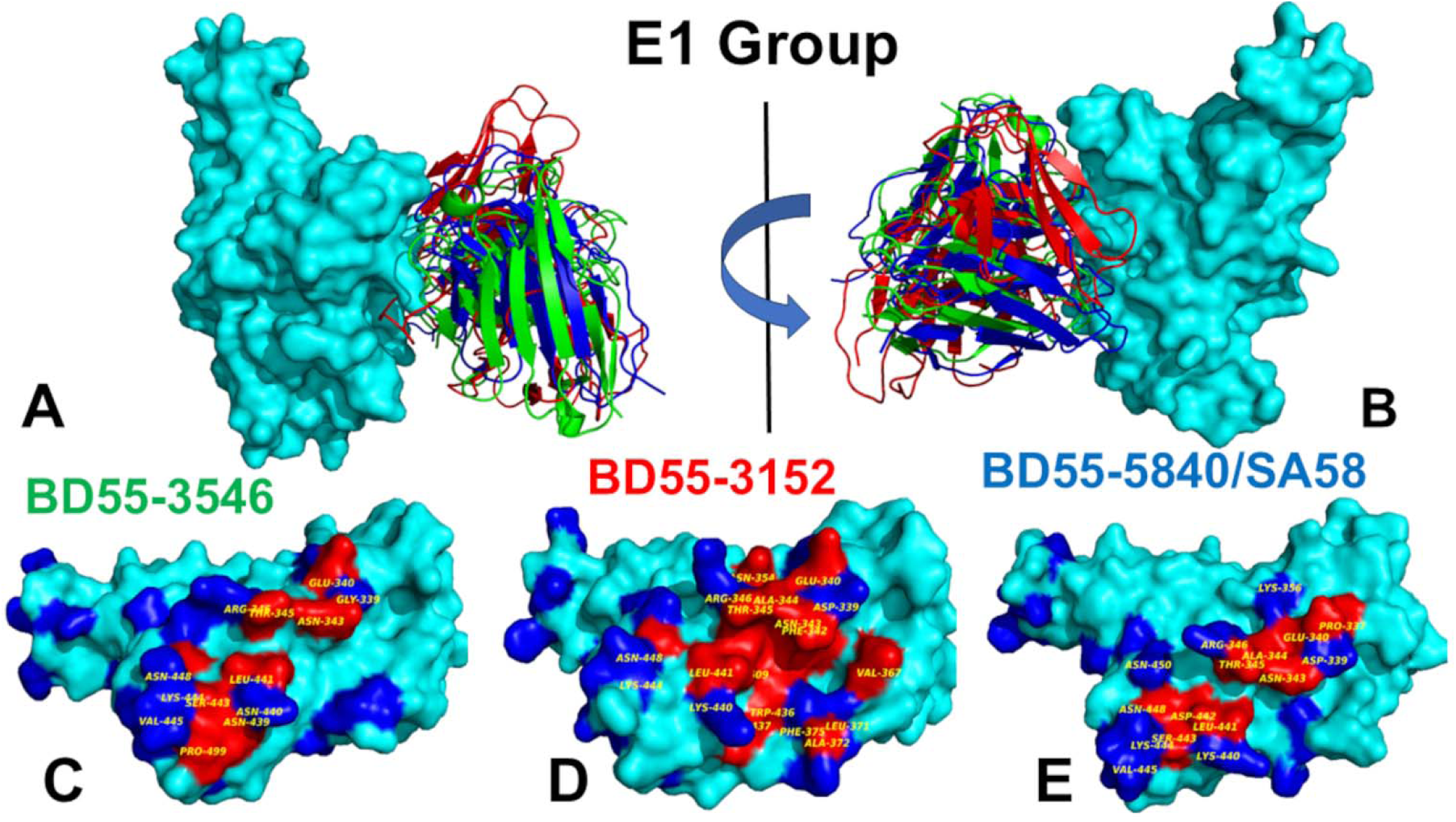
Structural organization of the RBD complexes and binding epitopes of the E1 group of Abs. (A,B) The two views of the RBD-Ab complexes. The S-RBD structure is shown in cyan surface. The E1 Abs are BD55-3546, pdb id 7WRY (in green ribbons), BD55-3152, pdb id 8WR8 (in red ribbons), an BD55-5840/SA58, pdb id 7Y0W (in blue ribbons). For clarity of presentation the heavy and light chains for the corresponding Abs are shown in the same color. (C) The RBD surface and binding epitope for BD55-3546. RBD is shown in cyan surface. The binding epitope residues are in red surface (G339, E340, V341, N343, A344, T345, R346, F347, K356, N439, N440, L441, D442, S443, K444, V445, N448, N450, Y41, P499, R509). (D) The RBD surface and binding epitope for BD55-3152. RBD is shown in cyan surface. The binding epitope residues are in red surface (D339, E340, V341, F342, N343, A344, T345, R346, F347, N354, V367, L368, L371, F375, W436, N437, S438, K440, L441, D442, K444, N448, Y451, R509). (E) The RBD surface and binding epitope for SA58. RBD is shown in cyan surface. The binding epitope residues are in red surface (P337 D339, E340, V341, N343, A344, T345, R346, K356, R357, I358, K440, L441, D442, S443, K444, V445, N448, N450, Y451, R509).The key sites of Omicron XBB, BA.2.86, and JN.1 lineages are shown in blue surface (residues 339, 346, 356, 371, 373, 375 376, 403, 405, 408, 417, 440, 444, 445, 446, 450, 452, 455, 456, 460, 475, 477, 478, 481, 484, 486, 493, 498, 501, 505).

Importantly, we found a large number of contacts made with T345 by R105, W94, Y103, L96, Y91 and N92 Ab residues. The fewer contacts are also formed with R346 position. R346 is particularly interesting as the FLiRT variant has an additional R346T mutation in the SLip backbone. R346T mutation is featured in KP.2 (JN.1 + S:R346T, S:F456L, S:V1104L), LB.1 (JN.1 + S:S31-, S:Q183H, S:R346T, S:F456L) and KP.2.3 variants but conspicuously absent in the KP.3 (JN.1 + S:F456L, S:Q493E, S:V1104L) and XEC variants (JN.1 + S:T22N, S:F59S, S:F456L, S:Q493E, S:V1104L). Another significant binding residue N440 is located in a pocket formed by the heavy chain residues W50, N52, T55, I57, P58 and T59 making extensive van der Waals and hydrogen-bond interactions with the Ab (Figure 3, Table S1). Another important hotspot K444 is involved in contacts with the heavy chain N31, N54 and L102 positions (Figure 3, Table S1). Interestingly, while most E1 Abs are sensitive to mutations of T345 and R346, BD55-3546 mainly escaped by T345 and N440 mutations [57]. It appears that these RBD positions form the largest number of contacts with BD55-3546 and are engaged in the dense interaction networks.

BD55-3152 targets a similar epitope that is more contiguous and even slightly larger than the one for BD55-3546 (Figure 3, Table S2). The epitope includes RBD residue regions 339-347, 354; 367-375; 436-448 and P509 (Figure 3). Hence, the BD55-3152 epitope includes additional residues 367-375 that overlap with the positions of Omicron mutations S371P and S373F. Structural analysis indicated that the densest contact networks for BD5-3152 with the RBD are formed by N343, A344, T345 and R346 (Figure 3, Table S2). In particular, T345 makes 10 strong contacts with Y33, Y31, G110, S111, P112, and L113 of the Ab (Table S2). Similarly, R346 is involved in multiple contacts with the Ab, particularly Q30 and D50 of the light chain (Table S2).

For SA58 the binding epitope is generally very similar, and includes residues 337-346, 356, and residues 440-450 (Figure 3, Table S3). Structural analysis showed that most contacts are made by E340, T345 and R346 RBD sites (Table S3). In particular, T345 is enclosed by Y93, L98, S94, N95, Y105, L94, and W96 and mutations in this position result in the immune escape [57]. Another sensitive site is K444 that forms interactions with T30, N32, S31, D54 and Y102 of the SA58 Ab (Table S3). According to the experimental data, neutralization by SA58 is affected by T345N, R346Q, R346T, E340D and escaped by E340K and K444E mutations [57]. Group E1 Abs are generally sensitive to the changes of G339, E340, T345 and especially R346 as revealed by their escaping mutation profiles [54,57]. According to escape calculator of the updated data from [46], the escape positions are indeed T345 and R346 [REF.]. In contrast to R346K, R346S/T would greatly compromise the binding activities of E1 Abs [57].

The F3 group BD55-3372 Ab binds to the conserved epitope formed by residues 372-375 and 404-408 as well as with the critical for ACE2 binding RBD segment 498-509 (Figure 4, Table S4). BD55-3372 exhibited high neutralization potency compared to other Abs in Group F3 [54]. BD55-4637 has a much larger binding epitope and forms stable contact with residues 374-378, 403-408, conserved stable residues 436-440, and ACE2 binding sites 500-508 (Figure 4, Table S5). BD55-4637 forms most of its contacts with the RBD residues critical for RBD stability and ACE2 binding while only several interactions are made with R403, D405 and R408 (Figure 4, Table S5). We also found that BD55-4637 makes multiple contacts with N439.

**Figure 4.**
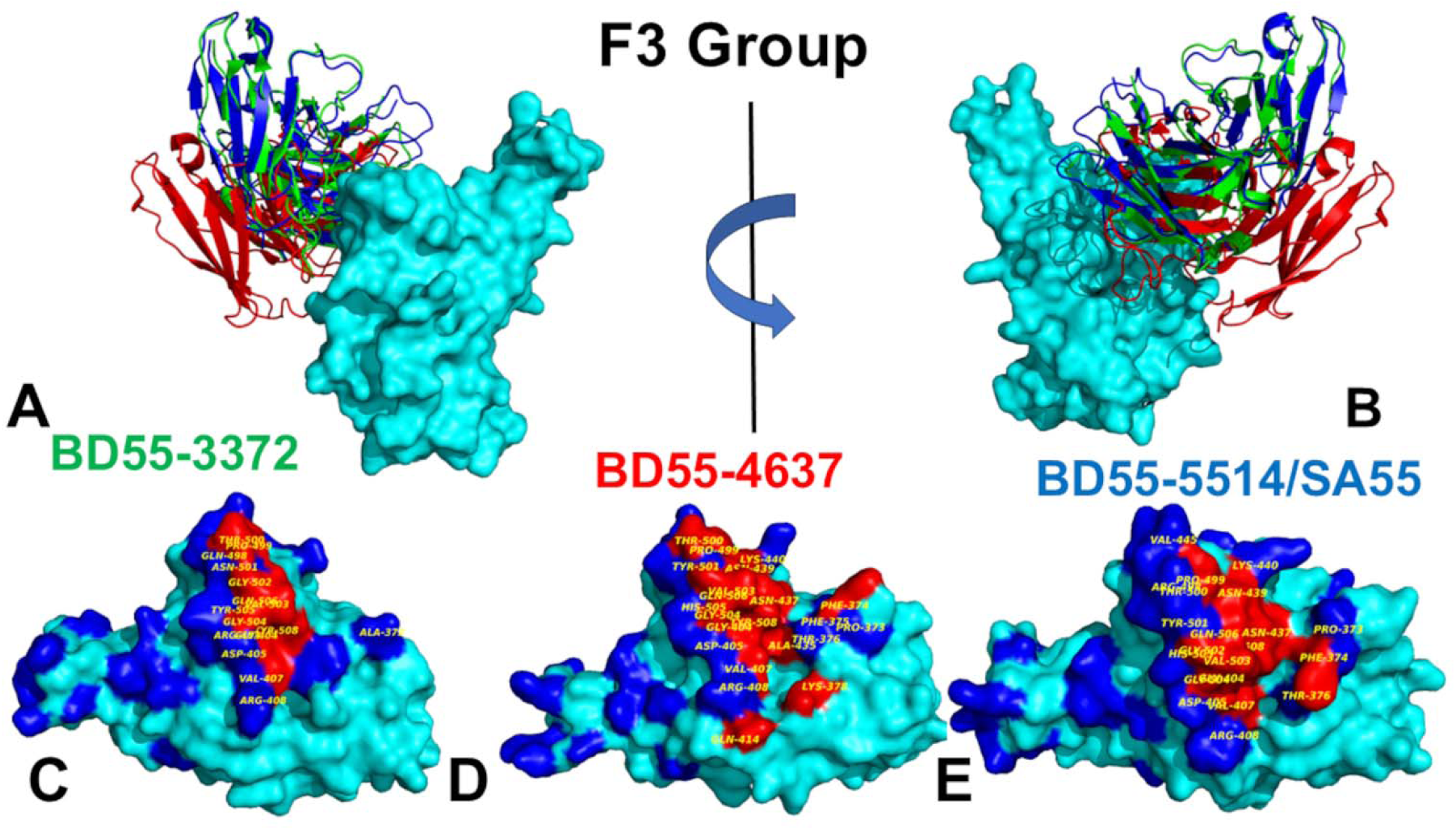
Structural organization of the RBD complexes and binding epitopes of the F3 group of Abs. (A,B) The two views of the RBD-Ab complexes. The S-RBD structure is shown in cyan surface. The E1 Abs are BD55-3372, pdb id 7WRO (in green ribbons), BD55-4637, pdb id 7WRJ (in red ribbons), an BD55-5514/SA55, pdb id 7Y0W (in blue ribbons). For clarity of presentation the heavy and light chains for the corresponding Abs are shown in the same color. (C) The RBD surface and binding epitope for BD55-3372. RBD is shown in cyan surface. The binding epitope residues are in red surface (A372, T403, G404, D405, E406, V407, R408, Q409, N437, N439, Q498, P499, T500, N501, G50, V503, G504, Y505, Q506, Y508). (D) The RBD surface and binding epitope for BD55-4637. RBD is shown in cyan surface. The binding epitope residues are in red surface (A37, P373, F374, F375, T376, F377, K378, R403, G404, D405, V407, R408, Q414, V433, A435, 436, N437, N439, K440, S496, R498, T500, Y501, G502, V503, G504, H505, Q506, Y508). (E) The RBD surface and binding epitope for SA55. RBD is shown in cyan surface. The binding epitope residues are in red surface (P373, F374, T376, R403, G404, D405, E406, V407, R408, N437, N439, K440, V445, Y495, 496, R498, P499, T500, Y501, G502, V503, G504, H505, Q506, Y508). The sites of Omicron XBB, BA.2.86, and JN.1 lineages are shown in blue surface (residues 339, 346, 356, 371, 373, 375 376, 403, 405, 408, 417, 440, 444, 445, 446, 450, 452, 455, 456, 460, 475, 477, 478, 481, 484, 486, 493, 498, 501, 505).

The structural analysis of the binding epitope for SA55 Ab showed RBD positions 373-376, 404-408, 436-440, 445,446, and extended RBD segment of residues 498-508 (Figure 4, Table S6). SA55 targets the critical for RBD functions segment 498-508 making multiple contacts with Y501, G502, V503, G504, H505 and T508 (Figure 4, Table S6). The core of the binding epitope corresponds to the conserved RBD region (residues 436-440) that is critical for RBD stability and ACE2-binding segment 498-508 where residues are unlikely to mutate to escape Abs as they are critical for ACE2 affinity. This segment includes Y501 and H505 energetic hotspots for ACE2 binding where the Ab competes with the host receptor for binding with the S-RBD, and as a result, these positions cannot be exploited by the virus for evolving Ab resistant mutations. However, there are a number of Omicron mutational sites such as S373P and S375F that interact with L94 and F55 of the heavy chain and where Omicron mutations could potentially alter binding with the Ab (Figure 4, Table S6).

Structural analysis also revealed multiple favorable contacts of SA55 with T376 using F55 of the heavy chain and interactions with D405 formed by L54, S31, T28, R30 of the heavy chain. Although T376, D405, and R408 are involved in the interaction with SA55, these residues are all located closer to the periphery of the binding epitope and some of the mutations in these positions may be tolerated. Indeed, according to the illuminating study by Cao and colleagues, SA55 displayed high potency against the Omicron subvariants sharing F456L mutation including HV.1 (L452R+F456L), JD.1.1(L455F/F456L+A475V) that typically greatly increase Ab evasion for other Abs [57]. Nonetheless, binding and neutralization activity of SA55 may be compromised by Y508H, G504S, K440E, and escaped by V503E and G504D [57].

Although structural analysis of the binding epitope and contacts alone is obviously not sufficient for prediction of escape hotspots and requires in-depth energetic assessment, the topological composition of the binding epitope revealed that positions V503 and G504 that are involved in many contacts with the Ab (Figure 4, Table S6) could be vulnerable to escape mutations. Indeed, we found that V503 makes multiple hydrophobic contacts with L54, P101, I52 and S31 while G504 is involved in interactions with L54, H32, S31 and R30 of the heavy chain (Figure 4, Table S6). In this context, it is not unexpected that changing the chemical identity of V503 to V503E and G504 to G504D can disrupt the interaction network and cause immune escape. In the next sections, we perform a detailed energetic analysis using full mutational scanning and MM-GBSA binding free energy calculations to dissect the effect of specific changes and contributions of the RBD residues on the binding affinity with E1 and F3 Abs.

### MD Simulations of the S-RBD Complexes with Abs Reveal Specific Patterns of Moderate RBD Mobility

All-atom MD simulations were performed for the S-RBD complexes with a panel of studied Abs to examine the conformational landscapes and determine specific dynamic signatures induced by Abs. Conformational dynamics profiles are described using the root mean square fluctuations (RMSF) obtained from simulations (Figure 5). Despite notable differences in the binding epitopes for F3 and E1 Abs, the conformational dynamics profiles for these Abs are fairly similar (Figure 5). The conformational mobility distributions for the S-RBD complex with E1 and F3 Abs displayed characteristic for the RBD minima corresponding to residues 374-377, the RBD core residue cluster (residues 396-403), residues 445-456 (Figure 5). The RBD core consisting of antiparallel β strands (β1 to β4 and β7) (residues 354-358, 376-380, 394-403, 431-438, 507-516). The β-sheets β5 and β6 (residues 451-454 and 492-495) correspond to highly stable positions that that anchor the RBM region to the central core.

**Figure 5.**
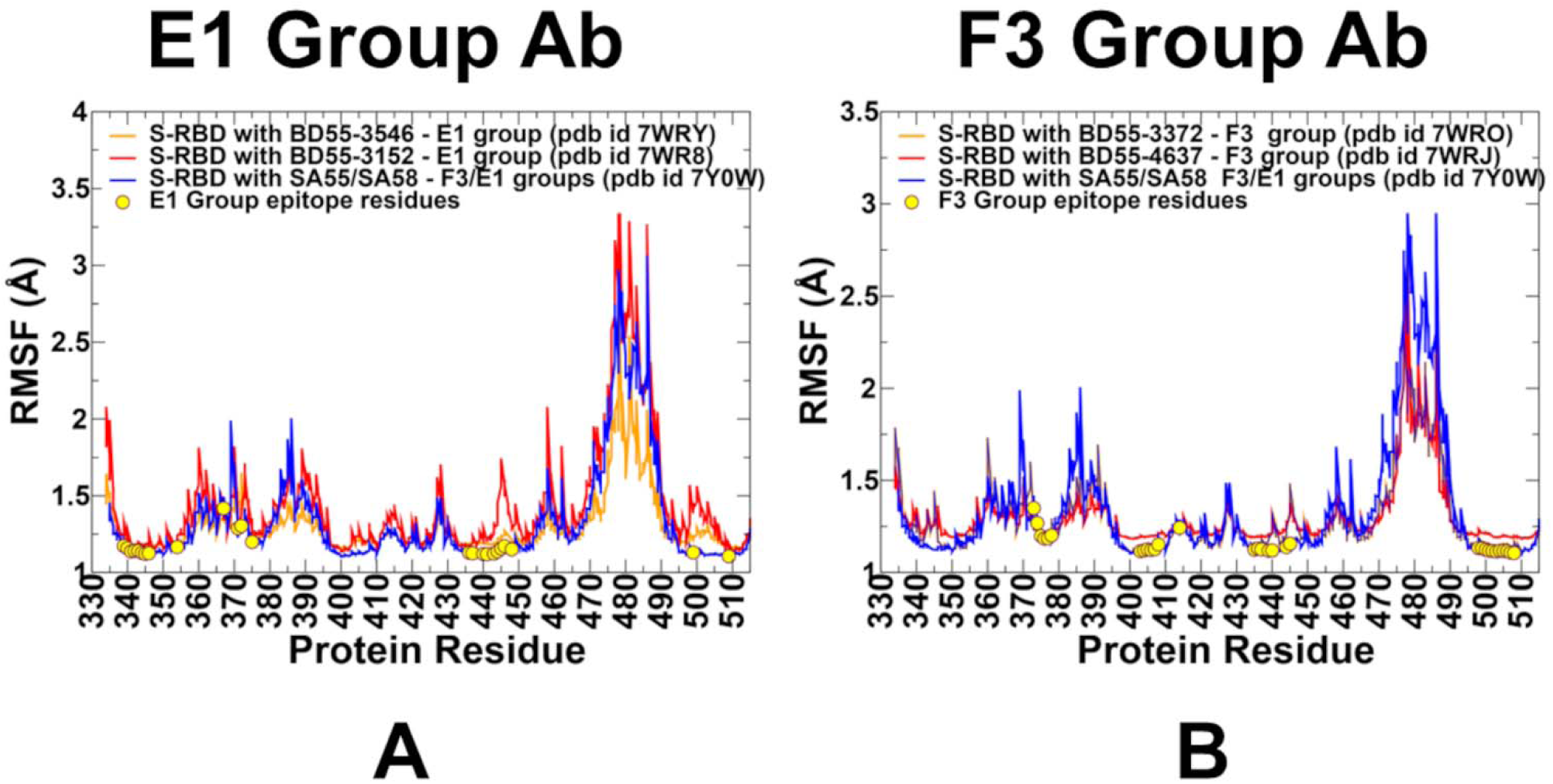
Conformational dynamics profiles obtained from simulations of the RBD-Ab complexes. (A) The RMSF profiles for the RBD residues obtained from MD simulations of the S-RBD complexes with E1 group Abs : BD55-3546 (in orange lines), BD55-3152 (in red lines), an BD55-5840/SA58, pdb id 7Y0W (in blue lines). (B) The RMSF profiles for the RBD residues obtained from MD simulations of the S-RBD complexes with F3 group Abs : BD55-3372 (in green lines). BD55-4637, 7WRJ (in red lines) and BD55-5514/SA555, pdb id 7Y0W (in blue ribbons). The positions of the binding epitope residues for E1 and F3 group of Abs are highlighted in yellow filled circles on panels (A,B).

For E1 group, BD55-3152 targets large epitope formed by RBD regions 339-347, 367-375; 436-448. The result of structural analysis showed significant contacts with T345 and R346 positions (Figure 3, Table S2). However, we observed appreciable fluctuations of residues 367-375 and 440-448 in this complex as well as in positions 458-460 (Figure 5A). In addition, the dynamics profile for this complex exhibited markedly increased RBM mobility (residues 470-490) as compared to another Ab BD55-3546 from this group (Figure 5A). Group E1 Abs are also generally sensitive to mutation of G339, E340, T345 and especially R346 [54,57] and according to our dynamics analysis these RBD positions become largely immobilized for all E1 Abs displaying only very small thermal fluctuations (Figure 5A). Hence, the dynamics profiles indicated that vulnerable to escape RBD positions in this group are associated with the epitope residues that become highly stable due to multiple contacts with the Abs, suggesting that mutations in these sites are likely to induce significant destabilizing effect. By projecting conformational fluctuation profiles onto structures of the E1 Ab-RBD complexes (Figure 6), we noticed considerable rigidification of the RBD with only the RBM region displaying high mobility.

**Figure 6.**
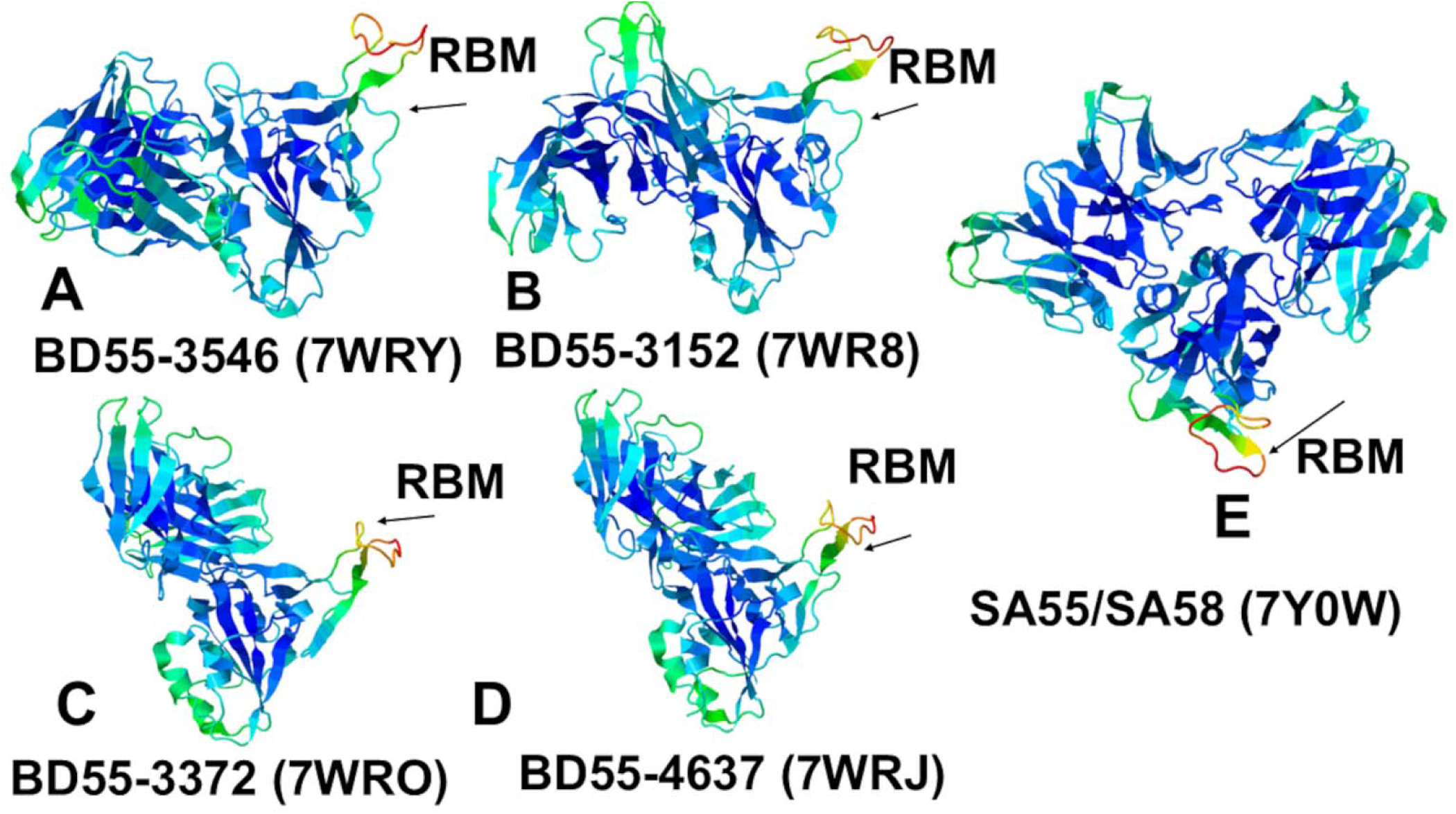
Conformational dynamics profiles projected onto structures of the RBD complexes with E1 group Abs : BD55-3546 (A), BD55-3152 (B), and RBD complexes with F3 group Abs : BD55-3372 (C), BD55-4637 (D). The conformational dynamics profile is projected onto RBD structure with the cocktail of SA55 and SA58 Abs (E). The structures are colored with the sliding color scale from blue (rigid regions) to red (flexible regions).

For the F3 Ab complexes, we also found that the binding epitope positions making contacts with the Abs are associated with stabilization of the corresponding regions (Figure 5B). Among F3 Abs, BD55-4637 exhibited an expanded binding epitope formed by residues 374-378, 403-408, 414, 436-445, and 500-508. Interestingly, the residues 439-446 in the complex with BD55-4637 showed some degree of plasticity despite interactions with the Ab. We also observed that the binding of F3 Abs still leaves some regions of the RBD moderately mobile including residues 365-385 and the RBM region (Figure 6C,D). At the same time, the RBD residues 400-456 are effectively immobilized in the BD55-4637 complex with the RBD (Figures 5B, 6C,D). Of special importance is the analysis of conformational dynamics profile for the RBD complex with Ab cocktail of SA55/SA58 in which SA55 belongs to the F3 group and SA58 is E1 group (Figures 5,6E). We found that binding these two Abs to different sides of the RBD can result in prominent stabilization of the RBD and changes in the RBD mobility, showing moderate fluctuations in the regions 380-395, 455-465 and the increased mobility in the RBM loops (residues 473-487) (Figures 5, 6E). Notably, SA55 neutralizing activity remains largely unimpaired upon S375F, R403K, D405N, R408S, N440K, V445H, Q498R, N501Y, and Y505H changes [57]. The potential positions of immune escape for SA55 are T376, K378, D405, R408 and G504, while accumulation of mutations on E340D/K, T345P, R346Q and K444N may affect the neutralization of SA58 [57]. According to the dynamics analysis, these positions belong to the RBD regions that are stabilized by the SA55/SA58 binding. Interestingly, R403, D405, R408, K440, V445 where mutations are tolerated by SA55 binding exhibited small thermal fluctuations (Figure 5). It is possible that for T376, D405 and R408 sites that are stable and involved in the interactions with SA55, their structural location at the periphery of SA55 binding epitope may enable a certain degree of tolerance to substitutions using conformational plasticity in these positions.

Another important observation of dynamic analysis for SA55/SA58 complex with the RBD is that stable positions implicated in immune escape are immediately adjacent to the RBD with the increased mobility, i.e. potentially susceptible sites of escape may correspond to local hinge sites that modulate movements of the RBD (Figure 5). Of particular notice is the significantly increased mobility of the intrinsically flexible RBM motif (residues 470-490) in the SA55/SA58-RBD complex that may allow for local changes in the RBM and modulation of the ACE2 binding affinity. While SA58 does not directly interfere with ACE2 and can engage both the up and down RBDs, SA55 only binds to the up RBD and can block ACE2 [57]. We found that structural rigidification of the RBD regions by SA55/SA58 may be counteracted through elevated RBM mobility where RBM may adopt a variety of conformations that may not be fully compatible with ACE2 binding, The increased flexibility of the RBM region may allow for hinge mechanism in which the rigid anchor residues 451-454 and 492-495 enable movements of the RBM that can expose and hide the ACE2 binding surface which was also observed in other MD simulations [115,116].

Using this dynamic mechanism, SA55/SA58 binding may accomplish both effective neutralization via interaction with conserved epitopes critical for spike function and also control the ACE2 binding by imposing dynamic changes in the exposed RBM conformations of the RBD-up. Our dynamics analysis may be useful in light of recent proposals by Bloom and colleagues that mutations that move the RBD more to an up conformation can generally increase binding to ACE2 by the complete S protein but may also increase Ab neutralization since the RBD in the up conformation exposes more neutralizing epitopes (https://jbloomlab.org/posts/2024-11-20_escapecalc_update.html). They suggested that most mutations at sites 403, 405, 503, 504, 505 may fall into this category as they affect RBD up-down movements and therefore may be constrained in the virus evolution. According to our results, F3 Ab binding may induce the elevated RBM mobility in the RBD-up conformation and modulation of the ACE2 exposure. At the same time, positions 503-505 are immobilized by SA55/SA58 binding and can form local hinges for controlling RBD movements. Combined, these factors suggest that E1 and F3 Abs may also constrain the ability of the virus to evolve mutations without sacrificing the ACE2 binding or potential exposure of neutralizing epitopes. In this context, it may be worth noting that recent highly circulating variants KP.3.1.1 and XEC tend to escape neutralization via NTD mutations that induce the RBD-down form and block neutralization at expense of efficient ACE2 binding [117,118].

### Mutational Profiling of Protein Binding Interfaces with E1 and F3 Abs

Using the conformational ensembles of the RBD complexes, we embarked on structure-based mutational analysis of the S protein binding with Abs. To provide a systematic comparison, we constructed mutational heatmaps for the RBD interface residues of the S complexes with E1 and F3 groups of Abs. We first analyzed mutational heatmap for the E1 group. The strongest destabilization mutations for BD55-3546 include T345A (ΔΔG = 2.38 kcal/mol), T345G (ΔΔG = 2.16 kcal/mol), T345E (ΔΔG = 2.16 kcal/mol), T345S (ΔΔG = 1.91 kcal/mol) and N440A (ΔΔG =1.89 kcal/mol) (Figure 7A). T345 is surrounded by R105, W94, Y103, L96, Y91 and N92 Ab residues making multiple favorable hydrogen bonding and van der Waals contacts. As a consequence, mutational scanning identified multiple highly destabilizing substitutions (Figure 7A). BD55-3546 is also sensitive to mutations of N440, as N440 protrudes into a pocket formed by W50, N52, T55, I57, and T59 and forms extensive van der Waals and hydrogen-bond interactions (Figure 7A). These results are consistent with the escape profiles showing that BD55-3546 has mainly escaped by T345 and N440 mutations [57]. These experimental studies also established that E1 Abs are predominantly susceptible to mutations of T345 and R346 residues [57]. Another E1 Ab BD55-3152 has binding energetic hotspots at T345, W436, L441 and 346 positions. For this Ab the largest destabilization changes are associated with mutations of T345, including T345E (ΔΔG = 3.17 kcal/mol), T345K (ΔΔG = 2.82 kcal/mol), T345A (ΔΔG = 2.58 kcal/mol) and T345D (ΔΔG = 2.32 kcal/mol) (Figure 7B).

**Figure 7.**
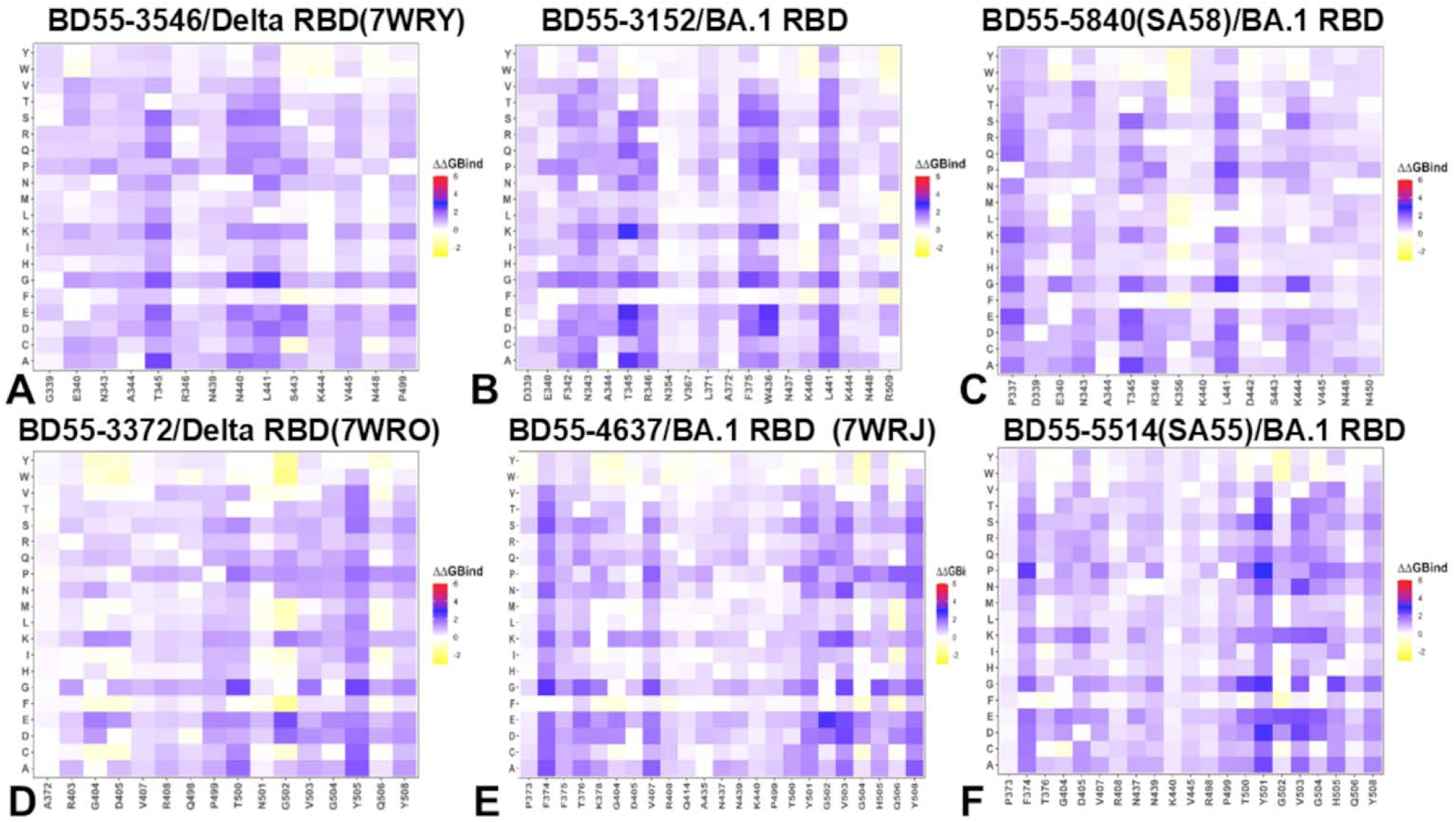
The ensemble-based mutational scanning of binding for the SARS-CoV-2 S-RBD complexes with E1 and F3 Abs. The mutational scanning heatmaps for the binding epitope residues in the S-RBD complexes with E1 Abs BD55-3546 (A), BD55-3152 (B), SA58 (C) and S-RBD complexes with F3 group Abs : BD55-3372 (D), BD55-4637 (E) and SA55 (F). The binding energy hotspots correspond to residues with high mutational sensitivity. The heatmaps show the computed binding free energy changes for 20 single mutations on the sites of variants. The squares on the heatmap are colored using a 3-colored scale blue-white-yellow, with yellow indicating the largest unfavorable effect on stability. The standard errors of the mean for binding free energy changes were based on a different number of selected samples from a given trajectory (500 and 1,000 samples) are within 0.04-0.13 kcal/mol.

Hence, for both BD55-3546 and BD55-3152 Abs T345 emerged as the dominant binding hotspot. As a result, the binding of these E1 Abs are highly sensitive and can be escaped by T345 mutations. Similarly, for SA58, the key binding hotspot is T345 where mutations T345A (ΔΔG = 2.36 kcal/mol), T345E (ΔΔG = 2.17 kcal/mol) and T345D (ΔΔG = 2.03 kcal/mol) lead to large destabilizing changes (Figure 7C) which is consistent with experiments showing that mutations of T345 result in significant escape from SA58.

In addition, we found that mutations of L441 may also reduce SA58 binding but the experiments showed only moderate sensitivity to mutations in this position. While T345 is clearly the dominant binding hotspot and key target for escape mutations, it was concluded that T345 substitutions are unlikely to emerge as this site is critical for the proper glycosylation of N343 that is essential for RBD function [57].

The mutational heatmap for F3 BD55-3372 showed large destabilization free energies for mutations of Y505A (ΔΔG = 2.16 kcal/mol), Y505C (ΔΔG = 2.01 kcal/mol) and Y505D changes as well as V503D (ΔΔG = 1.83 kcal/mol), Y508P (ΔΔG = 1.47 kcal/mol), G404E (ΔΔG = 1.67 kcal/mol) and G404K (ΔΔG = 1.52 kcal/mol) (Figure 7D). BD55-3372 is not susceptible to the changes of Y508 but are sensitive to the changes on Y505, V503 and G504 (Figure 7D). Nevertheless, these sites are critical for ACE2 binding and both V503 and G504 are conserved making it difficult for variants with mutations on these sites to emerge [57]. BD55-4637 has a rather unique escape profile, and is strongly affected by mutations in 404, 503, 505 and 508 positions [57]. The mutational map identifies as escape hotspots positions F374, V503, and Y508. Interestingly, mutations of V503 and Y508 are broadly destabilizing with particularly high unfavorable changes seen for V503D (ΔΔG = 2.38 kcal/mol), V503K (ΔΔG = 2.2 kcal/mol), V503N (ΔΔG = 2.24 kcal/mol), G504K (ΔΔG = 2.11 kcal/mol), Y508G (ΔΔG = 2.08 kcal/mol) and Y508N (ΔΔG = 1.97 kcal/mol) (Figure 7E). The results are consistent with the notion that BD55-4637 is sensitive to mutations of Y508 [57]. The detailed mutational heatmap of the SA55 interactions with the S protein showed large destabilization changes for Y501D (ΔΔG = 2.74 kcal/mol), Y501S (ΔΔG = 2.59 kcal/mol), V503D (ΔΔG = 2.41 kcal/mol), V503E (ΔΔG = 2.23 kcal/mol), and V503K mutations (ΔΔG = 2.02 kcal/mol) (Figure 7F). While these mutations are highly destabilizing for SA55 binding, these sites are fundamentally important for ACE2 binding and RBD functions. SA55 are sensitive to the changes on V503 and G504 but the effective mutations for destabilization of Ab binding are prohibitive as they would interfere with the key RBD functions [57]. As a result, evolution in these positions is highly constrained. A more detailed profiling of JN.1/KP.3 mutations against SA55 Ab showed only small destabilization changes upon mutations T376A (ΔΔG = 0.81 kcal/mol), R403K (ΔΔG = 0.65 kcal/mol), D405N (ΔΔG = 0.79 kcal/mol), R408S (ΔΔG = 0.34 kcal/mol), L455S (ΔΔG = 0.7 kcal/mol) and F456L (ΔΔG = 0.51 kcal/mol) (Figure 7pF). These changes reflect mostly a mild loss in the RBD stability and binding interactions, which is consistent with functional experiments showing group F3 Abs, such as SA55, are not sensitive to the D405N and R408S mutations of BA.2 making this Ab effective against a broad spectrum of recent variants from BA.2.86 to KP.2 and KP.3. To summarize, the results of mutational scanning using rapid computation of binding free energy changes revealed binding hotspots for E1 and F3 Abs that are consistent with the experimental DMS data and immune escape centers. In particular, consistent with the experiments [57], mutational scanning predicted T345 and R346 residues as dominant binding hotspots for E1 group of Abs. Similarly, our results predicted RBD sites V503, G504 and Y508 as the key binding hotspots for F3 group of Abs that agrees the experiments showing that these positions are major immune escape centers for these Abs.

### MM-GBSA Analysis of the Binding Affinities

The experimental and computational studies suggested that the cross-neutralization Ab activity against Omicron variants may be driven by balance and tradeoff of multiple energetic factors and interaction contributions of the evolving escape hotspots involved in antigenic drift and convergent evolution. However, the dynamic and energetic details quantifying the balance and contribution of these factors, particularly the balancing nature of specific interactions formed by Abs with the epitope hotspot residues remains scarcely characterized. Here, using the conformational equilibrium ensembles obtained MD simulations we computed the binding free energies for the RBD-Ab complexes using the MM-GBSA method [105–108]. In this analysis, we focused on the binding free energy decomposition and examination of the energetic contributions of individual RBD epitope residues. Through this analysis, we determined the binding hotspots for Ab binding and quantified the role of the van der Waals and electrostatic interactions in the binding mechanism. The computational predictions are systematically compared against the recent experimental data on average Ab escape scores (https://github.com/jbloomlab/SARS2-RBD-escape-calc/tree/main/Cao_data/JN1-evolving-Ab-response/data/DMS/Ab). These experimental data were generated with the escape calculator [119–121] and are reported in the update analysis by Bloom’s lab (https://jbloomlab.github.io/SARS2-RBD-escape-calc/) that included the latest yeast-display DMS data by Cao and colleagues [46]. In the MM-GBSA calculations, we examine whether the binding affinities and contributions of the major binding hotspots are largely determined by the van der Waals or electrostatic interactions and whether positions of immune escape can be associated with the binding hotspots where different energetic contributions act synergistically leading to significant loss of binding upon mutations. We started with the MM-GBSA analysis of the E1 group Abs for which the residue decomposition of the total energy revealed strong and consistent binding hotspots for T345 and R346 positions (Figure 8A-C). For BD55-3546 binding, the total binding energies in these positions amounted to ΔG = −6.88 kcal/mol and ΔG = −5.48 kcal/mol (Figure 8A,D). Similar deep peaks of the binding energies for T345/R346 positions are seen for BD55-3152 (ΔG = −7.53 kcal/mol and ΔG = −11.13 kcal/mol) (Figure 8B,E) and SA58 binding (ΔG = −8.10 kcal/mol and ΔG = −7.89 kcal/mol) (Figure 8C,F). Notably, the residue decomposition also showed very favorable binding for adjacent positions N343 and A344 (Figure 8). The epitopes of these E1 Abs include the N343 glycan, which plays a critical role in modulating RBD conformation [57]. We also observed strong binding contributions of E340, L441 and V345 for BD55-3546 binding (Figure 8A,D) and corresponding significant binding components of L441, K444 and V445 for SA58 (Figure 8C,F) while for BD55-3152 the strong binding contributions were seen for L441 and K444 (Figure 8B,E). Interestingly, for SA58 Ab the tiers of strongest binding hotspots include T345/R346 in the first tier and N343, L441, K444, and V445 in the second tier (Figure 8C,F). We suggested that the positions of strongest binding hotspots should correspond to the major Ab escape positions.

**Figure 8.**
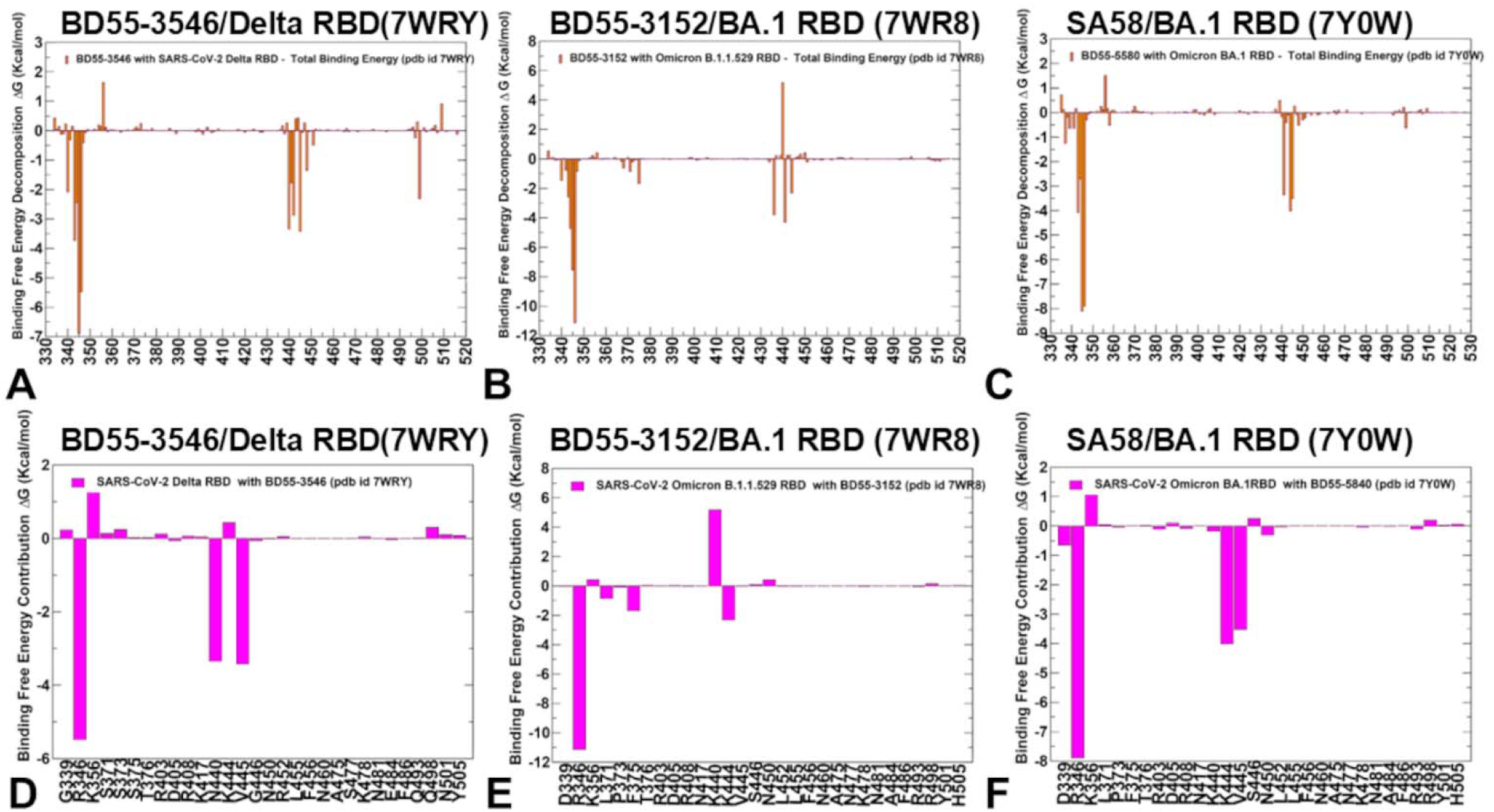
The residue-based decomposition of the binding MM-GBSA energies. (A) The residue-based decomposition of the total MM-GBSA binding energy ΔG contribution for the S-RBD complexes with E1 Abs BD55-3546 (A), BD55-3152 (B), SA58 (C). A closeup of the total MM-GBSA binding energy ΔG per residue for the sites of JN.1 mutations in the S-RBD complexes with E1 Abs BD55-3546 (D), BD55-3152 (E), SA58 (F). The MM-GBSA contributions are evaluated using 1,000 samples from the equilibrium MD simulations of respective RBD-ACE2 complexes. It is assumed that the entropy contributions for binding are similar and are not considered in the analysis. The statistical errors was estimated on the basis of the deviation between block average and are within 0.18-0.32 kcal/mol.

By comparing the predicted residue-based binding free energy contributions with the averaged Ab escape scores (Supporting Information, Figure S3), we found an excellent overall agreement between the predicted and experimental data. Indeed, as predicted the strongest immune escape sites for E1 Abs are T345 and R346 (Figure 8, Supporting Information, Figure S3). In particular, for BD55-3546, the main escape positions are E340, T345, R346 and L441 as was predicted in the energy analysis. For BD55-3142, the experiments revealed T345, R346 and L441 as major escape hotspots which is also consistent with our predictions. Finally, for SA58, the experiments singled out E340, T345, R346, L441, K444, and L452 as the dominant escape hotspots based on neutralization data against JN.1 and JN.1 subvariants (https://jbloomlab.github.io/SARS2-RBD-escape-calc/). According to the experimental data, neutralization by SA58 is moderately affected by T345N, R346Q, and R346T while strongly reduced by E340D and escaped by E340K and K444E mutations [57].

According to our computations, the top binding hotspots are E340, T345, R346, N342, K444, V445 and L441 (Figure 8C,F) showing the predictive ability of binding computations to identify key binding hotspots and associate the strongest binding centers with major escape positions. Combined with the accompanied mutational scanning of the RBD epitope residues, this strategy can represent a robust way to suggest escape sites and important escape mutations.

We also performed contribution-based energetic analysis (Figure 9) for E1 group of Abs showing strong and synergistic contributions of both van der Walls and electrostatic components for the key hotspots T345 and K346. While the electrostatic contributions are often offset by corresponding desolvation values (Figure 9, Tables S1-S3), for T345 and R346 the electrostatic interactions are strong drivers of binding and can only partly be counteracted by solvation penalties. In general, our results indicated that the dominant binding hotspots T345 and K346 for E1 Abs and their respective role as major Ab escape positions are determined by synergistic hydrophobic and electrostatic binding interactions of these residues (Figure 9). For BD55-3152, in addition to T345 and R346 hotspots, we found strong electrostatic interactions by K444 while the van der Walls component for this residue is rather small (Figure 9B,E). For this complex, E340 emerged as moderately favorable site but the electrostatic interactions of E340 with BD55-3152 are highly unfavorable (Figure 9B,E).

**Figure 9.**
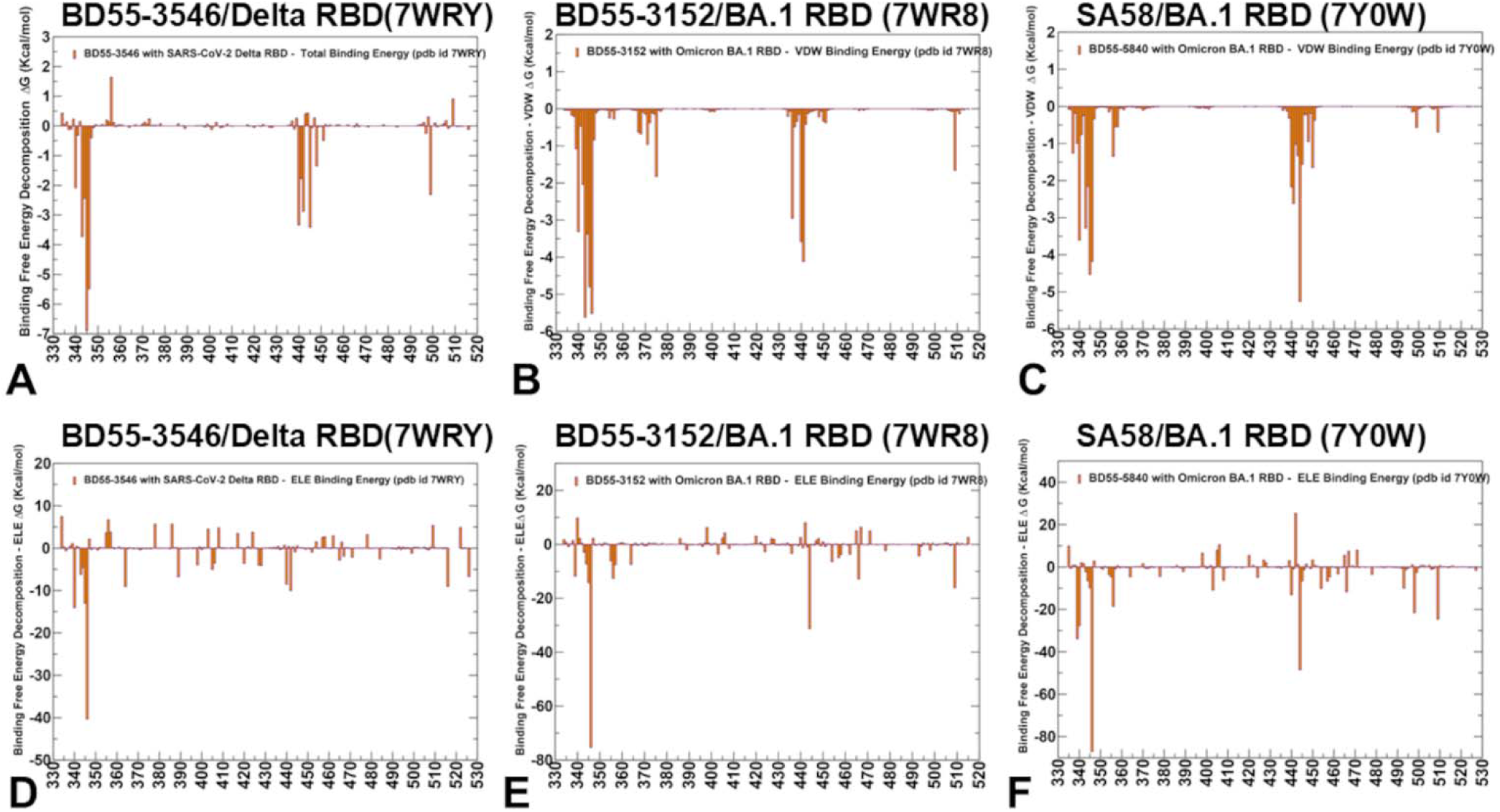
The residue-based decomposition of the binding MM-GBSA energies. (A) The residue-based decomposition of van der Walls contribution to the total MM-GBSA binding energy for the S-RBD complexes with E1 Abs BD55-3546 (A), BD55-3152 (B), SA58 (C). The residue-based decomposition of the electrostatic contribution to the total MM-GBSA binding energy for the S-RBD complexes the S-RBD complexes with E1 Abs BD55-3546 (D), BD55-3152 (E), SA58 (F). The MM-GBSA contributions are evaluated using 1,000 samples from the equilibrium MD simulations of respective RBD-ACE2 complexes. It is assumed that the entropy contributions for binding are similar and are not considered in the analysis.

A synergistic contribution of both electrostatic and hydrophobic interactions for K444 with SA58 resulted in predicting this position as a strong binding hotspot along with T345 and R346 (Figure 9C,F). As a result, most mutations in these positions can induce significant binding loss leading to the emergence of escape mechanism through convenient for virus modifications that do not compromise ACE2 binding. Indeed, SA58 showed a decreased activity due to K444N and completely escaped by the combination of K444N and T345P mutations in the experimental assays [57]. Mutations at these positions can disrupt these interactions, reducing the binding affinity of the Abs. In addition, T345 and K346 are located in regions of RBD that are crucial for maintaining its structural integrity and changes in these residues can alter the conformation of the RBD, making it more difficult for Abs to bind effectively. Interestingly, for the second tier of binding hotspots that include N343, L441, K444, and V445 residues the total binding energy is mainly determined by strong van der Waals and hydrophobic interactions while the electrostatic interactions are mostly compensated by the corresponding unfavorable desolvation contribution (Figure 9). The important result of this analysis is that MM-GBSA computations combined with mutational scanning can efficiently predict the binding hotspots and corresponding immune escape positions. Secondly, for E1 group Abs, the dominant role of T345 and K346 hotspots and immune escape centers may be due to synergistic contribution of both van der Waals and electrostatic interactions. The results also revealed that the binding of the E1 group Abs is determined by a small number of key binding energy hotspots T345, R346 and K444 while the contribution of other binding epitope positions is fairly small (Figures 8,9). Moreover, we noticed that long-range electrostatic interactions formed by the E1 group Abs are generally moderate and often unfavorable which is consistent with the notion that accumulation of positive charges on the RBD can often mediate immune resistance by positively charged S RBD-recognition surfaces [74–76]. Strikingly, however, the complementary electrostatic interactions of the E1 group Abs with T345, R346 and K444 are main determinants of binding and the emerged narrow scope of immune escape mutations (Figures 8,9).These results agree with the experiments showing that T345, K346 and K444 are involved in electrostatic interactions that are crucial for the binding of E1 Abs to the RBD [57].

The energetic drivers of F3 group of Abs are strongly determined by contributions of the RBD residues 500-508 that are also critical for RBD stability and ACE2 binding (Figure 10). For BD55-3372, the top binding hotspots are Y505, D405 and R408 (ΔG = −5.82 kcal/mol, ΔG = −4.76 kcal/mol and ΔG = −4.76 kcal/mol respectively) (Figure 10A,D). Only slightly less dominant are positions 501,502 and 503. The experiment-based Ab escape scores favored V503, G504, D405 as the main escape positions (Supporting Information, Figure S3) mostly because mutations in T500 and Y501 cannot emerge due to functional constraints even though mutations in these positions could significantly diminish BD55-3372 binding (Figure 10). Y508 is also essential to ACE2 binding and could hardly be evolved [57]. BD55-3372 is sensitive to mutations of V503 and G504 that are less critical for ACE2 binding but are still constrained for evolution due to role of these sites in RBD critical functions.

**Figure 10.**
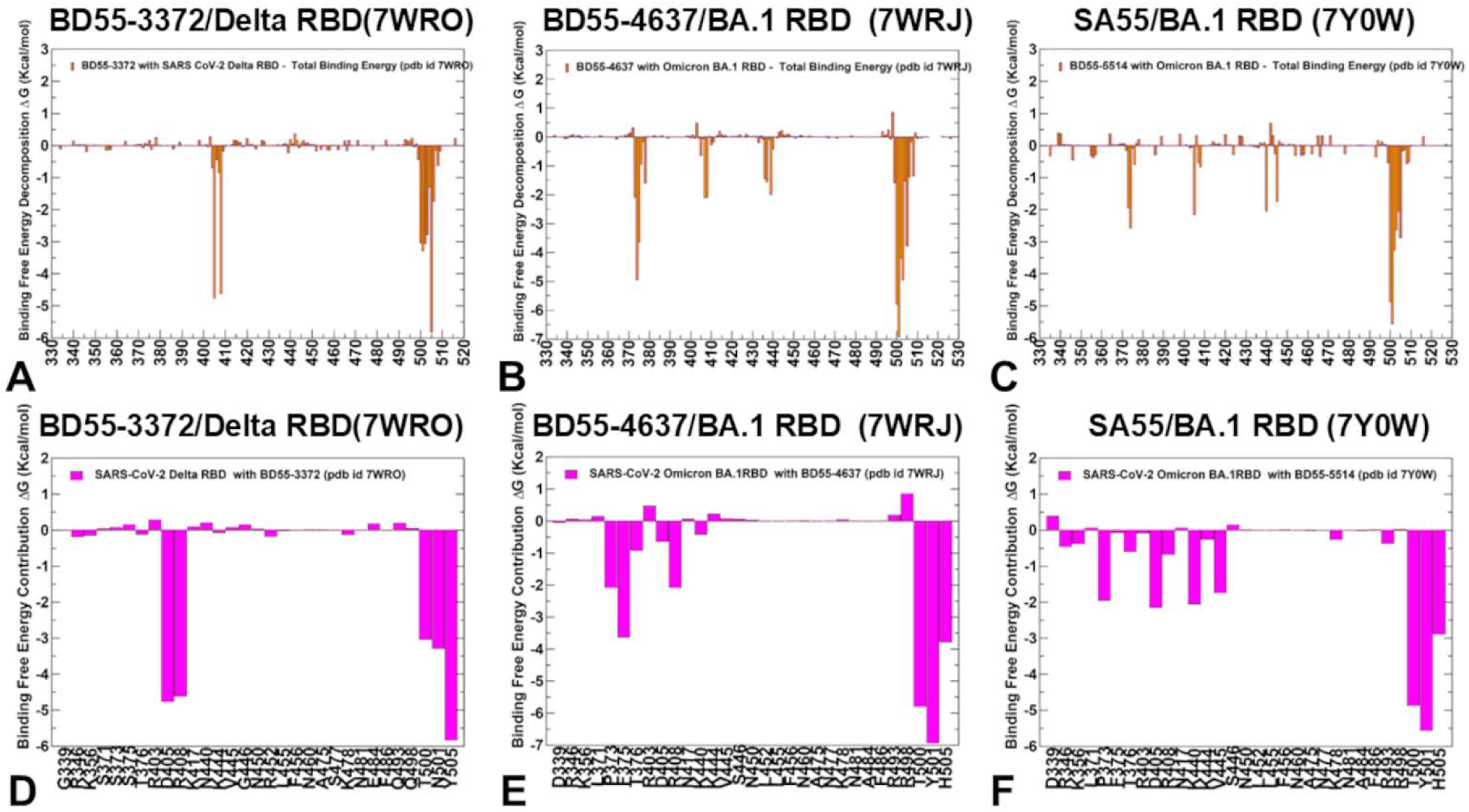
The residue-based decomposition of the binding MM-GBSA energies. (A) The residue-based decomposition of the total MM-GBSA binding energy ΔG contribution for the S-RBD complexes with F3 group Abs : BD55-3372 (A), BD55-4637 (B) and SA55 (C). A closeup of the total MM-GBSA binding energy ΔG per residue for the sites of JN.1 mutations in the S-RBD complexes with F3 group Abs : BD55-3372 (D), BD55-4637 (E) and SA55 (F). The MM-GBSA contributions are evaluated using 1,000 samples from the equilibrium MD simulations of respective RBD-ACE2 complexes. It is assumed that the entropy contributions for binding are similar and are not considered in the analysis. The statistical errors was estimated on the basis of the deviation between block average and are within 0.16-0.45 kcal/mol.

For BD55-4637, the energy contributions revealed major binding hotspots in positions Y501, T500, V503, F374, G502 and H505 (Figure 10B,E). Importantly, these sites and particularly Y501, T500 are fundamental for ACE2 binding function, while F374, G502 and V503 are also involved in the RBD stability and ACE2 interactions. The second tier of binding hotspots included F375, R408, N439 and K478 (Figure 10B,E). The experiments showed sites Y508, G502, V503, N439 and F378 as major escape centers (Supporting Information, Figure S3) thus revealing good general agreement with the binding energy computations. Similarly, for SA55 our result predicted Y501, T500, G502, H505, V503, G504, F374, D405, K440, and V445 positions as important RBD positions for binding, showing an expanded set of binding hotspots but still predicting region 500-505 as the dominant for binding (Figure 10C,F). The positions of immune escape for SA55 are G502, H505, V503, G504, Y508 (Supporting Information, Figure S3). The efficacy of SA55 is affected by Y508H, moderately affected by G504S, strongly affected by K440E, and escaped by V503E and G504D [57]. Hence, our computations correctly predict the most critical escape centers for SA55.

We followed with the analysis of individual energetic contributions for F3 group Abs revealing the major role of the van der Waals interaction component (Figure 11). This is consistent with the hydrophobic nature of the critical binding hotspot region (residues 500-505), showing the same overall shape and relative residue contributions for the total energy (Figure 10) and van der Waals interaction component (Figure 11). For BD55-3372, the favorable electrostatic interactions are formed by R403, R408 and K444 sites but these contributions are often offset by desolvation penalties with the exception of R408 (Figure 11 D). For BD55-4637, the electrostatics drives binding energy for K378 and R408 (Figure 11E). For SA55, the electrostatic component is the dominant force behind energetics of the binding hotspots K378, K440, R403, R408 and K444 (Figure 11F). For these F3 Abs the electrostatic interactions with the RBD residues with F3 Abs include both favorable and highly unfavorable contributions (Figure 11D-F). For these Abs, we also observed that some secondary tier hotspots emerge when both van der Waals, and electrostatic contributions act synergistically to mediate strong binding. Indeed, for BD55-3372, the favorable electrostatic by R403 and R408 are synergistic with the favorable hydrophobic and van der Waals contributions (Figure 11A,D). The strong electrostatic interactions made by K378 in the complex with BD55-4637 (Figure 11E) and by K440 in the complex with SA55 (Figure 11F) are the determining factors for these hotspots which is consistent with the experiments showing that SA55 binding can be significantly compromised by changes in the charge by K440E mutation [57]. K378 residue is less commonly targeted by specific Abs, but it can be involved in the binding sites of broadly neutralizing Abs that recognize conserved regions of the RBD. K444 is more frequently targeted by neutralizing Abs including group D Abs, such as REGN-10987, LY-CoV1404 and COV2-2130, that bind to the linear epitope 440–449 and showed effectiveness in neutralizing various Omicron variants [54–58]. N440K mutants along with K444N/T were resistant to C135 Ab which is class 3 Ab targeting the same binding epitope as F3 group Abs [122].

**Figure 11.**
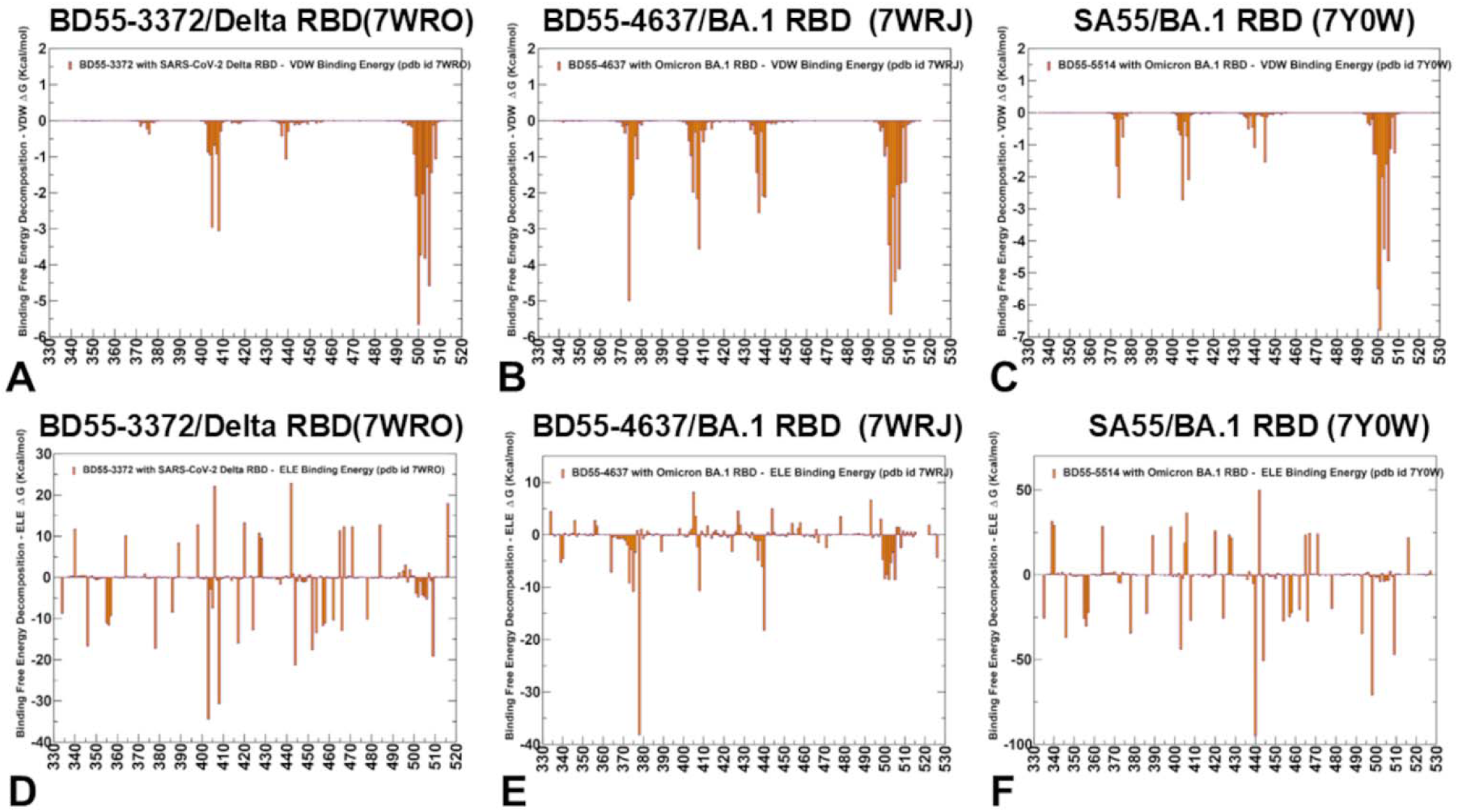
The residue-based decomposition of the binding MM-GBSA energies. (A) The residue-based decomposition of van der Walls contribution to the total MM-GBSA binding energy for the S-RBD complexes with F3 group Abs : BD55-3372 (A), BD55-4637 (B) and SA55 (C). The residue-based decomposition of the electrostatic contribution to the total MM-GBSA binding energy for the S-RBD complexes the S-RBD complexes with F3 group Abs : BD55-3372 (D), BD55-4637 (E) and SA55 (F).

## 4. Discussion

The SARS-CoV-2 virus must maintain a delicate balance between binding effectively to the ACE2 receptor and evading Abs. Broadly neutralizing monoclonal Abs recognize conserved antigenic sites outside the RBM as Omicron-mediated immune evasion marked a significant antigenic shift in SARS-CoV-2 [46, 123]. The recent evidence indicated that the latest variants and JN.1 sublineages can display lower affinity to ACE2 and the enhanced antibody evasion profile [46,123]. Mutations that shift the electrostatic potential can help the virus achieve this balance between ACE2 binding and immune escape. In general, the emergence of the JN.1 descendants accompanied by the increased Ab evasion and decrease in ACE2 affinity suggested the evolutionary objectives of the virus. In this context, it is important to understand the energetic mechanisms underlying the evolving Ab response to Omicron antigenic shift to JN.1 by which F3 Abs retain potency and neutralizing activity against JN.1 subvariants, whereas Abs from other groups largely escaped [46, 54–58].

Mutational scanning and binding free energy calculations provide insights why E1 and F3 Abs targeting more conserved epitope surfaces, particularly F3, are effective and exhibit broadly neutralizing activity against latest variants. The results showed that binding E1 Abs is strongly susceptible to changes in charges and electrostatic interactions of T345 and R346 which are the dominant binding hotspots and immune escape centers. As a result, mutations R346Q and R346T emerged as important Ab escape changes for this group [57]. On the other hand, binding F3 Abs is driven primarily by hydrophobic interactions that are formed with the RBD residues 500-508 implicated in stability, ACE2 binding and critical spike functions. Our results provide an interesting rationale to the experimental observation that mutations in the epitope consisting of sites R403, D405, V503, G504, and H505 can cause significant escape from binding by neutralizing Abs targeting the RBD and yet these mutations have not emerged in circulation [46,56,58]. We suggest that the evolution of the RBD mutations in these positions may be constrained due to synergistic energetic couplings between T500/Y501 that are indispensable for ACE2 binding and structurally proximal V503, G504 and H505 sites. According to our results, binding energetics for E1 and F3 groups of Abs is driven by a small number of key binding hotspots that can also include positively charged residues R346, K378, R403, R408, K440 and K444 forming strong electrostatic interactions. In particular, for F3 Abs we found that the electrostatic interactions are determining factors for the binding hotspots K378, K440, R403, R408 and K444. We previously showed that mutational sites that contribute to the ACE2-binding affinity can include RBD residues K378, R403, K424, K440, K444, K460, N477, K478 owing to electrostatic interactions mediated by lysine residues [66, 81,82]. Concurrently, the results of this study suggested that F3 group Abs may achieve broad neutralizing activity against JN.1 variants by targeting the specific charged RBD sites that could provide strong Ab binding but are also important for ACE2 binding. The observed immune escape mutations in these positions K444T/N, K440T/E are detrimental to Ab binding [57] by reducing the positive electrostatic potential on the RBD and possibly the conformation of the RBD which may also adversely affect RBD binding to the ACE2. Another functional study highlighted the expected disruptions of electrostatic interactions with the Abs resulting from the K444T and the R346T RBD mutations that can boost immune escape profile [124]. Based on our analysis, we argue that the broadly neutralizing activity of E1 and F3 Abs against the latest variants may also be partly attributed to the narrowed repertoire of immune escape mutations on the RBD to combat binding of these Abs where some of these mutations are forced to reduce the positive charge on the RBD.

It is also worth noting that although the NTD mutational change make the charge more negative [125] the overall charge on the S protein has been evolving towards progressively higher values. However, recently emerged lineages and escape mutants showed a greater diversity in the composition of ionizable amino acids. Recent analysis found that a progressive increase in the positive charge on the S protein may have reached a peak with XBB lineages in which the total charge of the S protein is still positive but shows a greater variability, while the JN.1 and its subvariants still displayed the increasing number of positively charged residues. [126]. Our analysis suggested that these positively charged positions in JN.1 subvariants can be successfully exploited by E1 and F3 Abs forcing the virus to alter the charge on the escape centers. According to our results, E1 and F3 groups of Abs effectively exploit binding hotspot clusters of hydrophobic sites critical for RBD functions along with selective complementary targeting of positively charged RBD sites that are important for ACE2 binding. Together with targeting conserved epitopes, these groups of Abs can lead to the expanded neutralization breadth and resilience to antigenic shift associated with viral evolution. These findings suggest that RBD mutations and their associated immune escape may have been reaching a certain plateau level under the constraint of the RBD stability and ACE2 binding and that selective pressures may explore epistatic effects and localized electrostatic changes to boost immune evasion.

## 5. Conclusions

Using dynamic ensembles of the S-Ab complexes and systematic mutational scanning of the RBD residues for binding with Abs, we characterize patterns of mutational sensitivity and compute mutational scanning heatmaps to identify escape hotspot centers for a panel of E1 and F3 Abs. Mutational scanning using rapid computation of binding free energy changes revealed binding hotspots for E1 and F3 Abs that are consistent with the experimental DMS data and immune escape centers. In particular, consistent with the experiments [57], mutational scanning predicted T345 and R346 residues as dominant binding hotspots for E1 group of Abs. Similarly, our results predicted RBD sites V503, G504 and Y508 as the key binding hotspots for F3 group of Abs that agrees with the experiments showing that these positions are major immune escape centers for these Abs. MM-GBSA binding free energy computations revealed group-specific binding energy hotspots that are consistent with the experimentally determined immune escape centers. Our analysis suggested that E1 and F3 groups of Abs targeting binding epitopes may exploit strong hydrophobic interactions with the binding epitope hotspots critical for the RBD stability and ACE2 binding, while escape mutations tend to emerge in sites associated with synergistically strong van der Waals and electrostatic interactions. Our analysis shows that the emergence of a small number of immune escape positions for E1 group Abs are associated with R346 and K444 positions. The binding of F3 group Abs yields Ab-specific balance of the van der Waals and electrostatic contributions. Our analysis confirms that V503, G504 and H505 correspond to the key escape positions for F3 Abs as T500 and Y501 are critical for ACE2 binding function. The second strongest group of binding hotspots for F3 Abs is R403 and D405 in which mutations are typically tolerant to the ACE2 affinity. In addition, despite the common binding epitope, different F3 Abs can also have unique binding hotspots in RBD positions F374, T376, and K378. These results are consistent with the experimental data on the strongest RBD sites of immune escape from binding by neutralizing Abs.

Recent studies of emerging Omicron variants suggested that the evolutionary paths for significant improvements in the binding affinity of the Omicron RBD variants with ACE2 are relatively narrow and may involve convergent mutational hotspots to primarily optimize immune escape while retaining sufficient ACE2 affinity. These mechanisms based on convergent adaptation may determine the scope of “ evolutionary opportunities” for the virus to adapt new mutations that improve immune resistance without compromising ACE2 binding affinity and stability. Our results may be helpful in rationalizing the roles and synergistic contributions of hydrophobic and electrostatic drivers of binding for broadly neutralizing groups of Abs by exploiting electrostatic complementarity to a targeted set of positively charged RBD residues and binding to hydrophobic patches of functionally important RBD positions on conserved epitopes.

## Supplementary Materials

Figure S1 highlights the evolution of SARSB-CoV-2 lineages. Figure S2 represents the prevalence of the JN.1 variant among all the samples in the world tested. Figure S3 presents the residue-based averaged Ab escape scores derived from the latest experimental data [46]. Table S1 lists the binding epitope residues and contacts with the RBD for BD55-3546. Table S2 lists the binding epitope residues and contacts with the RBD for BD55-3152. Table S3 lists the binding epitope residues and contacts with the RBD for SA58. Table S4 lists the binding epitope residues and contacts with the RBD for BD55-3372. Table S5 lists the binding epitope residues and contacts with the RBD for BD55-4637. Table S6 lists the binding epitope residues and contacts with the RBD for SA55.

## Author Contributions

Conceptualization, G.V.; Methodology, N.R., M.A., V.P., B.F., G.V.; Software, N.R., M.A., V.P., B.F., G.V.; Validation, N.R., G.V.; Formal analysis, N.R., G.V., M.A., V.P.,B.F.; Investigation, N.R., G.V.; Resources, N.R., G.V., M.A, G.V.; Data curation, N.R., M.A., G.C., G.V.; Writing—original draft preparation, N.R., G.V.; Writing—review and editing, G.V.; Visualization, N.R., G.V. Supervision G.V. Project administration, G.V.; Funding acquisition, G.V. All authors have read and agreed to the published version of the manuscript.

## Data Availability Statement

The original contributions presented in this study are included in the article/supplementary material. Crystal structures were obtained and downloaded from the Protein Data Bank (http://www.rcsb.org). All simulations were performed using NAMD 2.13 package that was obtained from website https://www.ks.uiuc.edu/Development/Download/. All simulations were performed using the all-atom additive CHARMM36 protein force field that can be obtained from http://mackerell.umaryland.edu/charmm_ff.shtml. The rendering of protein structures was done with interactive visualization program UCSF ChimeraX package (https://www.rbvi.ucsf.edu/chimerax/) and Pymol (https://pymol.org/2/).

## Acknowledgments

The authors acknowledge support from Schmid College of Science and Technology at Chapman University for providing computing resources at the Keck Center for Science and Engineering.

## References

1. Tai, W.; He, L.; Zhang, X.; Pu, J.; Voronin, D.; Jiang, S.; Zhou, Y.; Du, L. Characterization of the receptor-binding domain (RBD) of 2019 novel coronavirus: implication for development of RBD protein as a viral attachment inhibitor and vaccine. Cell. Mol. Immunol. 2020, 17, 613–620. doi: 10.1038/s41423-020-0400-4.

2. Wang, Q.; Zhang, Y.; Wu, L.; Niu, S.; Song, C.; Zhang, Z.; Lu, G.; Qiao, C.; Hu, Y.; Yuen, K. Y.; Wang, Q.; Zhou, H.; Yan, J.; Qi, J. Structural and functional basis of SARS-CoV-2 entry by using human ACE2. Cell 2020, 181, 894–904.e9. doi: 10.1016/j.cell.2020.03.045.

3. Walls, A. C.; Park, Y. J.; Tortorici, M. A.; Wall, A.; McGuire, A. T.; Veesler, D. Structure, Function, and Antigenicity of the SARS-CoV-2 Spike Glycoprotein. Cell 2020, 181, 281–292.e6. doi: 10.1016/j.cell.2020.02.058.

4. Wrapp, D.; Wang, N.; Corbett, K. S.; Goldsmith, J. A.; Hsieh, C. L.; Abiona, O.; Graham, B. S.; McLellan, J. S. Cryo-EM structure of the 2019-nCoV spike in the prefusion conformation. Science 2020, 367, 1260–1263. doi: 10.1126/science.abb2507.

5. Cai, Y.; Zhang, J.; Xiao, T.; Peng, H.; Sterling, S. M.; Walsh, R. M., Jr.; Rawson, S.; Rits-Volloch, S.; Chen, B. Distinct conformational states of SARS-CoV-2 spike protein. Science 2020, 369, 1586–1592. doi: 10.1126/science.abd4251.

6. Hsieh, C. L.; Goldsmith, J. A.; Schaub, J. M.; DiVenere, A. M.; Kuo, H. C.; Javanmardi, K.; Le, K. C.; Wrapp, D.; Lee, A. G.; Liu, Y., Chou, C.W.; Byrne, P.O.; Hjorth, C.K.; Johnson, N.V.; Ludes-Meyers J.; Nguyen, A.W.; Park, J.; Wang, N.; Amengor, D.; Lavinder, J.J.; Ippolito, G.C.; Maynard, J.A.; Finkelstein, I.J.; McLellan, J.S. Structure-based design of prefusion-stabilized SARS-CoV-2 spikes. Science 2020, 369, 1501–1505. doi: 10.1126/science.abd0826.

7. Henderson, R.; Edwards, R. J.; Mansouri, K.; Janowska, K.; Stalls, V.; Gobeil, S. M. C.; Kopp, M.; Li, D.; Parks, R.; Hsu, A. L., Borgnia, M.J.; Haynes, B.F.; Acharya, P. Controlling the SARS-CoV-2 spike glycoprotein conformation. Nat. Struct. Mol. Biol. 2020, 27, 925–933. doi: 10.1038/s41594-020-0479-4.

8. McCallum, M.; Walls, A. C.; Bowen, J. E.; Corti, D.; Veesler, D. Structure-guided covalent stabilization of coronavirus spike glycoprotein trimers in the closed conformation. Nat. Struct. Mol. Biol. 2020, 27, 942–949. doi: 10.1038/s41594-020-0483-8.

9. Xiong, X.; Qu, K.; Ciazynska, K. A.; Hosmillo, M.; Carter, A. P.; Ebrahimi, S.; Ke, Z.; Scheres, S. H. W.; Bergamaschi, L.; Grice, G. L., Zhang, Y.; CITIID-NIHR COVID-19 BioResource Collaboration, Nathan, J.A.; Baker, S.; James, L.C.; Baxendale, H.E.; Goodfellow, I.; Doffinger, R.; Briggs, J.A.G. A thermostable, closed SARS-CoV-2 spike protein trimer. Nat. Struct. Mol. Biol. 2020, 27, 934–941. doi: 10.1038/s41594-020-0478-5.

10. Costello, S.M.; Shoemaker, S.R.; Hobbs, H.T.; Nguyen, A.W.; Hsieh, C.L.; Maynard, J.A.; McLellan, J.S.; Pak, J.E.; Marqusee, S. The SARS-CoV-2 spike reversibly samples an open-trimer conformation exposing novel epitopes. Nat. Struct. Mol. Biol. 2022, 27, 229–238. doi: 10.1038/s41594-022-00735-5.

11. McCormick, K.D.; Jacobs, J.L.; Mellors, J.W. The emerging plasticity of SARS-CoV-2. Science 2021, 371, 1306–1308. doi: 10.1126/science.abg4493.

12. Ghimire, D.; Han, Y.; Lu, M. Structural Plasticity and Immune Evasion of SARS-CoV-2 Spike Variants. Viruses 2022, 14, 1255. 10.3390/v14061255.

13. Xu, C.; Wang, Y.; Liu, C.; Zhang, C.; Han, W.; Hong, X.; Wang, Y.; Hong, Q.; Wang, S.; Zhao, Q.; Wang, Y.; Yang, Y.; Chen, K.; Zheng, W.; Kong, L.; Wang, F.; Zuo, Q.; Huang, Z.; Cong, Y. Conformational dynamics of SARS-CoV-2 trimeric spike glycoprotein in complex with receptor ACE2 revealed by cryo-EM. Sci. Adv. 2021, 7, eabe5575. doi: 10.1126/sciadv.abe5575.

14. Benton, D. J.; Wrobel, A. G.; Xu, P.; Roustan, C.; Martin, S. R.; Rosenthal, P. B.; Skehel, J. J.; Gamblin, S. J. Receptor binding and priming of the spike protein of SARS-CoV-2 for membrane fusion. Nature 2020, 588, 327–330. doi: 10.1038/s41586-020-2772-0.

15. Turoňová, B.; Sikora, M.; Schuerman, C.; Hagen, W. J. H.; Welsch, S.; Blanc, F. E. C.; von Bülow, S.; Gecht, M.; Bagola, K.; Hörner, C.; van Zandbergen, G.; Landry, J.; de Azevedo, N. T. D.; Mosalaganti, S.; Schwarz, A.; Covino, R.; Mühlebach, M. D.; Hummer, G.; Krijnse Locker, J.; Beck, M. In situ structural analysis of SARS-CoV-2 spike reveals flexibility mediated by three hinges. Science 2020, 370, 203–208. doi: 10.1126/science.abd5223.

16. Lu, M.; Uchil, P. D.; Li, W.; Zheng, D.; Terry, D. S.; Gorman, J.; Shi, W.; Zhang, B.; Zhou, T.; Ding, S.; Gasser, R.; Prevost, J.; Beaudoin-Bussieres, G.; Anand, S. P.; Laumaea, A.; Grover, J. R.; Lihong, L.; Ho, D. D.; Mascola, J.R.; Finzi, A.; Kwong, P. D.; Blanchard, S. C.; Mothes, W. Real-time conformational dynamics of SARS-CoV-2 spikes on virus particles. Cell Host Microbe. 2020, 28, 880–891.e8. doi: 10.1016/j.chom.2020.11.001.

17. Yang, Z.; Han, Y.; Ding, S.; Shi, W.; Zhou, T.; Finzi, A.; Kwong, P.D.; Mothes, W.; Lu, M. SARS-CoV-2 Variants Increase Kinetic Stability of Open Spike Conformations as an Evolutionary Strategy. mBio 2022, 13, e0322721. doi: 10.1128/mbio.03227-21.

18. Díaz-Salinas, M.A.; Li, Q.; Ejemel, M.; Yurkovetskiy, L.; Luban, J.; Shen, K.; Wang, Y.; Munro, J.B. Conformational dynamics and allosteric modulation of the SARS-CoV-2 spike. Elife 2022, 11, e75433. doi: 10.7554/eLife.75433.

19. Wang, Y.; Liu, C.; Zhang, C.; Wang, Y.; Hong, Q.; Xu, S.; Li, Z.; Yang, Y.; Huang, Z.; Cong, Y. Structural Basis for SARS-CoV-2 Delta Variant Recognition of ACE2 Receptor and Broadly Neutralizing Antibodies. Nat. Commun. 2022, 13, 871. doi: 10.1038/s41467-022-28528-w.

20. Mannar, D.; Saville, J.W.; Zhu, X.; Srivastava, S.S.; Berezuk, A.M.; Tuttle, K.S.; Marquez, A.C.; Sekirov, I.; Subramaniam, S. SARS-CoV-2 Omicron Variant: Ab Evasion and Cryo-EM Structure of Spike Protein–ACE2 Complex. Science 2022, 375, 760–764. doi: 10.1126/science.abn7760.

21. Hong, Q.; Han, W.; Li, J.; Xu, S.; Wang, Y.; Xu, C.; Li, Z.; Wang, Y.; Zhang, C.; Huang, Z.; Cong, Y. Molecular Basis of Receptor Binding and Ab Neutralization of Omicron. Nature 2022. doi: 10.1038/s41586-022-04581-9.

22. McCallum, M.; Czudnochowski, N.; Rosen, L.E.; Zepeda, S.K.; Bowen, J.E.; Walls, A.C.; Hauser, K.; Joshi, A.; Stewart, C.; Dillen, J.R.; Powell, A.E.; Croll, T.I.; Nix, J.; Virgin, H.W.; Corti, D.; Snell, G.; Veesler, D. Structural Basis of SARS-CoV-2 Omicron Immune Evasion and Receptor Engagement. Science 2022, 375, 864–868. doi: 10.1126/science.abn8652.

23. Yin, W.; Xu, Y.; Xu, P.; Cao, X.; Wu, C.; Gu, C.; He, X.; Wang, X.; Huang, S.; Yuan, Q.; Wu, K.; Hu, W.; Huang, Z.; Liu, J.; Wang, Z.; Jia, F.; Xia, K.; Liu, P.; Wang, X.; Song, B.; Zheng, J.; Jiang, H.; Cheng, X.; Jiang, Y.; Deng, S.J.; Xu, H.E. Structures of the Omicron Spike Trimer with ACE2 and an Anti-Omicron Ab. Science 2022, 375, 1048–1053. doi: 10.1126/science.abn8863.

24. Gobeil, S. M.-C.; Henderson, R.; Stalls, V.; Janowska, K.; Huang, X.; May, A.; Speakman, M.; Beaudoin, E.; Manne, K.; Li, D.; Parks, R.; Barr, M.; Deyton, M.; Martin, M.; Mansouri, K.; Edwards, R. J.; Eaton, A.; Montefiori, D. C.; Sempowski, G. D.; Saunders, K. O.; Wiehe, K.; Williams, W.; Korber, B.; Haynes, B. F.; Acharya, P. Structural Diversity of the SARS-CoV-2 Omicron Spike. Mol Cell. 2022, 82, 2050–2068.e6. doi: 10.1016/j.molcel.2022.03.028.

25. Cui, Z.; Liu, P.; Wang, N.; Wang, L.; Fan, K.; Zhu, Q.; Wang, K.; Chen, R.; Feng, R.; Jia, Z.; Yang, M.; Xu, G.; Zhu, B.; Fu, W.; Chu, T.; Feng, L.; Wang, Y.; Pei, X.; Yang, P.; Xie, X.S.; Cao, L.; Cao, Y.; Wang, X. Structural and Functional Characterizations of Infectivity and Immune Evasion of SARS-CoV-2 Omicron. Cell 2022, 185, 860–871.e13. doi: 10.1016/j.cell.2022.01.019.

26. Parums DV. Editorial: The XBB.1.5 (’Kraken’) Subvariant of Omicron SARS-CoV-2 and its Rapid Global Spread. Med Sci Monit. 2023, 29, e939580. doi: 10.12659/MSM.939580.

27. Wang, Q.; Iketani, S.; Li, Z.; Liu, L.; Guo, Y.; Huang, Y.; Bowen, A. D.; Liu, M.; Wang, M.; Yu, J.; Valdez, R.; Lauring, A. S.; Sheng, Z.; Wang, H. H.; Gordon, A.; Liu, L.; Ho, D. D. Alarming Ab Evasion Properties of Rising SARS-CoV-2 BQ and XBB Subvariants. Cell 2023, 186, 279–286.e8. 10.1016/j.cell.2022.12.018.

28. Hoffmann, M.; Arora, P.; Nehlmeier, I.; Kempf, A.; Cossmann, A.; Schulz, S. R.; Morillas Ramos, G.; Manthey, L. A.; Jäck, H.-M.; Behrens, G. M. N.; Pöhlmann, S. Profound Neutralization Evasion and Augmented Host Cell Entry Are Hallmarks of the Fast-Spreading SARS-CoV-2 Lineage XBB.1.5. Cell Mol Immunol. 2023, 1–4. doi: 10.1038/s41423-023-00988-0.

29. Yamasoba, D.; Uriu, K.; Plianchaisuk, A.; Kosugi, Y.; Pan, L.; Zahradnik, J.; Ito, J.; Sato, K. Virological Characteristics of the SARS-CoV-2 Omicron XBB.1.16 Variant. Lancet Infect Dis. 2023, S1473-3099(23)00278-5. doi: 10.1016/S1473-3099(23)00278-5.

30. Tsujino, S.; Deguchi, S.; Nomai, T.; Padilla-Blanco, M.; Plianchaisuk, A.; Wang, L.; Begum, M. M.; Uriu, K.; Mizuma, K.; Nao, N.; Kojima, I.; Tsubo, T.; Li, J.; Matsumura, Y.; Nagao, M.; Oda, Y.; Tsuda, M.; Anraku, Y.; Kita, S.; Yajima, H.; Sasaki-Tabata, K.; Guo, Z.; Hinay, A. A., Jr.; Yoshimatsu, K.; Yamamoto, Y.; Nagamoto, T.; Asakura, H.; Nagashima, M.; Sadamasu, K.; Yoshimura, K.; Nasser, H.; Jonathan, M.; Putri, O.; Kim, Y.; Chen, L.; Suzuki, R.; Tamura, T.; Maenaka, K.; Irie, T.; Matsuno, K.; Tanaka, S.; Ito, J.; Ikeda, T.; Takayama, K.; Zahradnik, J.; Hashiguchi, T.; Fukuhara, T.; Sato, K. Virological Characteristics of the SARS-CoV-2 Omicron EG.5.1 Variant. bioRxiv 2023. 10.1101/2023.10.19.563209.

31. Wang, Q.; Guo, Y.; Zhang, R. M.; Ho, J.; Mohri, H.; Valdez, R.; Manthei, D. M.; Gordon, A.; Liu, L.; Ho, D. D. Ab Neutralization of Emerging SARS-CoV-2 Subvariants: EG.5.1 and XBC.1.6. Lancet Infect Dis. 2023, 23, e397–e398. doi: 10.1016/S1473-3099(23)00555-8.

32. Faraone, J. N.; Qu, P.; Goodarzi, N.; Zheng, Y.-M.; Carlin, C.; Saif, L. J.; Oltz, E. M.; Xu, K.; Jones, D.; Gumina, R. J.; Liu, S.-L. Immune Evasion and Membrane Fusion of SARS-CoV-2 XBB Subvariants EG.5.1 and XBB.2.3. Emerg. Microbes Infect. 2023, 12, 2270069. doi: 10.1080/22221751.2023.2270069.

33. Kosugi, Y.; Plianchaisuk, A.; Putri, O.; Uriu, K.; Kaku, Y.; Hinay, A. A., Jr; Chen, L.; Kuramochi, J.; Sadamasu, K.; Yoshimura, K.; Asakura, H.; Nagashima, M.; Ito, J.; Sato, K.; Misawa, N.; Guo, Z.; Tolentino, J. E. M.; Fujita, S.; Pan, L.; Suganami, M.; Chiba, M.; Yoshimura, R.; Yasuda, K.; Iida, K.; Ohsumi, N.; Strange, A. P.; Tanaka, S.; Fukuhara, T.; Tamura, T.; Suzuki, R.; Suzuki, S.; Ito, H.; Matsuno, K.; Sawa, H.; Nao, N.; Tanaka, S.; Tsuda, M.; Wang, L.; Oda, Y.; Ferdous, Z.; Shishido, K.; Nakagawa, S.; Shirakawa, K.; Takaori-Kondo, A.; Nagata, K.; Nomura, R.; Horisawa, Y.; Tashiro, Y.; Kawai, Y.; Takayama, K.; Hashimoto, R.; Deguchi, S.; Watanabe, Y.; Sakamoto, A.; Yasuhara, N.; Hashiguchi, T.; Suzuki, T.; Kimura, K.; Sasaki, J.; Nakajima, Y.; Yajima, H.; Irie, T.; Kawabata, R.; Tabata, K.; Ikeda, T.; Nasser, H.; Shimizu, R.; Begum, M. M.; Jonathan, M.; Mugita, Y.; Takahashi, O.; Ichihara, K.; Ueno, T.; Motozono, C.; Toyoda, M.; Saito, A.; Shofa, M.; Shibatani, Y.; Nishiuchi, T. Characteristics of the SARS-CoV-2 Omicron HK.3 Variant Harbouring the FLip Substitution. Lancet Microbe 2024, S2666-5247(23)00373-7. doi: 10.1016/S2666-5247(23)00373-7.

34. Wang, Q.; Guo, Y.; Liu, L.; Schwanz, L. T.; Li, Z.; Nair, M. S.; Ho, J.; Zhang, R. M.; Iketani, S.; Yu, J.; Huang, Y.; Qu, Y.; Valdez, R.; Lauring, A. S.; Huang, Y.; Gordon, A.; Wang, H. H.; Liu, L.; Ho, D. D. Antigenicity and Receptor Affinity of SARS-CoV-2 BA.2.86 Spike. Nature 2023. 10.1038/s41586-023-06750-w.

35. Yang, S.; Yu, Y.; Jian, F.; Song, W.; Yisimayi, A.; Chen, X.; Xu, Y.; Wang, P.; Wang, J.; Yu, L.; Niu, X.; Wang, J.; Xiao, T.; An, R.; Wang, Y.; Gu, Q.; Shao, F.; Jin, R.; Shen, Z.; Wang, Y.; Cao, Y. Antigenicity and Infectivity Characterization of SARS-CoV-2 BA.2.86. Lancet Infect Dis. 2023, 23, e457–e459. doi: 10.1016/S1473-3099(23)00573-X.

36. Tamura, T.; Mizuma, K.; Nasser, H.; Deguchi, S.; Padilla-Blanco, M.; Oda, Y.; Uriu, K.; Tolentino, J. E. M.; Tsujino, S.; Suzuki, R.; Kojima, I.; Nao, N.; Shimizu, R.; Wang, L.; Tsuda, M.; Jonathan, M.; Kosugi, Y.; Guo, Z.; Hinay, A. A., Jr.; Putri, O.; Kim, Y.; Tanaka, Y. L.; Asakura, H.; Nagashima, M.; Sadamasu, K.; Yoshimura, K.; Saito, A.; Ito, J.; Irie, T.; Tanaka, S.; Zahradnik, J.; Ikeda, T.; Takayama, K.; Matsuno, K.; Fukuhara, T.; Sato, K. Virological Characteristics of the SARS-CoV-2 BA.2.86 Variant. Cell Host Microbe. 2024, 32, 170–180.e12. doi: 10.1016/j.chom.2024.01.001.

37. Liu, C.; Zhou, D.; Dijokaite-Guraliuc, A.; Supasa, P.; Duyvesteyn, H. M. E.; Ginn, H. M.; Selvaraj, M.; Mentzer, A. J.; Das, R.; de Silva, T. I.; Ritter, T. G.; Plowright, M.; Newman, T. A. H.; Stafford, L.; Kronsteiner, B.; Temperton, N.; Lui, Y.; Fellermeyer, M.; Goulder, P.; Klenerman, P.; Dunachie, S. J.; Barton, M. I.; Kutuzov, M. A.; Dushek, O.; Fry, E. E.; Mongkolsapaya, J.; Ren, J.; Stuart, D. I.; Screaton, G. R. A Structure-Function Analysis SARS-CoV-2 BA.2.86 Balances Ab Escape and ACE2 Affinity. Cell Rep Med. 2024, 5, 101553. doi: 10.1016/j.xcrm.2024.101553.

38. Khan, K.; Lustig, G.; Römer, C.; Reedoy, K.; Jule, Z.; Karim, F.; Ganga, Y.; Bernstein, M.; Baig, Z.; Jackson, L.; Mahlangu, B.; Mnguni, A.; Nzimande, A.; Stock, N.; Kekana, D.; Ntozini, B.; van Deventer, C.; Marshall, T.; Manickchund, N.; Gosnell, B. I.; Lessells, R. J.; Karim, Q. A.; Abdool Karim, S. S.; Moosa, M.-Y. S.; de Oliveira, T.; von Gottberg, A.; Wolter, N.; Neher, R. A.; Sigal, A. Evolution and Neutralization Escape of the SARS-CoV-2 BA.2.86 Subvariant. Nat Commun. 2023, 14, 8078. doi: 10.1038/s41467-023-43703-3.

39. Yang, S.; Yu, Y.; Xu, Y.; Jian, F.; Song, W.; Yisimayi, A.; Wang, P.; Wang, J.; Liu, J.; Yu, L.; Niu, X.; Wang, J.; Wang, Y.; Shao, F.; Jin, R.; Wang, Y.; Cao, Y. Fast Evolution of SARS-CoV-2 BA.2.86 to JN.1 under Heavy Immune Pressure. Lancet Infect Dis. 2024, 24, e70–e72. doi: 10.1016/S1473-3099(23)00744-2.

40. Kaku, Y.; Okumura, K.; Padilla-Blanco, M.; Kosugi, Y.; Uriu, K.; Hinay, A. A., Jr; Chen, L.; Plianchaisuk, A.; Kobiyama, K.; Ishii, K. J.; Zahradnik, J.; Ito, J.; Sato, K., K. Virological Characteristics of the SARS-CoV-2 JN.1 Variant. Lancet Infect Dis. 2024, 24, e82. doi: 10.1016/S1473-3099(23)00813-7.

41. Li, P.; Faraone, J. N.; Hsu, C. C.; Chamblee, M.; Zheng, Y.-M.; Carlin, C.; Bednash, J. S.; Horowitz, J. C.; Mallampalli, R. K.; Saif, L. J.; Oltz, E. M.; Jones, D.; Li, J.; Gumina, R. J.; Xu, K.; Liu, S.-L. Characteristics of JN.1-Derived SARS-CoV-2 Subvariants SLip, FLiRT, and KP.2 in Neutralization Escape, Infectivity and Membrane Fusion. bioRxiv 2024, 2024.05.20.595020. doi: 10.1101/2024.05.20.595020.

42. Qu, P.; Faraone, J. N.; Evans, J. P.; Zheng, Y.-M.; Carlin, C.; Anghelina, M.; Stevens, P.; Fernandez, S.; Jones, D.; Panchal, A. R.; Saif, L. J.; Oltz, E. M.; Zhang, B.; Zhou, T.; Xu, K.; Gumina, R. J.; Liu, S.-L. Enhanced Evasion of Neutralizing Ab Response by Omicron XBB.1.5, CH.1.1, and CA.3.1 Variants. Cell Rep. 2023, 42, 112443. doi: 10.1016/j.celrep.2023.112443.

43. Kaku, Y.; Uriu, K.; Kosugi, Y.; Okumura, K.; Yamasoba, D.; Uwamino, Y.; Kuramochi, J.; Sadamasu, K.; Yoshimura, K.; Asakura, H.; Nagashima, M.; Ito, J.; Sato, K. Virological Characteristics of the SARS-CoV-2 KP.2 Variant. Lancet Infect Dis. 2024, 24, e416. doi: 10.1016/S1473-3099(24)00298-6.

44. Kaku, Y.; Yo, M. S.; Tolentino, J. E.; Uriu, K.; Okumura, K.; Ito, J.; Sato, K. Virological Characteristics of the SARS-CoV-2 KP.3, LB.1 and KP.2.3 Variants. bioRxiv 2024, 2024.06.05.597664; doi: 10.1101/2024.06.05.597664

45. Wang, Q.; Mellis, I. A.; Bowen, A.; Kowalski-Dobson, T.; Valdez, R.; Katsamba, P. S.; Shapiro, L.; Gordon, A.; Guo, Y.; Ho, D. D.; Liu, L. Recurrent SARS-CoV-2 Spike Mutations Confer Growth Advantages to Select JN.1 Sublineages. bioRxiv, 2024, 2024.05.29.596362; doi: 10.1101/2024.05.29.596362.

46. Jian, F.; Wang, J.; Yisimayi, A.; Song, W.; Xu, Y.; Chen, X.; Niu, X.; Yang, S.; Yu, Y.; Wang, P.; Sun, H.; Yu, L.; Wang, J.; Wang, Y.; An, R.; Wang, W.; Ma, M.; Xiao, T.; Gu, Q.; Shao, F.; Wang, Y.; Shen, Z.; Jin, R.; Cao, Y. Evolving Antibody Response to SARS-CoV-2 Antigenic Shift from XBB to JN.1. Nature 2024. doi: 10.1038/s41586-024-08315-x.

47. Taylor, A. L.; Starr, T. N. Deep Mutational Scanning of SARS-CoV-2 Omicron BA.2.86 and Epistatic Emergence of the KP.3 Variant. Virus Evol. 2024, 10, veae067. doi: 10.1093/ve/veae067.

48. Feng, L.; Sun, Z.; Zhang, Y.; Jian, F.; Yang, S.; Xia, K.; Yu, L.; Wang, J.; Shao, F.; Wang, X.; Cao, Y. Structural and Molecular Basis of the Epistasis Effect in Enhanced Affinity between SARS-CoV-2 KP.3 and ACE2. Cell Discov. 2024, 10, 123. doi: 10.1038/s41421-024-00752-2.

49. Liu, J.; Yu, Y.; Jian, F.; Yang, S.; Song, W.; Wang, P.; Yu, L.; Shao, F.; Cao, Y. Enhanced Immune Evasion of SARS-CoV-2 KP.3.1.1 and XEC through NTD Glycosylation, bioRxiv, 2024. doi: 10.1101/2024.10.23.619754

50. Kaku, Y.; Uriu, K.; Okumura, K.; Ito, J.; Sato, K. Virological Characteristics of the SARS-CoV-2 KP.3.1.1 Variant. Lancet Infect Dis. 2024, 24, e609. doi: 10.1016/S1473-3099(24)00505-X.

51. Kaku, Y.; Okumura, K.; Kawakubo, S.; Uriu, K.; Chen, L.; Kosugi, Y.; Uwamino, Y.; Begum, M. M.; Leong, S.; Ikeda, T.; Sadamasu, K.; Asakura, H.; Nagashima, M.; Yoshimura, K.; Ito, J.; Sato, K. Virological Characteristics of the SARS-CoV-2 XEC Variant. Lancet Infect Dis. 2024, S1473-3099(24)00731-X. doi: 10.1016/S1473-3099(24)00731-X.

52. Wang, Q.; Guo, Y.; Mellis, I. A.; Wu, M.; Mohri, H.; Gherasim, C.; Valdez, R.; Purpura, L. J.; Yin, M. T.; Gordon, A.; Ho, D. D. Antibody Evasiveness of SARS-CoV-2 Subvariants KP.3.1.1 and XEC. bioRxiv, 2024 2024.11.17.624037; doi: 10.1101/2024.11.17.624037.

53. Cao, Y.; Wang, J.; Jian, F.; Xiao, T.; Song, W.; Yisimayi, A.; Huang, W.; Li, Q.; Wang, P.; An, R.; Wang, J.; Wang, Y.; Niu, X.; Yang, S.; Liang, H.; Sun, H.; Li, T.; Yu, Y.; Cui, Q.; Liu, S.; Yang, X.; Du, S.; Zhang, Z.; Hao, X.; Shao, F.; Jin, R.; Wang, X.; Xiao, J.; Wang, Y.; Xie, X. S. Omicron Escapes the Majority of Existing SARS-CoV-2 Neutralizing Antibodies. Nature 2022, 602, 657–663. doi: 10.1038/s41586-021-04385-3.

54. Cao, Y.; Yisimayi, A.; Jian, F.; Song, W.; Xiao, T.; Wang, L.; Du, S.; Wang, J.; Li, Q.; Chen, X.; Yu, Y.; Wang, P.; Zhang, Z.; Liu, P.; An, R.; Hao, X.; Wang, Y.; Wang, J.; Feng, R.; Sun, H.; Zhao, L.; Zhang, W.; Zhao, D.; Zheng, J.; Yu, L.; Li, C.; Zhang, N.; Wang, R.; Niu, X.; Yang, S.; Song, X.; Chai, Y.; Hu, Y.; Shi, Y.; Zheng, L.; Li, Z.; Gu, Q.; Shao, F.; Huang, W.; Jin, R.; Shen, Z.; Wang, Y.; Wang, X.; Xiao, J.; Xie, X. S. BA.2.12.1, BA.4 and BA.5 Escape Antibodies Elicited by Omicron Infection. Nature 2022, 608, 593–602. doi: 10.1038/s41586-022-04980-y.

55. Barnes, C. O.; Jette, C. A.; Abernathy, M. E.; Dam, K.-M. A.; Esswein, S. R.; Gristick, H. B.; Malyutin, A. G.; Sharaf, N. G.; Huey-Tubman, K. E.; Lee, Y. E.; Robbiani, D. F.; Nussenzweig, M. C.; West, A. P., Jr; Bjorkman, P. J. SARS-CoV-2 Neutralizing Antibody Structures Inform Therapeutic Strategies. Nature 2020, 588, 682–687. doi: 10.1038/s41586-020-2852-1.

56. Cao, Y.; Jian, F.; Wang, J.; Yu, Y.; Song, W.; Yisimayi, A.; Wang, J.; An, R.; Chen, X.; Zhang, N.; Wang, Y.; Wang, P.; Zhao, L.; Sun, H.; Yu, L.; Yang, S.; Niu, X.; Xiao, T.; Gu, Q.; Shao, F.; Hao, X.; Xu, Y.; Jin, R.; Shen, Z.; Wang, Y.; Xie, X. S. Imprinted SARS-CoV-2 Humoral Immunity Induces Convergent Omicron RBD Evolution. Nature 2023, 614, 521–529. doi: 10.1038/s41586-022-05644-7.

57. Cao, Y.; Jian, F.; Zhang, Z.; Yisimayi, A.; Hao, X.; Bao, L.; Yuan, F.; Yu, Y.; Du, S.; Wang, J.; Xiao, T.; Song, W.; Zhang, Y.; Liu, P.; An, R.; Wang, P.; Wang, Y.; Yang, S.; Niu, X.; Zhang, Y.; Gu, Q.; Shao, F.; Hu, Y.; Yin, W.; Zheng, A.; Wang, Y.; Qin, C.; Jin, R.; Xiao, J.; Xie, X. S. Rational Identification of Potent and Broad Sarbecovirus-Neutralizing Antibody Cocktails from SARS Convalescents. Cell Rep. 2022, 41, 111845. doi: 10.1016/j.celrep.2022.111845.

58. Yisimayi, A.; Song, W.; Wang, J.; Jian, F.; Yu, Y.; Chen, X.; Xu, Y.; Yang, S.; Niu, X.; Xiao, T.; Wang, J.; Zhao, L.; Sun, H.; An, R.; Zhang, N.; Wang, Y.; Wang, P.; Yu, L.; Lv, Z.; Gu, Q.; Shao, F.; Jin, R.; Shen, Z.; Xie, X. S.; Wang, Y.; Cao, Y. Repeated Omicron Exposures Override Ancestral SARS-CoV-2 Immune Imprinting. Nature 2024, 625, 148–156. doi: 10.1038/s41586-023-06753-7.

59. Yu, L.; Wang, Y.; Liu, Y.; Xing, X.; Li, C.; Wang, X.; Shi, J.; Ma, W.; Li, J.; Chen, Y.; Qiao, R.; Zhao, X.; Gao, M.; Wen, S.; Xue, Y.; Guan, Y.; Chu, H.; Sun, L.; Wang, P. Potent and Broadly Neutralizing Antibodies against Sarbecoviruses Elicited by Single Ancestral SARS-CoV-2 Infection. bioRxiv 2024, 2024.06.06.597720; doi: 10.1101/2024.06.06.597720.

60. Rosen, L. E.; Tortorici, M. A.; De Marco, A.; Pinto, D.; Foreman, W. B.; Taylor, A. L.; Park, Y.-J.; Bohan, D.; Rietz, T.; Errico, J. M.; Hauser, K.; Dang, H. V.; Chartron, J. W.; Giurdanella, M.; Cusumano, G.; Saliba, C.; Zatta, F.; Sprouse, K. R.; Addetia, A.; Zepeda, S. K.; Brown, J.; Lee, J.; Dellota, E., Jr.; Rajesh, A.; Noack, J.; Tao, Q.; DaCosta, Y.; Tsu, B.; Acosta, R.; Subramanian, S.; de Melo, G. D.; Kergoat, L.; Zhang, I.; Liu, Z.; Guarino, B.; Schmid, M. A.; Schnell, G.; Miller, J. L.; Lempp, F. A.; Czudnochowski, N.; Cameroni, E.; Whelan, S. P. J.; Bourhy, H.; Purcell, L. A.; Benigni, F.; di Iulio, J.; Pizzuto, M. S.; Lanzavecchia, A.; Telenti, A.; Snell, G.; Corti, D.; Veesler, D.; Starr, T. N. A Potent Pan-Sarbecovirus Neutralizing Antibody Resilient to Epitope Diversification. Cell 2024, 187, 7196–7213.e26. 10.1016/j.cell.2024.09.026.

61. Jian, F.; Wec, A. Z.; Feng, L.; Yu, Y.; Wang, L.; Wang, P.; Yu, L.; Wang, J.; Hou, J.; Berrueta, D. M.; Lee, D.; Speidel, T.; Ma, L.; Kim, T.; Yisimayi, A.; Song, W.; Wang, J.; Liu, L.; Yang, S.; Niu, X.; Xiao, T.; An, R.; Wang, Y.; Shao, F.; Wang, Y.; Henry, C.; Pecetta, S.; Wang, X.; Walker, L. M.; Cao, Y. A Generalized Framework to Identify SARS-CoV-2 Broadly Neutralizing Antibodies. bioRxiv 2024. 2024.04.16.589454; doi: 10.1101/2024.04.16.58945.

62. Verkhivker, G.; Alshahrani, M.; Gupta, G. Balancing Functional Tradeoffs between Protein Stability and ACE2 Binding in the SARS-CoV-2 Omicron BA.2, BA.2.75 and XBB Lineages: Dynamics-Based Network Models Reveal Epistatic Effects Modulating Compensatory Dynamic and Energetic Changes. Viruses 2023, 15, 1143. 10.3390/v15051143.

63. Xiao, S.; Alshahrani, M.; Gupta, G.; Tao, P.; Verkhivker, G. Markov State Models and Perturbation-Based Approaches Reveal Distinct Dynamic Signatures and Hidden Allosteric Pockets in the Emerging SARS-Cov-2 Spike Omicron Variant Complexes with the Host Receptor: The Interplay of Dynamics and Convergent Evolution Modulates Allostery and Functional Mechanisms. J. Chem. Inf. Model. 2023, 63, 5272–5296. doi: 10.1021/acs.jcim.3c00778

64. Raisinghani, N.; Alshahrani, M.; Gupta, G.; Xiao, S.; Tao, P.; Verkhivker, G. AlphaFold2 Predictions of Conformational Ensembles and Atomistic Simulations of the SARS-CoV-2 Spike XBB Lineages Reveal Epistatic Couplings between Convergent Mutational Hotspots That Control ACE2 Affinity. J. Phys. Chem. B. 2024, 128, 4696–4715. doi: 10.1021/acs.jpcb.4c01341.

65. Raisinghani, N.; Alshahrani, M.; Gupta, G.; Verkhivker, G. Ensemble-Based Mutational Profiling and Network Analysis of the SARS-CoV-2 Spike Omicron XBB Lineages for Interactions with the ACE2 Receptor and Antibodies: Cooperation of Binding Hotspots in Mediating Epistatic Couplings Underlies Binding Mechanism and Immune Escape. Int. J. Mol. Sci. 2024, 25, 4281. doi: 10.3390/ijms25084281.

66. Raisinghani, N.; Alshahrani, M.; Gupta, G.; Verkhivker, G. AlphaFold2 Modeling and Molecular Dynamics Simulations of the Conformational Ensembles for the SARS-CoV-2 Spike Omicron JN.1, KP.2 and KP.3 Variants: Mutational Profiling of Binding Energetics Reveals Epistatic Drivers of the ACE2 Affinity and Escape Hotspots of Antibody Resistance. Viruses 2024, 16, 1458. doi: 10.3390/v16091458.

67. Verkhivker, G.M.; Di Paola, L. Dynamic Network Modeling of Allosteric Interactions and Communication Pathways in the SARS-CoV-2 Spike Trimer Mutants: Differential Modulation of Conformational Landscapes and Signal Transmission via Cascades of Regulatory Switches. J. Phys. Chem. B 2021, 125, 850–873. 10.1021/acs.jpcb.0c10637.

68. Verkhivker, G.M.; Di Paola, L. Integrated Biophysical Modeling of the SARS-CoV-2 Spike Protein Binding and Allosteric Interactions with Antibodies. J. Phys. Chem. B 2021, 125, 4596–4619. 10.1021/acs.jpcb.1c00395.

69. Verkhivker, G.M.; Agajanian, S.; Oztas, D.Y.; Gupta, G. Comparative Perturbation-Based Modeling of the SARS-CoV-2 Spike Protein Binding with Host Receptor and Neutralizing Antibodies: Structurally Adaptable Allosteric Communication Hotspots Define Spike Sites Targeted by Global Circulating Mutations. Biochemistry 2021, 60, 1459–1484. 10.1021/acs.biochem.1c00139.

70. Verkhivker, G.M.; Agajanian, S.; Oztas, D.Y.; Gupta, G. Dynamic Profiling of Binding and Allosteric Propensities of the SARS-CoV-2 Spike Protein with Different Classes of Antibodies: Mutational and Perturbation-Based Scanning Reveals the Allosteric Duality of Functionally Adaptable Hotspots. J. Chem. Theory Comput. 2021, 17, 4578–4598. 10.1021/acs.jctc.1c00372.

71. Verkhivker, G.M.; Agajanian, S.; Oztas, D.Y.; Gupta, G. Allosteric Control of Structural Mimicry and Mutational Escape in the SARS-CoV-2 Spike Protein Complexes with the ACE2 Decoys and Miniprotein Inhibitors: A Network-Based Approach for Mutational Profiling of Binding and Signaling. J. Chem. Inf. Model. 2021, 61, 5172–5191. 10.1021/acs.jcim.1c00766.

72. Verkhivker, G.; Agajanian, S.; Kassab, R.; Krishnan, K. Integrating Conformational Dynamics and Perturbation-Based Network Modeling for Mutational Profiling of Binding and Allostery in the SARS-CoV-2 Spike Variant Complexes with Antibodies: Balancing Local and Global Determinants of Mutational Escape Mechanisms. Biomolecules 2022, 12, 964. doi: 10.3390/biom12070964.

73. Focosi, D.; Quiroga, R.; McConnell, S.; Johnson, M.C.; Casadevall, A. Convergent Evolution in SARS-CoV-2 Spike Creates a Variant Soup from Which New COVID-19 Waves Emerge. Int. J. Mol. Sci. 2023, 24, 2264. 10.3390/ijms24032264.

74. Gan, H.H.; Twaddle, A.; Marchand, B.; Gunsalus, K.C. Structural Modeling of the SARS-CoV-2 Spike/Human ACE2 Complex Interface can Identify High-Affinity Variants Associated with Increased Transmissibility. J. Mol. Biol. 2021, 433, 167051. doi: 10.1016/j.jmb.2021.167051.

75. Gan, H. H.; Zinno, J.; Piano, F.; Gunsalus, K. C. Omicron Spike Protein Has a Positive Electrostatic Surface That Promotes ACE2 Recognition and Antibody Escape. Front. Virol. 2022, 2. 10.3389/fviro.2022.894531.

76. Barroso da Silva, F. L.; Giron, C. C.; Laaksonen, A. Electrostatic Features for the Receptor Binding Domain of SARS-COV-2 Wildtype and Its Variants. Compass to the Severity of the Future Variants with the Charge-Rule. J. Phys. Chem. B. 2022, 126, 6835–6852. doi: 10.1021/acs.jpcb.2c04225.

77. Li, L.; Shi, K.; Gu, Y.; Xu, Z.; Shu, C.; Li, D.; Sun, J.; Cong, M.; Li, X.; Zhao, X.; Yu, G.; Hu, S.; Tan, H.; Qi, J.; Ma, X.; Liu, K.; Gao, G. F. Spike Structures, Receptor Binding, and Immune Escape of Recently Circulating SARS-CoV-2 Omicron BA.2.86, JN.1, EG.5, EG.5.1, and HV.1 Sub-Variants. Structure 2024, 32, 1055–1067.e6. doi: 10.1016/j.str.2024.06.012.

78. Yang, H.; Guo, H.; Wang, A.; Cao, L.; Fan, Q.; Jiang, J.; Wang, M.; Lin, L.; Ge, X.; Wang, H.; Zhang, R.; Liao, M.; Yan, R.; Ju, B.; Zhang, Z. Structural Basis for the Evolution and Antibody Evasion of SARS-CoV-2 BA.2.86 and JN.1 Subvariants. Nat Commun. 2024, 15, 7715. doi: 10.1038/s41467-024-51973-8.

79. Yajima, H.; Nomai, T.; Okumura, K.; Maenaka, K.; Ito, J.; Hashiguchi, T.; Sato, K.; Matsuno, K.; Nao, N.; Sawa, H.; Mizuma, K.; Li, J.; Kida, I.; Mimura, Y.; Ohari, Y.; Tanaka, S.; Tsuda, M.; Wang, L.; Oda, Y.; Ferdous, Z.; Shishido, K.; Mohri, H.; Iida, M.; Fukuhara, T.; Tamura, T.; Suzuki, R.; Suzuki, S.; Tsujino, S.; Ito, H.; Kaku, Y.; Misawa, N.; Plianchaisuk, A.; Guo, Z.; Hinay, A. A., Jr.; Usui, K.; Saikruang, W.; Lytras, S.; Uriu, K.; Yoshimura, R.; Kawakubo, S.; Nishumura, L.; Kosugi, Y.; Fujita, S.; M. Tolentino, J. E.; Chen, L.; Pan, L.; Li, W.; Yo, M. S.; Horinaka, K.; Suganami, M.; Chiba, M.; Yasuda, K.; Iida, K.; Strange, A. P.; Ohsumi, N.; Tanaka, S.; Ogawa, E.; Fukuda, T.; Osujo, R.; Yoshimura, K.; Sadamas, K.; Nagashima, M.; Asakura, H.; Yoshida, I.; Nakagawa, S.; Takayama, K.; Hashimoto, R.; Deguchi, S.; Watanabe, Y.; Nakata, Y.; Futatsusako, H.; Sakamoto, A.; Yasuhara, N.; Suzuki, T.; Kimura, K.; Sasaki, J.; Nakajima, Y.; Irie, T.; Kawabata, R.; Sasaki-Tabata, K.; Ikeda, T.; Nasser, H.; Shimizu, R.; Begum, M. M.; Jonathan, M.; Mugita, Y.; Leong, S.; Takahashi, O.; Ueno, T.; Motozono, C.; Toyoda, M.; Saito, A.; Kosaka, A.; Kawano, M.; Matsubara, N.; Nishiuchi, T.; Zahradnik, J.; Andrikopoulos, P.; Padilla-Blanco, M.; Konar, A. Molecular and Structural Insights into SARS-CoV-2 Evolution: From BA.2 to XBB Subvariants. mBio. 2024, 15, e0322023. doi: 10.1128/mbio.03220-23.

80. Xue, S.; Han, Y.; Wu, F.; Wang, Q. Mutations in the SARS-CoV-2 Spike Receptor Binding Domain and Their Delicate Balance between ACE2 Affinity and Antibody Evasion. Protein Cell. 2024, 15, 403–418. doi: 10.1093/procel/pwae007.

81. Raisinghani, N.; Alshahrani, M.; Gupta, G.; Xiao, S.; Tao, P.; Verkhivker, G. AlphaFold2 Predictions of Conformational Ensembles and Atomistic Simulations of the SARS-CoV-2 Spike XBB Lineages Reveal Epistatic Couplings between Convergent Mutational Hotspots That Control ACE2 Affinity. J Phys Chem B. 2024, 128, 4696–4715. doi: 10.1021/acs.jpcb.4c01341.

82. Raisinghani, N.; Alshahrani, M.; Gupta, G.; Xiao, S.; Tao, P.; Verkhivker, G. Exploring Conformational Landscapes and Binding Mechanisms of Convergent Evolution for the SARS-CoV-2 Spike Omicron Variant Complexes with the ACE2 Receptor Using AlphaFold2-Based Structural Ensembles and Molecular Dynamics Simulations. Phys Chem Chem Phys. 2024, 26, 17720–17744. doi: 10.1039/d4cp01372g.

83. Rose, P. W.; Prlic, A.; Altunkaya, A.; Bi, C.; Bradley, A. R.; Christie, C. H.; Costanzo, L. D.; Duarte, J. M.; Dutta, S.; Feng, Z.; Green, R. K.; Goodsell, D. S.; Hudson, B.; Kalro, T.; Lowe, R.; Peisach, E.; Randle, C.; Rose, A. S.; Shao, C.; Tao, Y. P.; Valasatava, Y.; Voigt, M.; Westbrook, J. D.; Woo, J.; Yang, H.; Young, J. Y.; Zardecki, C.; Berman, H. M.; Burley, S. K. The RCSB protein data bank: integrative view of protein, gene and 3D structural information. Nucleic Acids Res. 2017, 45, D271–D281. doi: 10.1093/nar/gkw1000.

84. Hekkelman, M.L.; Te Beek, T.A.; Pettifer, S.R.; Thorne, D.; Attwood, T.K.; Vriend, G. WIWS: A protein structure bioinformatics web service collection. Nucleic Acids Res. 2010, 38, W719–W723. 10.1093/nar/gkq453.

85. Fernandez-Fuentes, N.; Zhai, J.; Fiser, A. ArchPRED: A template based loop structure prediction server. Nucleic Acids Res. 2006, 34, W173–W176. 10.1093/nar/gkl113.

86. Krivov, V.P., B.F.; Shapovalov, M.V.; Dunbrack, R.L., Jr. Improved prediction of protein side-chain conformations with SCWRL4. Proteins 2009, 77, 778–795. 10.1002/prot.22488.

87. Søndergaard C. R.; Olsson M. H.; Rostkowski M.; Jensen J. H. Improved treatment of ligands and coupling effects in empirical calculation and rationalization of pKa values. J. Chem. Theory Comput. 2011, 7, 2284–2295. 10.1021/ct200133y.

88. Olsson M. H.; Søndergaard C. R.; Rostkowski M.; Jensen J. H. PROPKA3: consistent treatment of internal and surface residues in empirical pKa predictions. J. Chem. Theory Comput. 2011, 7, 525–537. 10.1021/ct100578z.

89. Bhattacharya, D.; Cheng, J. 3Drefine: Consistent Protein Structure Refinement by Optimizing Hydrogen Bonding Network and Atomic-Level Energy Minimization. Proteins 2013, 81, 119–131. doi: 10.1002/prot.24167.

90. Bhattacharya, D.; Nowotny, J.; Cao, R.; Cheng, J. 3Drefine: An Interactive Web Server for Efficient Protein Structure Refinement. Nucleic Acids Res. 2016, 44, W406–W409. doi: 10.1093/nar/gkw336.

91. Phillips, J.C.; Hardy, D.J.; Maia, J.D.C.; Stone, J.E.; Ribeiro, J.V.; Bernardi, R.C.; Buch, R.; Fiorin, G.; Hénin, J.; Jiang, W.;, et al. Scalable Molecular Dynamics on CPU and GPU Architectures with NAMD. J. Chem. Phys. 2020, 153, 044130. 10.1063/5.0014475.

92. Huang, J.; Rauscher, S.; Nawrocki, G.; Ran, T.; Feig, M.; de Groot, B.L.; Grubmüller, H.; MacKerell, A.D., Jr. CHARMM36m: An improved force field for folded and intrinsically disordered proteins. Nat. Methods 2017, 14, 71–73. 10.1038/nmeth.4067.

93. Fernandes, H.S.; Sousa, S.F.; Cerqueira, N.M.F.S.A. VMD Store-A VMD Plugin to Browse, Discover, and Install VMD Extensions. J. Chem. Inf. Model. 2019, 59, 4519–4523. doi: 10.1021/acs.jcim.9b00739.

94. Jo, S.; Kim, T.; Iyer, V. G.; Im, W. CHARMM GUI: A Web based Graphical User Interface for CHARMM. J Comput Chem. 2008, 29, 1859–1865. doi: 10.1002/jcc.20945.

95. Lee, J.; Cheng, X.; Swails, J. M.; Yeom, M. S.; Eastman, P. K.; Lemkul, J. A.; Wei, S.; Buckner, J.; Jeong, J. C.; Qi, Y.; Jo, S.; Pande, V. S.; Case, D. A.; Brooks, C. L., III; MacKerell, A. D., Jr.; Klauda, J. B.; Im, W. CHARMM-GUI Input Generator for NAMD, GROMACS, AMBER, OpenMM, and CHARMM/OpenMM Simulations Using the CHARMM36 Additive Force Field. J Chem Theory Comput. 2016, 12, 405–413. doi: 10.1021/acs.jctc.5b00935.

96. Jorgensen, W.L.; Chandrasekhar, J.; Madura, J.D.; Impey, R.W.; Klein, M.L. Comparison of Simple Potential Functions for Simulating Liquid Water. J. Chem. Phys. 1983, 79, 926–935. 10.1063/1.445869.

97. Ross, G.A.; Rustenburg, A.S.; Grinaway, P.B.; Fass, J.; Chodera, J.D. Biomolecular Simulations under Realistic Macroscopic Salt Conditions. J. Phys. Chem. B 2018, 122, 5466–5486. 10.1021/acs.jpcb.7b11734.

98. Di Pierro, M.; Elber, R.; Leimkuhler, B. A Stochastic Algorithm for the Isobaric-Isothermal Ensemble with Ewald Summations for All Long Range Forces. J. Chem. Theory Comput. 2015, 11, 5624–5637. 10.1021/acs.jctc.5b00648.

99. Martyna, G.J.; Tobias, D.J.; Klein, M.L. Constant pressure molecular dynamics algorithms. J. Chem. Phys. 1994, 101, 4177–4189. 10.1063/1.467468.

100. Feller, S.E.; Zhang, Y.; Pastor, R.W.; Brooks, B.R. Constant pressure molecular dynamics simulation: The Langevin piston method. J. Chem. Phys. 1995, 103, 4613–4621. 10.1063/1.470648.

101. Davidchack, R.L.; Handel, R.; Tretyakov, M.V. Langevin thermostat for rigid body dynamics. J. Chem. Phys. 2009, 130, 234101. 10.1063/1.3149788.

102. Dehouck, Y.; Kwasigroch, J. M.; Rooman, M.; Gilis, D. BeAtMuSiC: Prediction of changes in protein-protein binding affinity on mutations. Nucleic Acids Res. 2013, 41, W333–W339. doi: 10.1093/nar/gkt450.

103. Dehouck, Y.; Gilis, D.; Rooman, M. A new generation of statistical potentials for proteins. Biophys. J. 2006, 90, 4010–4017. doi: 10.1529/biophysj.105.079434.

104. Dehouck, Y.; Grosfils, A.; Folch, B.; Gilis, D.; Bogaerts, P.; Rooman, M. Fast and accurate predictions of protein stability changes upon mutations using statistical potentials and neural networks:PoPMuSiC-2.0. Bioinformatics 2009, 25, 2537–2543. Doi:10.1093/bioinformatics/btp445.

105. Srinivasan, J.; Cheatham, T. E.; Cieplak, P.; Kollman, P. A.; Case, D. A. Continuum Solvent Studies of the Stability of DNA, RNA, and Phosphoramidate−DNA Helices. J. Amer. Chem. Soc. 1998, 120, 9401–9409. 10.1021/ja981844.

106. Kollman, P. A.; Massova, I.; Reyes, C.; Kuhn, B.; Huo, S.; Chong, L.; Lee, M.; Lee, T.; Duan, Y.; Wang, W.; Donini, O.; Cieplak, P.; Srinivasan, J.; Case, D. A.; Cheatham, T. E. Calculating Structures and Free Energies of Complex Molecules: Combining Molecular Mechanics and Continuum Models. Acc. Chem. Res. 2000, 33, 889–897. 10.1021/ar000033j.

107. Hou, T.; Wang, J.; Li, Y.; Wang, W. Assessing the Performance of the MM/PBSA and MM/GBSA Methods. 1. The Accuracy of Binding Free Energy Calculations Based on Molecular Dynamics Simulations. J. Chem. Inf. Model. 2011, 51, 69–82. 10.1021/ci100275a.

108. Weng, G.; Wang, E.; Wang, Z.; Liu, H.; Zhu, F.; Li, D.; Hou, T. HawkDock: A Web Server to Predict and Analyze the Protein–Protein Complex Based on Computational Docking and MM/GBSA. Nucleic Acids Res. 2019, 47, W322–W330. 10.1093/nar/gkz397.

109. Mongan, J.; Simmerling, C.; McCammon, J. A.; Case, D. A.; Onufriev, A. Generalized Born Model with a Simple, Robust Molecular Volume Correction. J Chem Theory Comput. 2007, 3, 156–169. doi: 10.1021/ct600085e.

110. Williams, A. H.; Zhan, C.-G. Generalized Methodology for the Quick Prediction of Variant SARS-CoV-2 Spike Protein Binding Affinities with Human Angiotensin-Converting Enzyme II. J. Phys. Chem. B. 2022, 126, 2353–2360. doi: 10.1021/acs.jpcb.1c10718.

111. Sun, H.; Duan, L.; Chen, F.; Liu, H.; Wang, Z.; Pan, P.; Zhu, F.; Zhang, J. Z. H.; Hou, T. Assessing the Performance of MM/PBSA and MM/GBSA Methods. 7. Entropy Effects on the Performance of End-Point Binding Free Energy Calculation Approaches. Phys. Chem. Chem. Phys. 2018, 20, 14450–14460. doi: 10.1039/c7cp07623a.

112. Miller, B. R., III; McGee, T. D., Jr.; Swails, J. M.; Homeyer, N.; Gohlke, H.; Roitberg, A. E. MMPBSA.Py: An Efficient Program for End-State Free Energy Calculations. J Chem Theory Comput. 2012, 8, 3314–3321. doi: 10.1021/ct300418h.

113. Valdés-Tresanco, M. S.; Valdés-Tresanco, M. E.; Valiente, P. A.; Moreno, E. gmx_MMPBSA: A New Tool to Perform End-State Free Energy Calculations with GROMACS. J Chem Theory Comput. 2021, 17, 6281–6291. doi: 10.1021/acs.jctc.1c00645.

114. Hadfield, J.; Megill, C.; Bell, S. M.; Huddleston, J.; Potter, B.; Callender, C.; Sagulenko, P.; Bedford, T.; Neher, R. A. Nextstrain: Real-Time Tracking of Pathogen Evolution. Bioinformatics 2018, 34, 4121–4123. doi: 10.1093/bioinformatics/bty407.

115. Bhattarai, N.; Baral, P.; Gerstman, B. S.; Chapagain, P. P. Structural and Dynamical Differences in the Spike Protein RBD in the SARS-CoV-2 Variants B.1.1.7 and B.1.351. J Phys Chem B. 2021, 125, 7101–7107. doi: 10.1021/acs.jpcb.1c01626.

116. Valério, M.; Borges-Araújo, L.; Melo, M. N.; Lousa, D.; Soares, C. M. SARS-CoV-2 Variants Impact RBD Conformational Dynamics and ACE2 Accessibility. Front Med Technol. 2022, 4, 1009451. doi: 10.3389/fmedt.2022.1009451.

117. Li, P.; Faraone, J. N.; Hsu, C. C.; Chamblee, M.; Liu, Y.; Zheng, Y.-M.; Xu, Y.; Carlin, C.; Horowitz, J. C.; Mallampalli, R. K.; Saif, L. J.; Oltz, E. M.; Jones, D.; Li, J.; Gumina, R. J.; Bednash, J. S.; Xu, K.; Liu, S.-L. Immune Evasion, Cell-Cell Fusion, and Spike Stability of the SARS-CoV-2 XEC Variant: Role of Glycosylation Mutations at the N-Terminal Domain. bioRxiv 2024. doi: 10.1101/2024.11.12.623078.

118. Wang, Q.; Guo, Y.; Mellis, I. A.; Wu, M.; Mohri, H.; Gherasim, C.; Valdez, R.; Purpura, L. J.; Yin, M. T.; Gordon, A.; Ho, D. D. Antibody Evasiveness of SARS-CoV-2 Subvariants KP.3.1.1 and XEC, 2024. 10.1101/2024.11.17.624037. Ab evasiveness of SARS-CoV-2 subvariants KP.3.1.1 and XEC. bioRxiv 2024. doi: 10.1101/2024.11.17.624037

119. Greaney, A. J.; Starr, T. N.; Bloom, J. D. An Antibody-Escape Estimator for Mutations to the SARS-CoV-2 Receptor-Binding Domain. Virus Evol. 2022, 8, veac021. doi: 10.1093/ve/veac021.

120. Dadonaite, B.; Crawford, K. H. D.; Radford, C. E.; Farrell, A. G.; Yu, T. C.; Hannon, W. W.; Zhou, P.; Andrabi, R.; Burton, D. R.; Liu, L.; Ho, D. D.; Chu, H. Y.; Neher, R. A.; Bloom, J. D. A Pseudovirus System Enables Deep Mutational Scanning of the Full SARS-CoV-2 Spike. Cell 2023, 186, 1263–1278.e20. doi: 10.1016/j.cell.2023.02.001.

121. Dadonaite, B.; Brown, J.; McMahon, T. E.; Farrell, A. G.; Figgins, M. D.; Asarnow, D.; Stewart, C.; Lee, J.; Logue, J.; Bedford, T.; Murrell, B.; Chu, H. Y.; Veesler, D.; Bloom, J. D. Spike Deep Mutational Scanning Helps Predict Success of SARS-CoV-2 Clades. Nature. 2024, 631, 617–626. doi: 10.1038/s41586-024-07636-1.

122. Weisblum, Y.; Schmidt, F.; Zhang, F.; DaSilva, J.; Poston, D.; Lorenzi, J. C.; Muecksch, F.; Rutkowska, M.; Hoffmann, H.-H.; Michailidis, E.; Gaebler, C.; Agudelo, M.; Cho, A.; Wang, Z.; Gazumyan, A.; Cipolla, M.; Luchsinger, L.; Hillyer, C. D.; Caskey, M.; Robbiani, D. F.; Rice, C. M.; Nussenzweig, M. C.; Hatziioannou, T.; Bieniasz, P. D. Escape from Neutralizing Antibodies by SARS-CoV-2 Spike Protein Variants. Elife 2020, 9, e61312. doi: 10.7554/eLife.61312.

123. Cameroni, E.; Bowen, J. E.; Rosen, L. E.; Saliba, C.; Zepeda, S. K.; Culap, K.; Pinto, D.; VanBlargan, L. A.; De Marco, A.; di Iulio, J.; Zatta, F.; Kaiser, H.; Noack, J.; Farhat, N.; Czudnochowski, N.; Havenar-Daughton, C.; Sprouse, K. R.; Dillen, J. R.; Powell, A. E.; Chen, A.; Maher, C.; Yin, L.; Sun, D.; Soriaga, L.; Bassi, J.; Silacci-Fregni, C.; Gustafsson, C.; Franko, N. M.; Logue, J.; Iqbal, N. T.; Mazzitelli, I.; Geffner, J.; Grifantini, R.; Chu, H.; Gori, A.; Riva, A.; Giannini, O.; Ceschi, A.; Ferrari, P.; Cippà, P. E.; Franzetti-Pellanda, A.; Garzoni, C.; Halfmann, P. J.; Kawaoka, Y.; Hebner, C.; Purcell, L. A.; Piccoli, L.; Pizzuto, M. S.; Walls, A. C.; Diamond, M. S.; Telenti, A.; Virgin, H. W.; Lanzavecchia, A.; Snell, G.; Veesler, D.; Corti, D. Broadly Neutralizing Antibodies Overcome SARS-CoV-2 Omicron Antigenic Shift. Nature 2022, 602, 664–670. doi: 10.1038/s41586-021-04386-2.

124. Addetia, A.; Piccoli, L.; Case, J. B.; Park, Y.-J.; Beltramello, M.; Guarino, B.; Dang, H.; de Melo, G. D.; Pinto, D.; Sprouse, K.; Scheaffer, S. M.; Bassi, J.; Silacci-Fregni, C.; Muoio, F.; Dini, M.; Vincenzetti, L.; Acosta, R.; Johnson, D.; Subramanian, S.; Saliba, C.; Giurdanella, M.; Lombardo, G.; Leoni, G.; Culap, K.; McAlister, C.; Rajesh, A.; Dellota, E., Jr; Zhou, J.; Farhat, N.; Bohan, D.; Noack, J.; Chen, A.; Lempp, F. A.; Quispe, J.; Kergoat, L.; Larrous, F.; Cameroni, E.; Whitener, B.; Giannini, O.; Cippà, P.; Ceschi, A.; Ferrari, P.; Franzetti-Pellanda, A.; Biggiogero, M.; Garzoni, C.; Zappi, S.; Bernasconi, L.; Kim, M. J.; Rosen, L. E.; Schnell, G.; Czudnochowski, N.; Benigni, F.; Franko, N.; Logue, J. K.; Yoshiyama, C.; Stewart, C.; Chu, H.; Bourhy, H.; Schmid, M. A.; Purcell, L. A.; Snell, G.; Lanzavecchia, A.; Diamond, M. S.; Corti, D.; Veesler, D. Neutralization, Effector Function and Immune Imprinting of Omicron Variants. Nature 2023, 621, 592–601. doi: 10.1038/s41586-023-06487-6.

125. Parsons, R. J.; Acharya, P. Evolution of the SARS-CoV-2 Omicron Spike. Cell Rep. 2023, 42, 113444. doi: 10.1016/j.celrep.2023.113444.

126. Božič, A.; Podgornik, R. Changes in Total Charge on Spike Protein of SARS-CoV-2 in Emerging Lineages.. Bioinform Adv. 2024, 4, vbae053. doi: 10.1093/bioadv/vbae053.

