## Supplemental Figures S1-S3, Tables S1-S6 for "Quantitative Characterization and Prediction of the Binding Determinants and Immune Escape Hotspots for Groups of Broadly Neutralizing Antibodies Against Omicron Variants: Atomistic Modeling of the SARS-CoV-2 Spike Complexes with Antibodies"

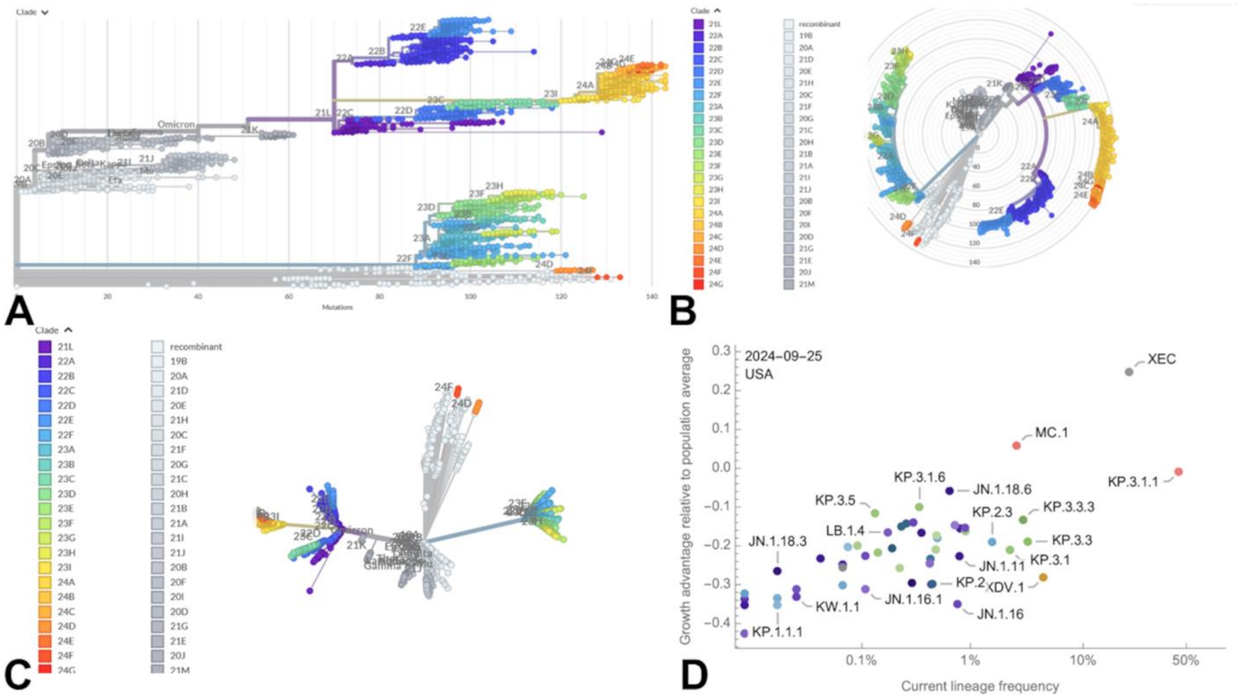


**Figure S1.** An overview of the phylogenetic analysis and SARS-CoV-2 clade classification highlights the evolution of SARSB-CoV-2 lineages using rectangular phylogenetic tree representation (A), radial phylogenetic tree representation (B) and unrooted phylogenetic tree representation (C ). The recent data on growth advantage relative to population average in US (https://github.com/nextstrain/ncov/pull/1152) (D). These plots illustrate evolutionary trajectories of Omicron lineages can proceed through complex recombination, antigenic drift and convergent evolution. The graphs are generated using Nextstrain, an open-source project for real time tracking of evolving pathogen populations (https://nextstrain.org/).

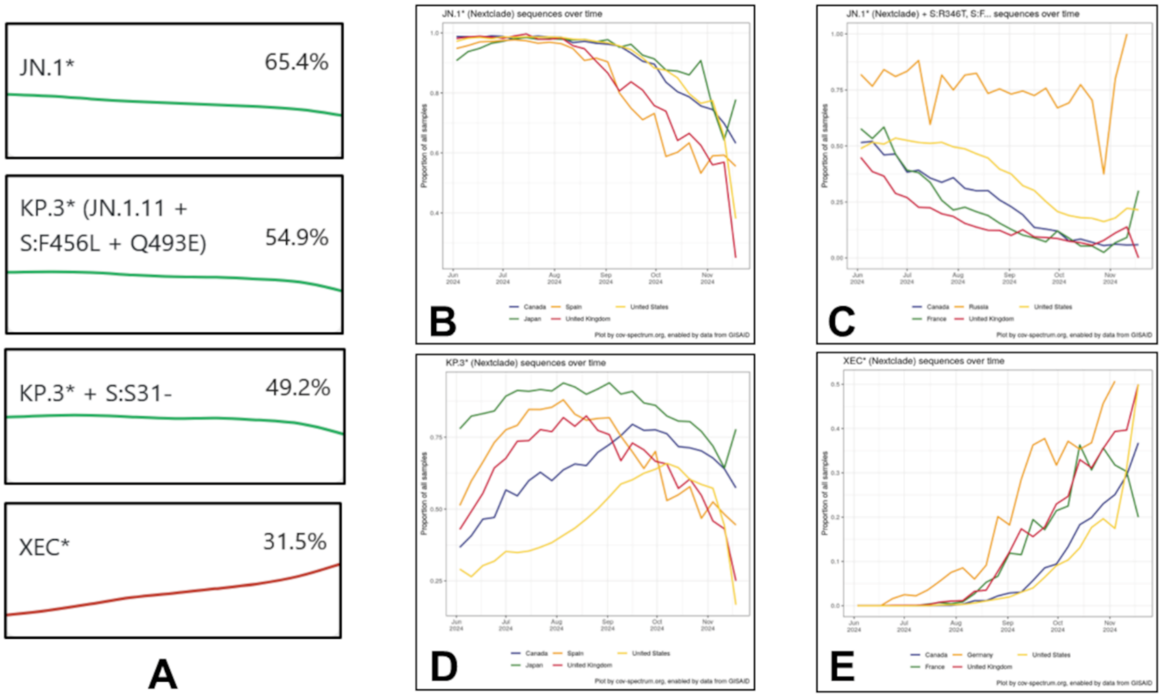


**Figure S2.** Cov-Spectrum data ( https://cov-spectrum.org/explore/World/AllSamples/) for the dominant variants of the samples collected and analyzed within the specified date range (June 3, 2024, to November 26, 2024). This percentage represents the prevalence of the JN.1 variant among all the samples in the world tested during that period. The high proportion of the world samples collected and analyzed during this latest period is dominated by other JN.1 descendants, particularly 54.8% for KP.3, 49.2 % for KP.3.1.1 and 31.5 % for XEC variant. Notice that the sum of percentages for the presented variants on Cov-Spectrum can exceed 100% because some samples may contain multiple variants

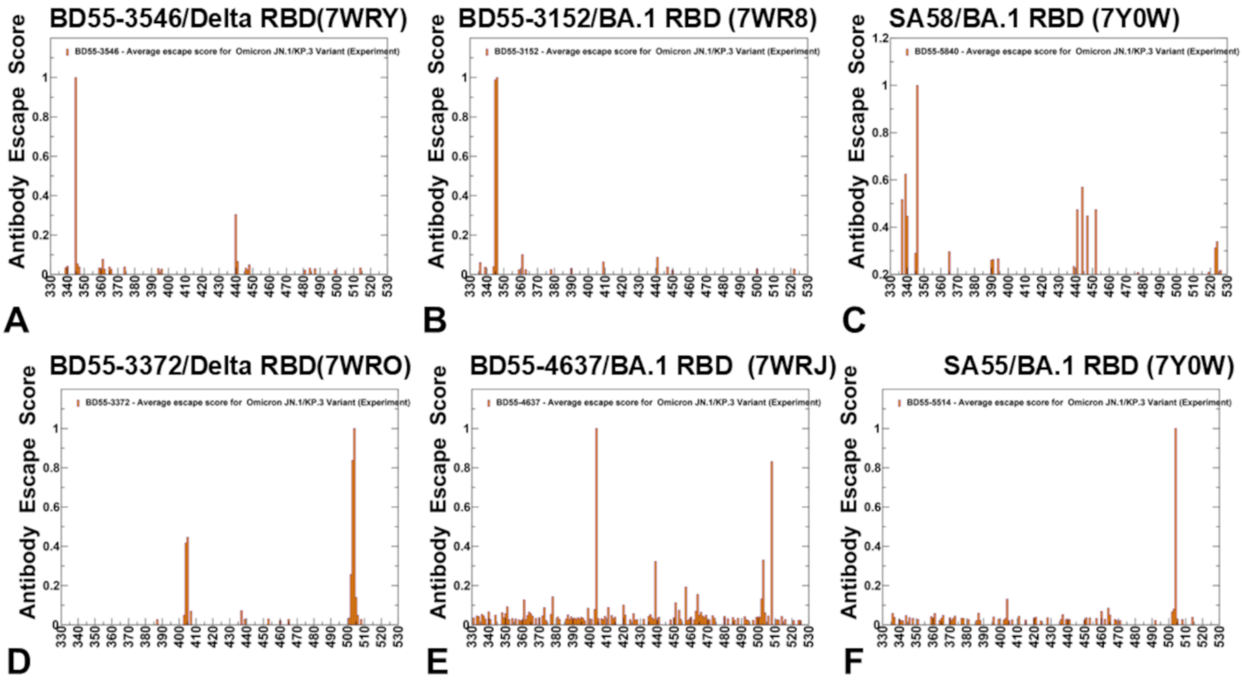


**Figure S3.** The residue-based averaged Ab escape scores derived from the latest experimental data [46] for E1 group Abs BD55-3546 (A), BD55-3152 (B), SA58 (C), and F3 group Ab BD55-3372 (D), BD55-4637 (E) and SA55 (F).

**Table S1.** The list of the binding epitope residues and BD55-3546 Ab residues forming interfacial contacts

| RBD Residue | RBD Residue Number | RBD Chain | Antibody residue | Antibody residue number | Antibody chain |
|---|---|---|---|---|---|
| GLY | 339 | R | TYR | 93 | L |
| GLY | 339 | R | TRP | 94 | L |
| GLU | 340 | R | ASN | 28 | L |
| GLU | 340 | R | TYR | 93 | L |
| GLU | 340 | R | HIS | 27 | L |
| VAL | 341 | R | TYR | 93 | L |
| ASN | 343 | R | TYR | 93 | L |
| ASN | 343 | R | PRO | 95 | L |
| ASN | 343 | R | ASN | 92 | L |
| ASN | 343 | R | TRP | 94 | L |
| ALA | 344 | R | TRP | 94 | L |
| ALA | 344 | R | TYR | 93 | L |
| ALA | 344 | R | ASN | 92 | L |
| THR | 345 | R | LEU | 102 | H |
| THR | 345 | R | TYR | 93 | L |
| THR | 345 | R | ARG | 105 | H |
| THR | 345 | R | TRP | 94 | L |
| THR | 345 | R | TYR | 103 | H |

| | | | | | |
|---|---|---|---|---|---|
| THR | 345 | R | LEU | 96 | L |
| THR | 345 | R | TYR | 91 | L |
| THR | 345 | R | ASN | 92 | L |
| ARG | 346 | R | TYR | 103 | H |
| ARG | 346 | R | GLU | 104 | H |
| ARG | 346 | R | ASN | 92 | L |
| PHE | 347 | R | TYR | 103 | H |
| LYS | 356 | R | TYR | 93 | L |
| ASN | 439 | R | ASN | 52 | H |
| ASN | 439 | R | THR | 55 | H |
| ASN | 440 | R | PRO | 58 | H |
| ASN | 440 | R | ILE | 57 | H |
| ASN | 440 | R | THR | 59 | H |
| ASN | 440 | R | ARG | 105 | H |
| ASN | 440 | R | LEU | 102 | H |
| ASN | 440 | R | TRP | 50 | H |
| ASN | 440 | R | THR | 55 | H |
| ASN | 440 | R | ASN | 52 | H |
| LEU | 441 | R | ARG | 105 | H |
| LEU | 441 | R | TRP | 94 | L |
| LEU | 441 | R | ASN | 52 | H |
| LEU | 441 | R | TYR | 103 | H |
| LEU | 441 | R | LEU | 102 | H |

| | | | | | |
|---|---|---|---|---|---|
| LEU | 441 | R | TRP | 50 | H |
| ASP | 442 | R | LEU | 102 | H |
| ASP | 442 | R | TYR | 103 | H |
| SER | 443 | R | LEU | 102 | H |
| SER | 443 | R | ASN | 52 | H |
| SER | 443 | R | THR | 55 | H |
| SER | 443 | R | TYR | 103 | H |
| SER | 443 | R | ASN | 54 | H |
| LYS | 444 | R | ASN | 31 | H |
| LYS | 444 | R | ASN | 54 | H |
| LYS | 444 | R | LEU | 102 | H |
| VAL | 445 | R | ASN | 31 | H |
| VAL | 445 | R | ASN | 54 | H |
| VAL | 445 | R | ILE | 30 | H |
| ASN | 448 | R | LEU | 102 | H |
| ASN | 448 | R | TYR | 103 | H |
| ASN | 450 | R | TYR | 103 | H |
| TYR | 451 | R | TYR | 103 | H |
| PRO | 499 | R | THR | 55 | H |
| PRO | 499 | R | ASN | 54 | H |
| ARG | 509 | R | TRP | 94 | L |
| ARG | 509 | R | TYR | 103 | H |

**Table S2.** The list of the binding epitope residues and BD55-3152 Ab residues forming interfacial contacts

| RBD Residue | RBD Residue Number | RBD Chain | Antibody residue | Antibody residue number | Antibody chain |
|---|---|---|---|---|---|
| ASP | 339 | R | SER | 93 | B |
| ASP | 339 | R | ALA | 92 | B |
| ASP | 339 | R | THR | 94 | B |
| GLU | 340 | R | ALA | 92 | B |
| GLU | 340 | R | ASP | 91 | B |
| GLU | 340 | R | SER | 93 | B |
| GLU | 340 | R | SER | 29 | B |
| VAL | 341 | R | ALA | 92 | B |
| PHE | 342 | R | GLY | 110 | A |
| PHE | 342 | R | TRP | 109 | A |
| ASN | 343 | R | SER | 93 | B |
| ASN | 343 | R | SER | 111 | A |
| ASN | 343 | R | GLY | 110 | A |
| ASN | 343 | R | THR | 94 | B |
| ASN | 343 | R | ALA | 92 | B |
| ASN | 343 | R | TRP | 109 | A |
| ASN | 343 | R | GLN | 30 | B |
| ASN | 343 | R | TYR | 107 | A |

| | | | | | |
|---|---|---|---|---|---|
| ALA | 344 | R | GLY | 110 | A |
| ALA | 344 | R | SER | 29 | B |
| ALA | 344 | R | ALA | 92 | B |
| ALA | 344 | R | SER | 93 | B |
| ALA | 344 | R | TRP | 109 | A |
| ALA | 344 | R | GLN | 30 | B |
| THR | 345 | R | SER | 93 | B |
| THR | 345 | R | GLY | 110 | A |
| THR | 345 | R | TYR | 33 | B |
| THR | 345 | R | TYR | 31 | B |
| THR | 345 | R | SER | 111 | A |
| THR | 345 | R | PRO | 112 | A |
| THR | 345 | R | LEU | 113 | A |
| THR | 345 | R | SER | 29 | B |
| THR | 345 | R | THR | 94 | B |
| THR | 345 | R | GLN | 30 | B |
| ARG | 346 | R | PRO | 28 | B |
| ARG | 346 | R | SER | 29 | B |
| ARG | 346 | R | GLY | 67 | B |
| ARG | 346 | R | TYR | 31 | B |
| ARG | 346 | R | LEU | 27 | B |
| ARG | 346 | R | THR | 65 | B |
| ARG | 346 | R | VAL | 32 | B |

| | | | | | |
|---|---|---|---|---|---|
| ARG | 346 | R | GLN | 30 | B |
| ARG | 346 | R | ASP | 50 | B |
| PHE | 347 | R | TRP | 109 | A |
| ASN | 354 | R | PRO | 28 | B |
| ASN | 354 | R | SER | 29 | B |
| VAL | 367 | R | TRP | 109 | A |
| LEU | 368 | R | TRP | 109 | A |
| LEU | 371 | R | TRP | 109 | A |
| ALA | 372 | R | TYR | 107 | A |
| PHE | 375 | R | TRP | 109 | A |
| PHE | 375 | R | TYR | 107 | A |
| TRP | 436 | R | VAL | 108 | A |
| TRP | 436 | R | TRP | 109 | A |
| ASN | 437 | R | VAL | 108 | A |
| SER | 438 | R | VAL | 108 | A |
| LYS | 440 | R | PRO | 102 | A |
| LYS | 440 | R | LEU | 103 | A |
| LYS | 440 | R | PHE | 101 | A |
| LYS | 440 | R | SER | 104 | A |
| LYS | 440 | R | VAL | 108 | A |
| LYS | 440 | R | ASP | 105 | A |
| LEU | 441 | R | TRP | 109 | A |
| LEU | 441 | R | VAL | 108 | A |

| | | | | | |
|---|---|---|---|---|---|
| LEU | 441 | R | GLY | 110 | A |
| LEU | 441 | R | PHE | 101 | A |
| LEU | 441 | R | TYR | 31 | B |
| LEU | 441 | R | SER | 111 | A |
| LEU | 441 | R | PRO | 112 | A |
| ASP | 442 | R | TYR | 31 | B |
| LYS | 444 | R | GLU | 52 | B |
| ASN | 448 | R | GLU | 52 | B |
| ASN | 448 | R | TYR | 31 | B |
| TYR | 451 | R | TYR | 31 | B |
| ARG | 509 | R | TRP | 109 | A |
| ARG | 509 | R | GLY | 110 | A |
| ARG | 509 | R | TYR | 31 | B |
| ARG | 509 | R | SER | 111 | A |

**Table S3.** The list of the binding epitope residues and SA58 Ab residues forming interfacial contacts

| RBD Residue | RBD Residue Number | RBD Chain | Antibody residue | Antibody residue number | Antibody chain |
|---|---|---|---|---|---|
| PRO | 337 | R | LEU | 29 | L |
| PRO | 337 | R | SER | 28 | L |
| ASP | 339 | R | ASN | 95 | L |
| ASP | 339 | R | GLU | 1 | L |
| GLU | 340 | R | ALA | 27 | L |
| GLU | 340 | R | SER | 28 | L |
| GLU | 340 | R | ASN | 95 | L |
| GLU | 340 | R | GLU | 1 | L |
| GLU | 340 | R | ARG | 26 | L |
| GLU | 340 | R | GLY | 30 | L |
| GLU | 340 | R | LEU | 29 | L |
| GLU | 340 | R | VAL | 2 | L |
| VAL | 341 | R | ASN | 95 | L |
| VAL | 341 | R | LEU | 29 | L |
| ASN | 343 | R | SER | 94 | L |
| ASN | 343 | R | PRO | 97 | L |
| ASN | 343 | R | ASN | 95 | L |

| | | | | | |
|---|---|---|---|---|---|
| ASN | 343 | R | TRP | 96 | L |
| ALA | 344 | R | SER | 94 | L |
| ALA | 344 | R | ASN | 95 | L |
| ALA | 344 | R | TRP | 96 | L |
| THR | 345 | R | TRP | 96 | L |
| THR | 345 | R | TYR | 105 | H |
| THR | 345 | R | ASP | 34 | L |
| THR | 345 | R | TYR | 93 | L |
| THR | 345 | R | LEU | 98 | L |
| THR | 345 | R | SER | 94 | L |
| THR | 345 | R | ASN | 95 | L |
| ARG | 346 | R | PHE | 106 | H |
| ARG | 346 | R | SER | 94 | L |
| ARG | 346 | R | SER | 103 | H |
| ARG | 346 | R | ASP | 104 | H |
| ARG | 346 | R | TYR | 105 | H |
| ARG | 346 | R | ASP | 34 | L |
| ARG | 346 | R | TYR | 93 | L |
| LYS | 356 | R | LEU | 29 | L |
| ARG | 357 | R | LEU | 29 | L |
| ILE | 358 | R | LEU | 29 | L |
| LYS | 440 | R | TRP | 50 | H |
| LYS | 440 | R | THR | 57 | H |

| | | | | | |
|---|---|---|---|---|---|
| LYS | 440 | R | ASN | 32 | H |
| LYS | 440 | R | PRO | 58 | H |
| LYS | 440 | R | ASN | 52 | H |
| LYS | 440 | R | TYR | 102 | H |
| LYS | 440 | R | THR | 59 | H |
| LEU | 441 | R | ASN | 52 | H |
| LEU | 441 | R | TRP | 96 | L |
| LEU | 441 | R | TYR | 102 | H |
| LEU | 441 | R | TRP | 50 | H |
| LEU | 441 | R | SER | 103 | H |
| LEU | 441 | R | TYR | 105 | H |
| ASP | 442 | R | TYR | 102 | H |
| ASP | 442 | R | SER | 103 | H |
| ASP | 442 | R | TYR | 105 | H |
| SER | 443 | R | ASP | 54 | H |
| SER | 443 | R | TYR | 102 | H |
| SER | 443 | R | ASN | 32 | H |
| LYS | 444 | R | THR | 30 | H |
| LYS | 444 | R | ASN | 32 | H |
| LYS | 444 | R | SER | 31 | H |
| LYS | 444 | R | ASP | 54 | H |
| LYS | 444 | R | TYR | 102 | H |
| VAL | 445 | R | ASP | 54 | H |

| | | | | | |
|---|---|---|---|---|---|
| ASN | 448 | R | TYR | 102 | H |
| ASN | 448 | R | SER | 103 | H |
| ASN | 450 | R | TYR | 102 | H |
| ASN | 450 | R | SER | 103 | H |
| TYR | 451 | R | SER | 103 | H |
| ARG | 509 | R | TYR | 105 | H |
| ARG | 509 | R | TRP | 96 | L |

**Table S4.** The list of the binding epitope residues and BD55-3372 Ab residues forming interfacial contacts

| RBD Residue | RBD Residue Number | RBD Chain | Antibody residue | Antibody residue number | Antibody chain |
|---|---|---|---|---|---|
| ALA | 372 | R | LEU | 116 | L |
| ARG | 403 | R | ASN | 50 | H |
| GLY | 404 | R | SER | 75 | H |
| GLY | 404 | R | PHE | 76 | H |
| ASP | 405 | R | PHE | 76 | H |
| ASP | 405 | R | THR | 71 | H |
| ASP | 405 | R | ASN | 50 | H |
| ASP | 405 | R | SER | 73 | H |
| ASP | 405 | R | SER | 75 | H |
| ASP | 405 | R | GLY | 72 | H |
| ASP | 405 | R | SER | 74 | H |
| GLU | 406 | R | SER | 73 | H |
| VAL | 407 | R | SER | 75 | H |
| VAL | 407 | R | PHE | 76 | H |
| ARG | 408 | R | SER | 75 | H |
| ARG | 408 | R | SER | 74 | H |
| ARG | 408 | R | SER | 73 | H |

| | | | | | |
|---|---|---|---|---|---|
| GLN | 409 | R | SER | 73 | H |
| ASN | 437 | R | SER | 114 | L |
| ASN | 439 | R | TYR | 52 | L |
| GLN | 498 | R | ASP | 120 | H |
| GLN | 498 | R | TYR | 122 | H |
| GLN | 498 | R | ASP | 121 | H |
| PRO | 499 | R | TYR | 52 | L |
| PRO | 499 | R | ALA | 50 | L |
| PRO | 499 | R | GLY | 51 | L |
| THR | 500 | R | GLY | 51 | L |
| THR | 500 | R | ASP | 123 | H |
| THR | 500 | R | ASP | 121 | H |
| THR | 500 | R | GLU | 53 | L |
| THR | 500 | R | THR | 124 | H |
| THR | 500 | R | TYR | 122 | H |
| THR | 500 | R | TYR | 112 | L |
| THR | 500 | R | TYR | 52 | L |
| ASN | 501 | R | ARG | 119 | H |
| ASN | 501 | R | SER | 114 | L |
| ASN | 501 | R | TYR | 122 | H |
| ASN | 501 | R | TYR | 112 | L |
| ASN | 501 | R | ASP | 120 | H |
| ASN | 501 | R | TYR | 52 | L |

| | | | | | |
|---|---|---|---|---|---|
| ASN | 501 | R | ASP | 123 | H |
| ASN | 501 | R | ASP | 121 | H |
| GLY | 502 | R | TYR | 122 | H |
| GLY | 502 | R | TYR | 112 | L |
| GLY | 502 | R | TYR | 52 | L |
| GLY | 502 | R | ASP | 123 | H |
| GLY | 502 | R | GLU | 118 | H |
| VAL | 503 | R | THR | 71 | H |
| VAL | 503 | R | TYR | 112 | L |
| VAL | 503 | R | SER | 119 | L |
| VAL | 503 | R | TYR | 52 | L |
| VAL | 503 | R | PHE | 76 | H |
| VAL | 503 | R | PHE | 78 | H |
| VAL | 503 | R | SER | 114 | L |
| GLY | 504 | R | PHE | 76 | H |
| GLY | 504 | R | THR | 71 | H |
| GLY | 504 | R | TYR | 112 | L |
| TYR | 505 | R | ASP | 120 | H |
| TYR | 505 | R | ASN | 50 | H |
| TYR | 505 | R | TYR | 51 | H |
| TYR | 505 | R | ASP | 123 | H |
| GLN | 506 | R | TYR | 52 | L |
| GLN | 506 | R | SER | 114 | L |

| | | | | | |
|---|---|---|---|---|---|
| GLN | 506 | R | TYR | 112 | L |
| TYR | 508 | R | PHE | 76 | H |

**Table S5.** The list of the binding epitope residues and BD55-4637 Ab residues forming interfacial contacts

| RBD Residue | RBD Residue Number | RBD Chain | Antibody residue | Antibody residue number | Antibody chain |
|---|---|---|---|---|---|
| ALA | 372 | R | LYS | 31 | A |
| PRO | 373 | R | LYS | 31 | A |
| PHE | 374 | R | ASN | 32 | A |
| PHE | 374 | R | LYS | 30 | A |
| PHE | 374 | R | LYS | 31 | A |
| PHE | 374 | R | SER | 28 | A |
| PHE | 375 | R | ASN | 32 | A |
| PHE | 375 | R | LYS | 31 | A |
| THR | 376 | R | ASP | 102 | A |
| THR | 376 | R | LEU | 103 | A |
| THR | 376 | R | LYS | 31 | A |
| THR | 376 | R | LEU | 107 | A |
| THR | 376 | R | ASN | 32 | A |
| THR | 376 | R | GLY | 33 | A |
| PHE | 377 | R | LYS | 31 | A |
| PHE | 377 | R | LEU | 107 | A |
| LYS | 378 | R | SER | 105 | A |

| | | | | | |
|---|---|---|---|---|---|
| LYS | 378 | R | LEU | 107 | A |
| LYS | 378 | R | ASP | 106 | A |
| ARG | 403 | R | ARG | 31 | B |
| GLY | 404 | R | VAL | 109 | A |
| ASP | 405 | R | THR | 33 | B |
| ASP | 405 | R | VAL | 109 | A |
| ASP | 405 | R | HIS | 51 | B |
| VAL | 407 | R | ILE | 108 | A |
| VAL | 407 | R | VAL | 109 | A |
| VAL | 407 | R | LEU | 107 | A |
| ARG | 408 | R | ILE | 108 | A |
| ARG | 408 | R | LEU | 107 | A |
| ARG | 408 | R | ASP | 106 | A |
| GLN | 414 | R | ASP | 106 | A |
| VAL | 433 | R | LEU | 107 | A |
| ALA | 435 | R | ASN | 32 | A |
| ALA | 435 | R | LEU | 107 | A |
| TRP | 436 | R | ASN | 32 | A |
| ASN | 437 | R | ASN | 32 | A |
| ASN | 437 | R | GLY | 33 | A |
| ASN | 437 | R | ASP | 102 | A |
| ASN | 437 | R | TRP | 55 | A |
| ASN | 439 | R | TYR | 54 | A |

| | | | | | |
|---|---|---|---|---|---|
| ASN | 439 | R | SER | 58 | A |
| ASN | 439 | R | ASP | 56 | A |
| ASN | 439 | R | ARG | 60 | A |
| LYS | 440 | R | ASP | 56 | A |
| LYS | 440 | R | SER | 58 | A |
| SER | 496 | R | ARG | 31 | B |
| ARG | 498 | R | ARG | 31 | B |
| ARG | 498 | R | ASP | 94 | B |
| PRO | 499 | R | ARG | 60 | A |
| THR | 500 | R | ASP | 93 | B |
| THR | 500 | R | SER | 97 | B |
| THR | 500 | R | SER | 95 | B |
| THR | 500 | R | ARG | 60 | A |
| THR | 500 | R | TRP | 92 | B |
| THR | 500 | R | ASP | 94 | B |
| TYR | 501 | R | TRP | 92 | B |
| TYR | 501 | R | ARG | 31 | B |
| TYR | 501 | R | ARG | 60 | A |
| TYR | 501 | R | ASN | 32 | B |
| TYR | 501 | R | ASP | 94 | B |
| GLY | 502 | R | ASN | 32 | B |
| GLY | 502 | R | TRP | 92 | B |
| GLY | 502 | R | ASP | 94 | B |

| | | | | | |
|---|---|---|---|---|---|
| GLY | 502 | R | ASP | 111 | A |
| VAL | 503 | R | ASP | 102 | A |
| VAL | 503 | R | VAL | 109 | A |
| VAL | 503 | R | TRP | 92 | B |
| VAL | 503 | R | ASP | 111 | A |
| VAL | 503 | R | PRO | 101 | A |
| VAL | 503 | R | ASN | 32 | B |
| VAL | 503 | R | TYR | 54 | A |
| GLY | 504 | R | ASN | 32 | B |
| GLY | 504 | R | VAL | 109 | A |
| GLY | 504 | R | ASP | 111 | A |
| HIS | 505 | R | ASP | 94 | B |
| HIS | 505 | R | ARG | 31 | B |
| HIS | 505 | R | ASN | 32 | B |
| GLN | 506 | R | ARG | 60 | A |
| GLN | 506 | R | TYR | 54 | A |
| GLN | 506 | R | TRP | 55 | A |
| TYR | 508 | R | GLY | 33 | A |

**Table S6.** The list of the binding epitope residues and SA55 Ab residues forming interfacial contacts

| RBD Residue | RBD Residue Number | RBD Chain | Antibody residue | Antibody residue number | Antibody chain |
|---|---|---|---|---|---|
| PRO | 373 | R | LEU | 94 | B |
| PHE | 374 | R | THR | 57 | A |
| PHE | 374 | R | PHE | 55 | A |
| THR | 376 | R | PHE | 55 | A |
| ARG | 403 | R | PRO | 105 | A |
| ARG | 403 | R | ASN | 106 | A |
| GLY | 404 | R | PHE | 55 | A |
| GLY | 404 | R | LEU | 54 | A |
| GLY | 404 | R | ARG | 30 | A |
| ASP | 405 | R | LEU | 54 | A |
| ASP | 405 | R | SER | 31 | A |
| ASP | 405 | R | THR | 28 | A |
| ASP | 405 | R | ARG | 30 | A |
| GLU | 406 | R | ARG | 30 | A |
| VAL | 407 | R | ARG | 30 | A |
| VAL | 407 | R | PHE | 55 | A |
| VAL | 407 | R | LEU | 54 | A |

| | | | | | |
|---|---|---|---|---|---|
| ARG | 408 | R | ARG | 30 | A |
| ASN | 437 | R | ASP | 93 | B |
| ASN | 439 | R | TYR | 91 | B |
| ASN | 439 | R | ASP | 93 | B |
| LYS | 440 | R | ASP | 93 | B |
| VAL | 445 | R | HIS | 53 | B |
| TYR | 495 | R | PRO | 105 | A |
| SER | 496 | R | PRO | 105 | A |
| ARG | 498 | R | PHE | 112 | A |
| ARG | 498 | R | TYR | 49 | B |
| PRO | 499 | R | PHE | 100 | A |
| PRO | 499 | R | PRO | 101 | A |
| PRO | 499 | R | ASP | 50 | B |
| PRO | 499 | R | TYR | 91 | B |
| THR | 500 | R | GLY | 103 | A |
| THR | 500 | R | PHE | 112 | A |
| THR | 500 | R | PHE | 100 | A |
| THR | 500 | R | TYR | 49 | B |
| THR | 500 | R | ASP | 104 | A |
| THR | 500 | R | PRO | 101 | A |
| THR | 500 | R | ASN | 102 | A |
| THR | 500 | R | ASP | 50 | B |
| TYR | 501 | R | PRO | 105 | A |

| | | | | | |
|---|---|---|---|---|---|
| TYR | 501 | R | GLY | 103 | A |
| TYR | 501 | R | PHE | 112 | A |
| TYR | 501 | R | ASN | 102 | A |
| TYR | 501 | R | ASP | 104 | A |
| TYR | 501 | R | PRO | 101 | A |
| GLY | 502 | R | ASP | 104 | A |
| GLY | 502 | R | PRO | 101 | A |
| GLY | 502 | R | SER | 31 | A |
| GLY | 502 | R | GLY | 103 | A |
| GLY | 502 | R | HIS | 32 | A |
| GLY | 502 | R | ASN | 102 | A |
| VAL | 503 | R | PRO | 95 | B |
| VAL | 503 | R | PHE | 55 | A |
| VAL | 503 | R | VAL | 33 | A |
| VAL | 503 | R | ASN | 102 | A |
| VAL | 503 | R | LEU | 54 | A |
| VAL | 503 | R | HIS | 32 | A |
| VAL | 503 | R | PRO | 101 | A |
| VAL | 503 | R | ILE | 52 | A |
| VAL | 503 | R | SER | 31 | A |
| GLY | 504 | R | LEU | 54 | A |
| GLY | 504 | R | HIS | 32 | A |
| GLY | 504 | R | SER | 31 | A |

| | | | | | |
|---|---|---|---|---|---|
| GLY | 504 | R | ARG | 30 | A |
| HIS | 505 | R | PRO | 105 | A |
| HIS | 505 | R | HIS | 32 | A |
| HIS | 505 | R | ASP | 104 | A |
| HIS | 505 | R | SER | 31 | A |
| HIS | 505 | R | GLY | 103 | A |
| GLN | 506 | R | PRO | 101 | A |
| GLN | 506 | R | TYR | 91 | B |
| GLN | 506 | R | ASP | 93 | B |
| TYR | 508 | R | LEU | 54 | A |
| TYR | 508 | R | PHE | 55 | A |
